# Characterisation and zoonotic risk of tick viruses in public datasets

**DOI:** 10.1101/2022.12.05.518373

**Authors:** Yuting Lin, David J Pascall

## Abstract

Tick-borne viruses remain a substantial zoonotic risk worldwide, so knowledge of the diversity of tick viruses has potential health consequences. Despite their importance, large amounts of sequences in public datasets from tick meta-genomic and –transcriptomic projects remain unannotated, sequence data that could contain undocumented viruses. Through data mining and bioinformatic analyses of more than 37,800 public meta-genomic and -transcriptomic datasets, we found 83 unannotated contigs exhibiting high identity with known tick viruses. These putative viral contigs were classified into three RNA viral families (*Alphatetraviridae*, *Orthomyxoviridae*, *Chuviridae*) and one DNA viral family (*Asfaviridae*). After manual checking of quality and dissimilarity toward other sequences in the dataset, these 83 contigs were reduced to five putative novel Alphatetra-like viral contigs, four putative novel Orthomyxo-like viral contigs, and one Chu-like viral contig which clustered with known tick-borne viruses, forming a separate clade within the viral families. We further attempted to assess which previously known tick viruses likely represent zoonotic risks and thus deserve further investigation. We ranked the human infection potential of 136 known tick-borne viruses using a genome composition-based machine learning model. We found five high-risk tick-borne viruses (Langat virus, Lonestar tick chuvirus 1, Grotenhout virus, Taggert virus, and Johnston Atoll virus) that have not been known to infect human and two viral families (*Nairoviridae* and *Phenuiviridae*) that contain a large proportion of potential zoonotic tick-borne viruses. This adds to the knowledge of tick virus diversity and highlights the importance of surveillance of newly emerging tick-borne diseases.

**Importance:** Ticks are important hosts of pathogens. Despite this, numerous tick-borne viruses are still unknown or poorly characterised. To overcome this, we re-examined currently known tick-borne viruses and identified putative novel viruses associated with ticks in public datasets. Using genome-based machine learning approach, we predicted five high-risk tick-borne viruses that have not yet been reported to cause human infections. Additionally, we highlighted two viral families, *Nairoviridae* and *Phenuiviridae*, which are potential public health threats. Our analysis also revealed 10 putative novel RNA viral contigs clustered with known tick-borne viruses. Our study highlights the importance of monitoring ticks and the viruses they carry in endemic areas to prevent and control zoonotic infectious disease outbreaks. To achieve this, we advocate for a multidisciplinary approach within a One Health and EcoHealth framework that considers the relationship between zoonotic disease outbreaks and their hosts, humans, and the environment.

## Introduction

The role of ticks in the transmission of viruses has been known for over 100 years (1) with ticks being second only to mosquitoes as vectors of pathogens to humans and the primary vector of pathogens to livestock, wildlife, and companion animals (2). Nonetheless, new tick viruses are still being detected (3). Two recent examples are Yezo virus identified in Japan (4) and Songling virus identified in China (5), which have both been associated with acute human febrile disease.

Prior to the advent of molecular biology, tick-borne viruses were identified through microscopy, isolation in culture or serology (6–11). Over the last two decades, meta-genomics and – transcriptomics tools have revolutionised virus discovery. These approaches employ high-throughput sequencing technologies to capture total viral communities in a relatively unbiased manner (12,13) and have facilitated studies of the tick virome that resulted in the discovery and characterisation of hundreds of new tick-borne viruses globally (14–23). While this has led to dramatic increases in the understanding of tick virus communities, being able to evaluate the human infection potential of the discovered viruses would additionally allow the leveraging of this data to provide public health benefits.

Models of zoonotic risk assessment often rely on the existence of zoonotic viruses in the same viral family or evolutionary relatedness to known zoonoses (24). These models rely on phenotypic information commonly unavailable for novel viruses or of insufficient resolution to estimate different levels of zoonotic potential for closely related viruses (25–27). More recent models based on the genomic features of viruses which do not strictly rely on phylogeny or taxonomy avoid this issue. Such models use genomic features that may individually contain weak signals but can be combined and exploited via machine learning algorithms (28–31). These models, built from genomic data alone, have particular strength in evaluating newly discovered viruses that lack other information sources, but are limited by the need for full viral genomes for accurate assessment.

In this study, we aimed to detect novel viruses that may have been overlooked by previous tick meta-genomic and –transcriptomic studies, and assess the zoonotic risk posed by tick virus diversity as it is currently understood. Thus, we searched publicly available tick sequence data for signs of viruses misidentified as tick sequences and evaluated the zoonotic risk of these and other tick viruses to find those which deserve further investigation.

## Materials and methods

### Datasets

We collected tick virus data from reports published prior to December 2021 and constructed a tick virus dataset that contained 261 tick-borne viruses from 23 viral families (34 genera) and 12 putative tick-borne viruses sampled from tick pools. Tick-borne viruses with unknown tick vectors (N = 13) or sequences unavailable (N = 13) were excluded from analyses. This resulted in 235 viruses in the final tick virus database, of which 3 of them (Amblyomma dissimile mivirus, Nuomin virus, and Granville quaranjavirus) were added later and not included in the database for virus detection. Among the 235 tick-borne viruses, more than 92% of viruses were RNA viruses, while few other viral sequences were identified (2.2% DNA viruses and 5.6% viruses with unknown genome composition). Among the RNA viruses, most sequences were negative-sense (-) single-stranded RNA (ssRNA) viruses (63.0%), followed by positive-sense (+) single-stranded RNA (ssRNA) (20.8%), double-stranded RNA (dsRNA) (8.5%), and unclassified RNA (5.6%) viruses, depending on version 2020 of the International Committee on Taxonomy of Viruses (ICTV) master species list (https://talk.ictvonline.org/files/master-species-lists/). We labelled each virus as being capable of infecting humans or not known to infect humans using published reports as ground truth. In all cases, only viruses detected in humans by either PCR or sequencing were considered to have proven ability to infect humans. See the summary table of currently known tick-borne viruses in **Appendix Table 1**.

### Viral identification and annotation

We used representative genomic sequences of the 232 tick-borne viruses as our reference database, with genome segments of segmented viruses concatenated into one. Using the ‘blastn_vdb’ (corresponding to nucleotide BLAST) executable in the BLAST+ v2.13.0 (32–34), we searched the Sequence Read Archive (SRA), Transcriptome Shotgun Assembly (TSA), and Whole Genome Shotgun (WGS) datasets that derived from ticks and subordinate taxa directly with an expectation cut-off of 0.05. We also performed a ‘tBLASTx’ search for the TSA database to find distant relationships between nucleotide sequences; SRA and WGS datasets were not suitable for ‘tBLASTx’ search because it was computationally prohibitive. This resulted in 1328 TSA, 19,990 SRA, and 1861 WGS sequences after removing duplicates. The GenBank accession numbers of the candidate sequences are provided at https://github.com/ytlin2021/TickVirus.git.

Candidate sequences were filtered according to the pipeline modified from Webster et al. (35). The thresholds used in each step can be seen in **Figure 1**. To confirm final RNA candidates, we checked the presence of RNA-dependent RNA polymerase (RdRp), an essential protein encoded in the genomes of all replicating RNA viruses without a DNA stage except deltaviruses, and not present in the genome of the eukaryotic or prokaryotic cell. We adopted a combined analysis approach to predict RdRps by using a general protein function prediction software package InterProScan v5.55-88.0 (36) and a new open bioinformatics toolkit - RdRp-scan - that allowed the detection of divergent viral RdRp sequences (37). Candidate sequences less than 600 nucleotides (nt) in length were discarded. The translated candidate sequences were searched according to the RdRp-scan workflow (37) and were also predicted using the InterProScan search function. The conserved RdRp motifs were annotated manually based on the RdRp motif database in RdRp-scan, and final RNA viral candidates were confirmed. The workflow of detecting viral RdRp sequences and the thresholds used in each step can be seen in **Appendix Figure 1**.

**Figure 1.**
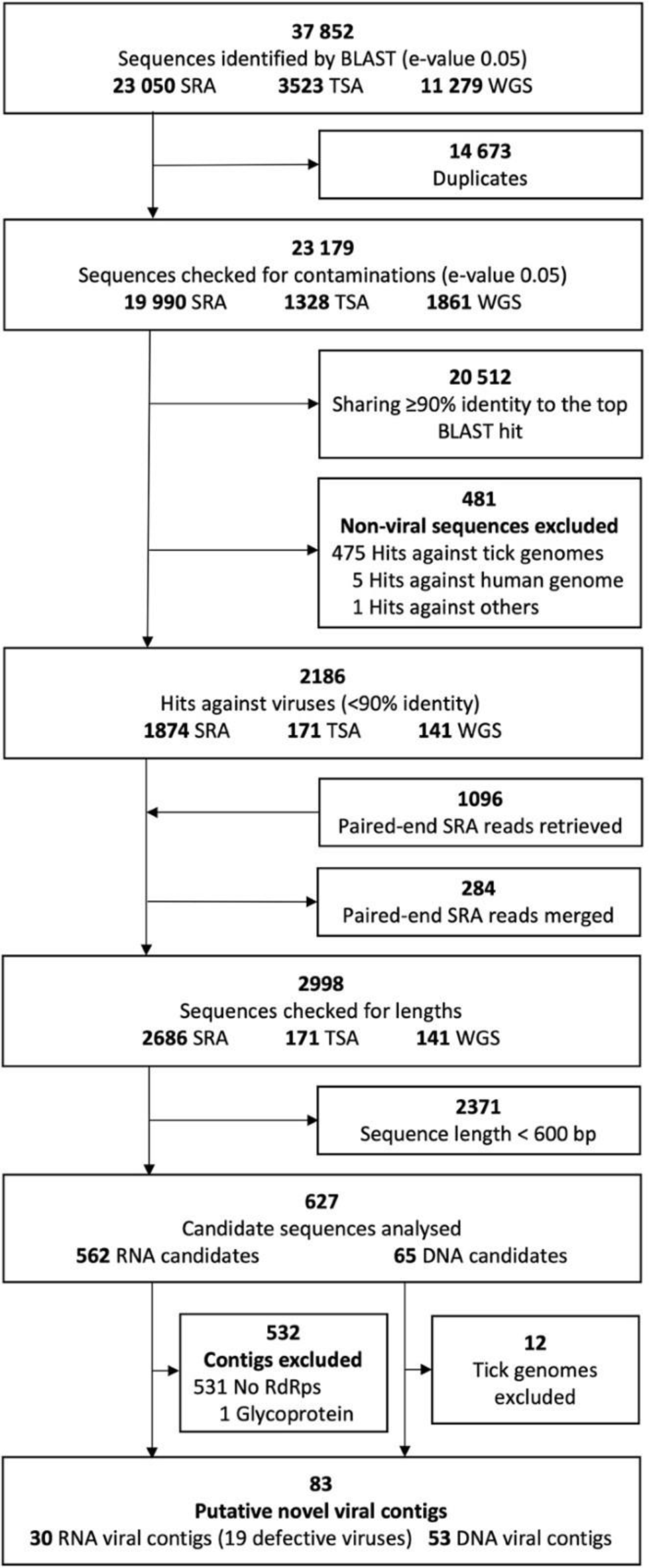
The workflow of identification of novel tick-associated viral contigs. Through BLAST search and curation, 53 DNA sequences (identified as African swine fever like elements) and 30 RNA sequences (putative viruses in the family *Alphatetraviridae, Orthomyxoviridae* and *Chuviridae*) from a collection of tick meta-genomic and –transcriptomic datasets (3 March 2022) were identified as putative novel viral contigs. 10 putative RNA viral contigs were further analysed after filtering. TSA = Transcriptome Shotgun Assembly, SRA = Sequence Read Archive, WGS = Whole Genome Shotgun, E-value = Expectation value.

Protein functions of putative novel viral contigs were annotated using InterProScan and ‘BLASTx’. Open Reading Frames (ORFs) were predicted using ORF Finder tool in NCBI (38) with a 150 nt length cut-off. InterProScan searched for protein function annotations in its available databases, including the Gene Ontology (39,40) and Pfam (41,42) databases. At the same time, ‘BLASTx’ were used to search against the non-redundant protein database with an expectation cut-off of 1e-5 following Charon et al. (37).

### Phylogenetics

We searched for close and distant relatives of the identified novel viral contigs in viruses (NCBI taxa ID: 10239) non-redundant nucleotide and protein databases by performing ‘BLASTn’ and ‘BLASTx’ separately with an expectation cut-off of 0.005. For each contig with BLAST hits, the phylogenetic hierarchy of the best hits (top 10 hits from ‘BLASTx’) was traversed upward to identify the lowest taxonomic classification displaying a 75% majority taxonomic agreement following Webster et al. (35). If the best hits were unclassified viruses, the upper order was identified, and the protein sequences were aligned in each family under the order to determine the classification. This allowed putative novel viral contigs in the study to be classified into three RNA viral families (*Alphatetraviridae*, *Orthomyxoviridae*, *Chuviridae*) and one DNA viral family (*Asfaviridae*). All virus species in the same family, as well as unassigned viruses, were aligned and used to construct phylogenetic trees.

Phylogeny of RNA viruses was inferred on the basis of RNA-dependent RNA polymerase protein sequences. We aligned viral RdRp protein sequences using the ClustalW algorithm (43) embedded in the GUIDANCE2 server v2.02 (44). GUIDANCE2 server removed any sequences with a score of less than 0.6 and columns with a score of less than 0.93, aiming to leave sufficient information in alignments while allowing large taxonomic groups of viruses to be compared.

We performed tree estimation for all trees in BEAST2, averaging over protein models using the OBAMA module (45,46). A lognormal relaxed clock model was applied (56–58), and a Yule tree prior was used (47,48). As convergence repeatedly failed under standard MCMC for the *Chuviridae* and *Orthomyxoviridae* trees, models were fit using BEAST2’s Metropolis-coupled MCMC routines with 4 chains and a target cross-chain move acceptance probability of 0.029 for the *Alphatetraviridae* and *Chuviridae* trees, and a fixed heating parameter of 0.3 for the *Orthomyxoviridae* tree (49,50). The Yule process birth rate was given a vague Exponential prior with mean 1000, all other parameters took default prior values. Trees were run for 100,000,000 iterations with 50,000,000 discarded as burn-in, and the posterior then being thinned to 1000 trees. A second run of each tree was used to assess convergence with the R package RWTY (51).

### Zoonotic risk analyses

The zoonotic risk (precisely, probability of human infection given biologically relevant exposure) for 136 tick-borne viruses with complete genome sequences were evaluated using the best performing model from Mollentze et al. (31), referred to here as genome composition-based (GCB), which was trained on a range of viral genome features and human similarity features. A representative genome was selected for each virus, giving preference to sequences from NCBI Reference Sequence Database (RefSeq) (www.ncbi.nlm.nih.gov/refseq/) wherever possible. RefSeq sequences that had annotation issues or were not judged to be representative of the naturally circulating virus were replaced with alternative genomes. Following Mollentze et al. (31), the predicted probabilities from the GCB model were used to categorise viruses into four priority categories based on the overlap of confidence intervals (CIs) the value of 0.293 (low: entire 95% CI of predicted probability ≤ cut-off; medium: mean prediction ≤ cut-off, but CI crosses it; high: mean prediction > cut-off, but CI crosses it; very high: entire CI > cut-off). Since the current GCB model was trained for viruses with complete sequences, 99 tick viruses with only partial genomes available were excluded from the analyses. To characterise the zoonotic potential of the identified novel viral contigs, we did qualitative analyses by evaluating the published complete genomes of known viruses most similar to these incomplete viral sequences as determined by a nucleotide BLAST against GenBank.

## Results

### Identification of putative novel viruses associated with ticks

We examined contigs that showed nucleotide similarity (< 90% identity) to previously characterised tick virus sequences from an extensive collection of tick meta-genomic and –transcriptomic datasets (23,050 SRA, 3523 TSA, and 11,279 WGS sequences; collected on 3 March 2022). The pipeline of detecting novel tick-associated viral contigs can be found in **Figure 1**. For more information on the tick virus database and tick meta-genomic and –transcriptomic datasets, see Methods.

In total, we identified 83 putative novel viral contigs, including 53 double-stranded DNA viral contigs, 5 positive-sense and 25 negative-sense RNA viral contigs (**Appendix Table 2–4**). The 53 DNA viral contigs belong to the family *Asfaviridae*, with homology to African swine fever virus. These African swine fever virus-like contigs were found in *Ornithodoros porcinus* and *Ornithodoros moubata* ticks (**Appendix Table 4**), and all but two of these contigs derived from a study that had previously reported the integration of African swine fever like elements into the genomes of these soft tick species (52), so they will not be discussed further. The 5 positive-sense RNA viral contigs belong to the family *Alphatetraviridae*. Of the negative-sense RNA viral contigs, 5 belong to the family *Orthomyxoviridae*, and 20 belong to the family *Chuviridae*, including 19 chuviruses believed to be replication defective from tick SRA projects. After filtering duplicate sequences (GenBank IDs:

HACP01027211.1 and HACW01024387.1), we obtained 10 putative RNA viral contigs ranging from 728 bp to 5496 bp (**Table 1**). The putative novel viruses found in this study were named after their host order and related viral family, followed by a number (e.g., Alphatetra-like tick virus 1). All sequences identified in the present study with can be found on GenBank (see accession numbers in **Table 1** and **Appendix Table 2–4**).

**Table 1.**
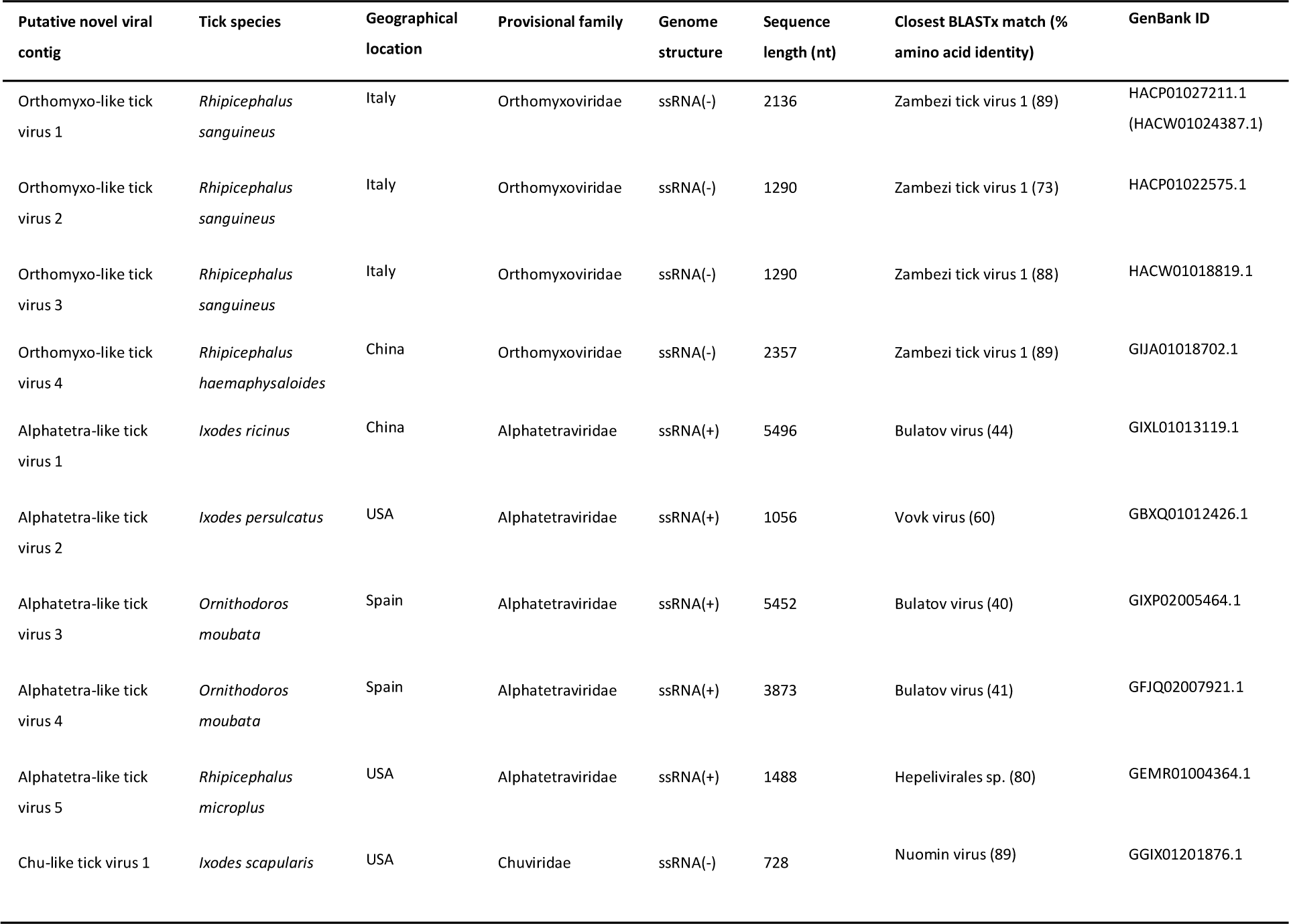
The putative novel RNA viral contigs found after bioinformatic checking.

#### (i) Orthomyxoviridae-like contigs

The contigs we found contained homologs of Influenza RdRp subunit PB1, the most conserved of the orthomyxovirus genes (**Appendix Figure 2**). Phylogenetic analysis revealed that the putative novel Orthomyxo-like viral contigs fell into the genus *Quaranjavirus* and formed a separate clade with other tick-borne quaranjaviruses (**Appendix Figure 3**; **Table 1**). Within PB1, the contigs were phylogenetically closely related to Zambezi tick virus 1 (ZaTV-1) – a highly divergent virus identified in Rhipicephalus ticks from Mozambique (53) and shared approximately 77% amino-acids identity (**Appendix Figure 3**; **Table 1**). Quaranjaviruses are enveloped negative-sense single-stranded segmented RNA viruses, generally with multiple segments (54) which have been recorded in argasid and ixodid ticks, other arachnids, insects and vertebrates (55,56). In the present study, we identified 4 putative novel Orthomyxo-like tick viruses from ixodid tick metagenomic projects, 3 of which derived from sequence sampled from *Rhipicephalus sanguineus* in Italy, and one from *Rhipicephalus haemaphysaloides* in China (**Table 1**).

We note that four of the six the viral species that have complete genomes in this clade were either predicted to have very high (N = 1) or high (N = 3) zoonotic potential based on the GCB model (**Table 2**; the GenBank ID can be found in **Appendix Table 5**), with Quaranfil virus (QRFV), one of the viruses rated to have high zoonotic potential, being confirmed to lead to infections in humans (54). Taken together, these results suggest that the novel Orthomyxo-like viral contigs identified in the current study may represent a potential risk of human infection given exposure (which is possible, as *R. haemaphysaloides* is known to parasitise humans).

**Table 2.**
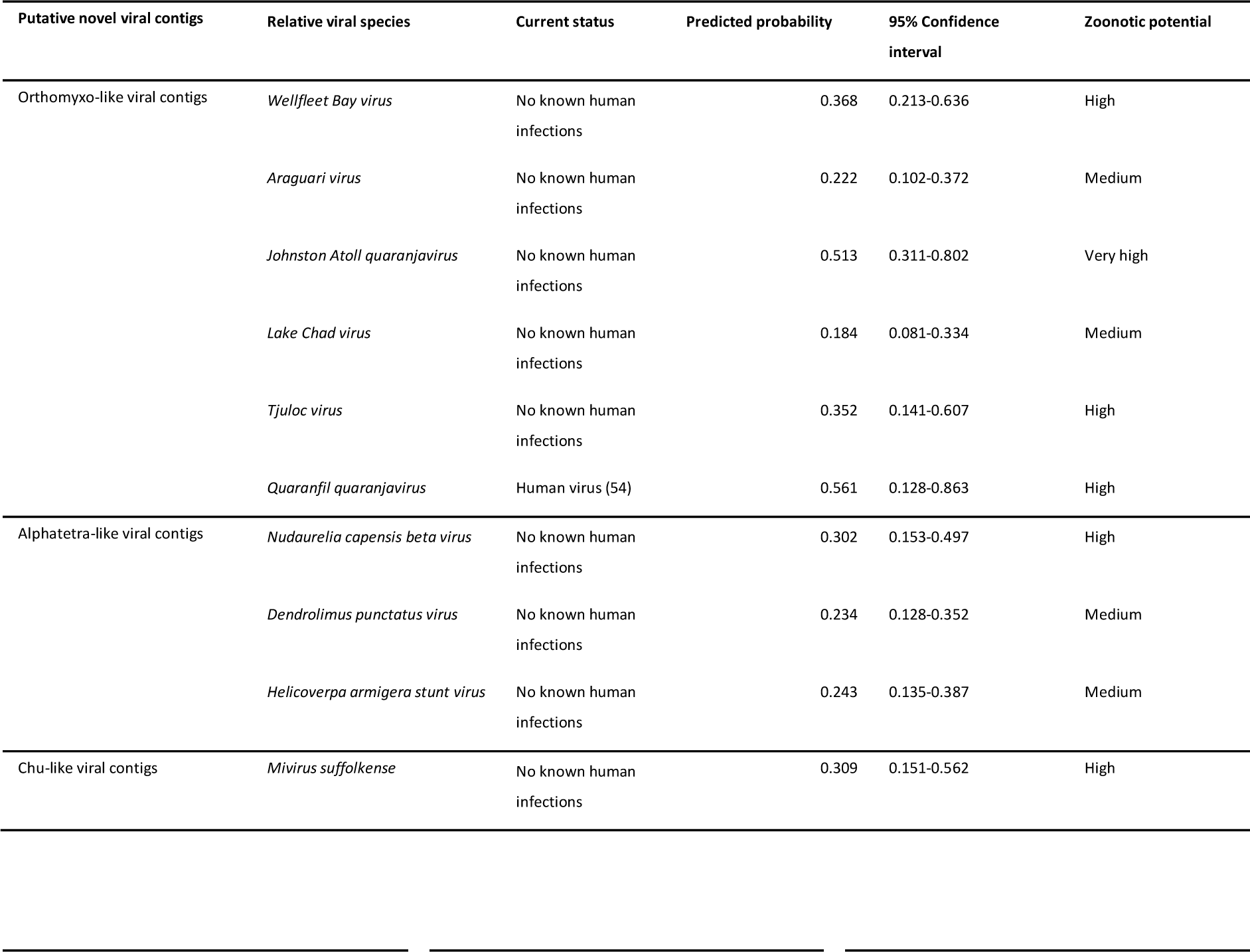
Zoonotic risks of virus species closely related to the putative novel Alphatetra-like viral contigs. Zoonotic risks of sequences in GenBank format were predicted using the GCB model.

#### (ii) Alphatetraviridae-like contigs

Members of the *Alphatetraviridae* are non-enveloped positive-sense single-stranded RNA viruses (57). The identified Alphatetra-like contigs ranged from 1488 nt to 5496 nt, detected via the presence of a putative RdRp 2 motif; some of the longer sequences also encoded putative nucleoside triphosphate hydrolase, viral RNA helicase, and viral methyltransferase three motifs (**Figure 2B**). Viruses in the family *Alphatetraviridae* had been previously believed to infect predominantly moths and butterflies (57), but the recent discovery of several Alphatetra-like viruses in ticks (23,58) implies that this may have been due to sampling biases. Consistent with this, we found the presence of potential Alphatetra-like viruses in four ixodid tick species (*Ixodes ricinus*, *Ixodes persulcatus*, *Ornithodoros moubata* and *Rhipicephalus microplus*) sampled in three geographically diverse locations (China, Spain, and USA) (**Table 1**). Phylogenetic analysis suggested that the novel Alphatetra-like tick viruses formed a separate clade, being placed with the tick-borne viruses recently identified in ixodid ticks from Antarctica (Vovk virus and Bulatov virus from the seabird tick*, Ixodes uriae*) (58) and China (Hubei tick hepe-like virus found in *R. microplus* and *Haemaphysalis longicornis*) (23), and Heilongjiang sediment betatetravirus of unknown host, but originally sampled from river sediment in China (59) (**Figure 3B**; **Table 1**).

**Figure 2.**
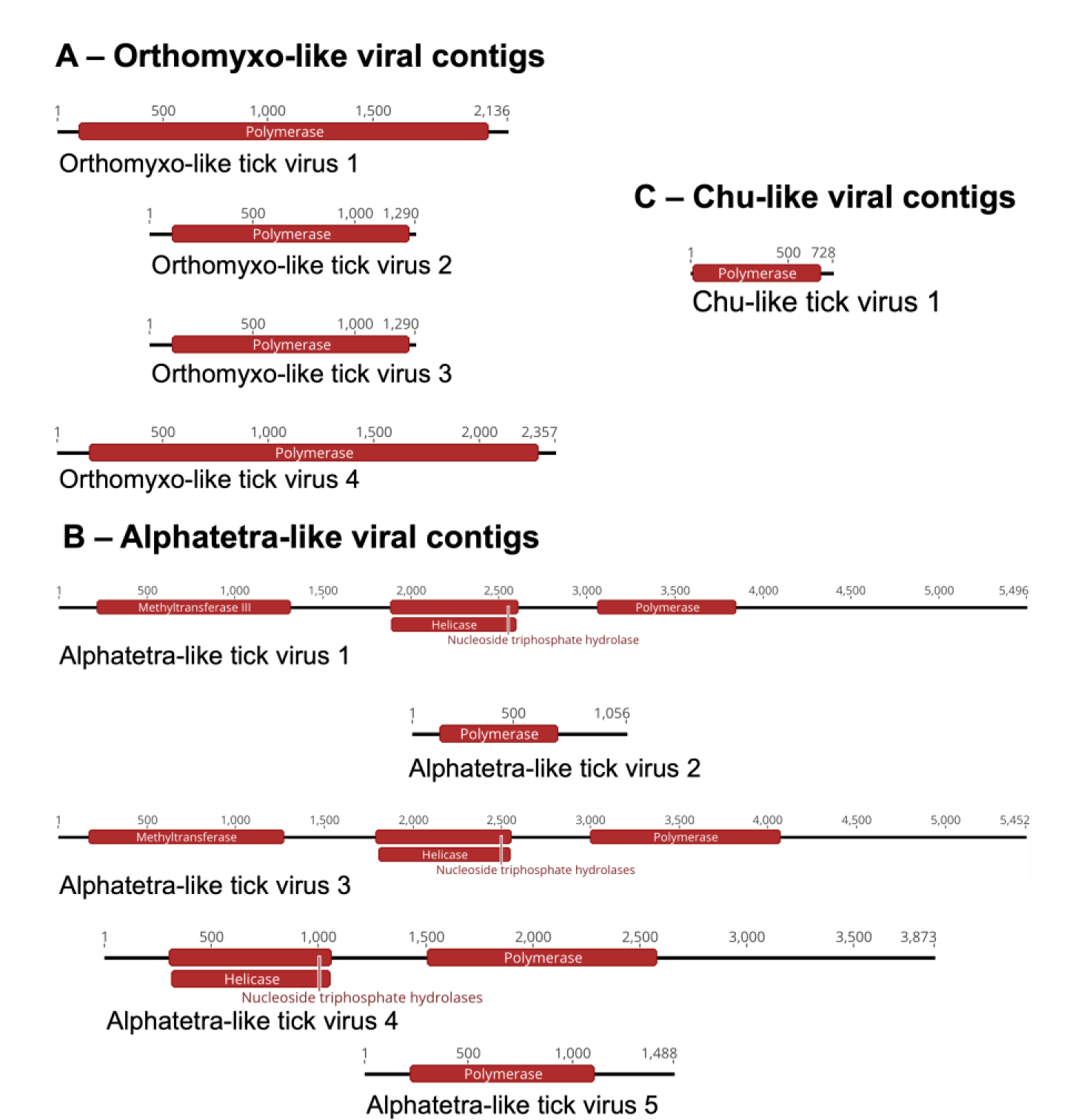
Genome structures of novel viral contigs.

**Figure 3.**
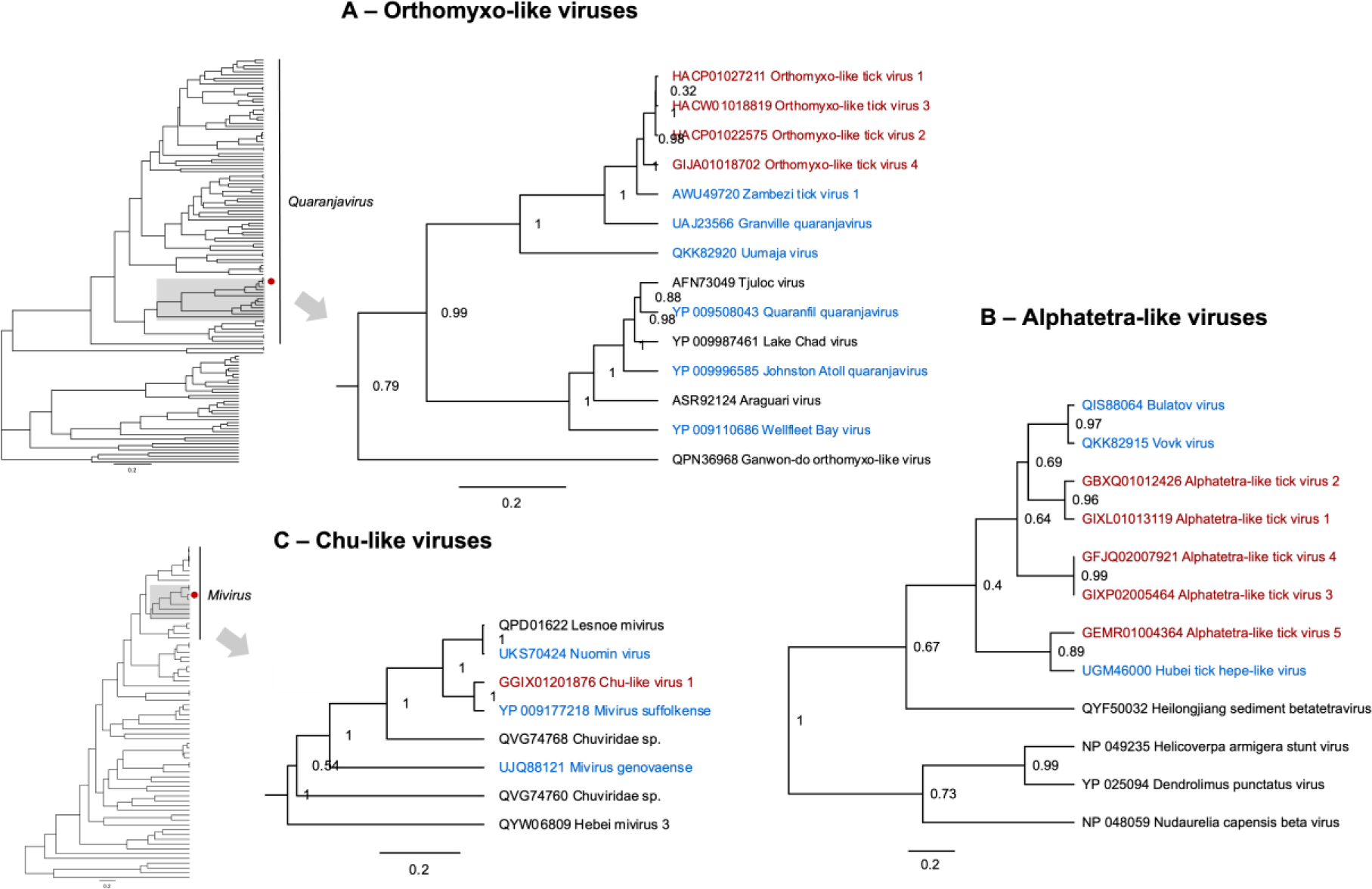
Phylogenetic relationship of the novel viral contigs. Maximum clade credibility trees were inferred from the Bayesian posterior sample. Posterior clade probabilities are shown at the nodes, and the scale is given in amino-acid substitutions per site. Putative novel viral contigs found in the present study are shown in red, and previously known tick-borne viruses are shown in blue. GenBank ID can be found at **Appendix Figure 2–4**. Alignments using ClustalW and maximum clade credibility trees are provided in at https://github.com/ytlin2021/TickVirus.git.

No members of the *Alphatetraviridae* are known to infect humans, and, unfortunately, only partial genomes were available for the previously identified viruses in the proposed new clade (Bulatov virus, Vovk virus and Heilongjiang sediment betatetravirus), thus they were inappropriate for zoonotic risk analysis based on the GCB model. The closest related viruses with full genomes were all isolated from moths. Of these viruses Nudaurelia capensis beta virus was predicted to have high zoonotic potential; Dendrolimus punctatus virus and Helicoverpa armigera stunt virus were predicted to have medium zoonotic potential (**Table 2**; the GenBank IDs can be found in **Appendix Table 5**). This result illustrates a deficiency of the currently available zoonotic risk assessment tools for viruses discovered through meta-genomic or –transcriptomic methods. Zoonotic risk estimation using genome information generally requires complete viral genomes, but viruses found through these methods are often incomplete or exhibit uncertainty in segment linkage for segmented viruses, which impacts the ability to make fast decisions on virus research and surveillance at the earliest stage of virus discovery. Training existing models on features from partial viral genomes provides a potential avenue for future work to help alleviate this issue.

#### (iii) Chuviridae-like contigs

We identified 20 Chu-like viral contigs from *Ixodes scapularis* and *Hyalomma asiaticum* sampled in the US and China (**Table 1**; **Appendix Table 3**). Members of the *Chuviridae* are negative-sense single-stranded RNA viruses, which display a wide variety of genome organisations, including unsegmented, bi-segmented, and circular forms (15). Only the contig labelled Chu-like tick virus 1 is consistent with a live virus, as all other contigs showed signs of mutational degradation with many stop codons present in the region with homology to the RdRp. All but one of the contigs exhibiting evidence of degradation were generated from a laboratory population of *Hy. asiaticum*, maintained under controlled conditions (60). As such, we speculate that they likely result from the presence of an integrated Chu-like virus in the *Hy. asiaticum* genome present in this laboratory population, and, likely, more broadly.

The viral sequence of Chu-like tick virus 1 was 728 nt in length, comprising one putative ORF that encodes a 240-amino acid protein containing the potential RdRp domain, nucleoside triphosphate hydrolase, viral RNA helicase, and viral methyltransferase 3 (**Figure 2C**). Chu-like tick virus 1 was clustered with other miviruses and was phylogenetically closely related to Suffolk virus, originally sampled in USA (20), with approximately 87% nucleotide identity (**Figure C**; **Table 1**). With regards to the human infection potential of Chu-like tick virus 1, it is notable that it showed a RdRp amino-acid identity of around 90% with Nuomin virus which has recently been associated with human febrile illness in China (61). The prediction results also classified Suffolk virus into the high zoonotic potential category (**Table 2**; the GenBank IDs can be found in **Appendix Table 5**). These results suggest that Chu-like tick virus 1 may pose a zoonotic risk. This may be especially relevant as the sequence was detected in *I. scapularis*, an important vector of other human pathogens, and so the potential for human exposure is high. However, given the small number of known species in the family *Chuviridae* when the GCB model was trained (31), caution must be applied, as the model might not be able to accurately predict the zoonotic potential viruses in this family. There would therefore seem to be a definite need for characterising viruses in the family *Chuviridae* in terms of their viability in human cell lines with these results feeding through to future zoonotic modelling efforts.

### Zoonotic ranking of known tick-borne viruses

We used the prediction framework illustrated by Mollentze and colleagues (31) to rank 136 known tick-borne viruses with complete genomes. The viruses here were ranked based on predicted mean probability of zoonotic potential and further converted into four zoonotic potential categories, describing the overlap of confidence intervals (CIs) with the 0.293 cut-off (see Methods). The zoonotic prediction results were given in **Appendix Table 6**. This subset dataset contained representatives from 13 viral families (29 genera) and unassigned viruses, including 21 viruses that were known to infect humans by our criteria.

In total, 5.1% of tick-borne viruses were identified as having very high zoonotic potential (N = 7), 39.7% having high zoonotic potential (N = 54), 44.1% having medium zoonotic potential (N = 60) and 11.0% having low zoonotic potential (N = 15; **Appendix Figure 4**). Among the 22 currently known human-infecting tick-borne viruses, 81.8% were correctly identified as having either very high (N = 2) or high zoonotic potential (N = 16), and the remaining human-associated viruses were classified as medium zoonotic potential (N = 4; **Appendix Figure 4**). Among the tick-borne viruses with unknown human infectivity that were sequenced from nonhuman animal or tick samples, 37.7% were predicted to have either very high (N = 5) or high zoonotic potential (N = 38; **Appendix Figure 4**), including viruses in the family *Chuviridae*, *Nyamiviridae* and *Parvoviridae* that currently do not contain tick-borne viruses known to infect humans. Langat virus, Lonestar tick chuvirus 1, Grotenhout virus, Taggert virus, and Johnston Atoll virus were predicted to have very high zoonotic risk, although they were not known to infect humans (**Figure 4B**). We therefore recommend these viruses be a high priority for further virological research, in order to assess whether greater surveillance of them in wild tick populations would be warranted.

**Figure 4.**
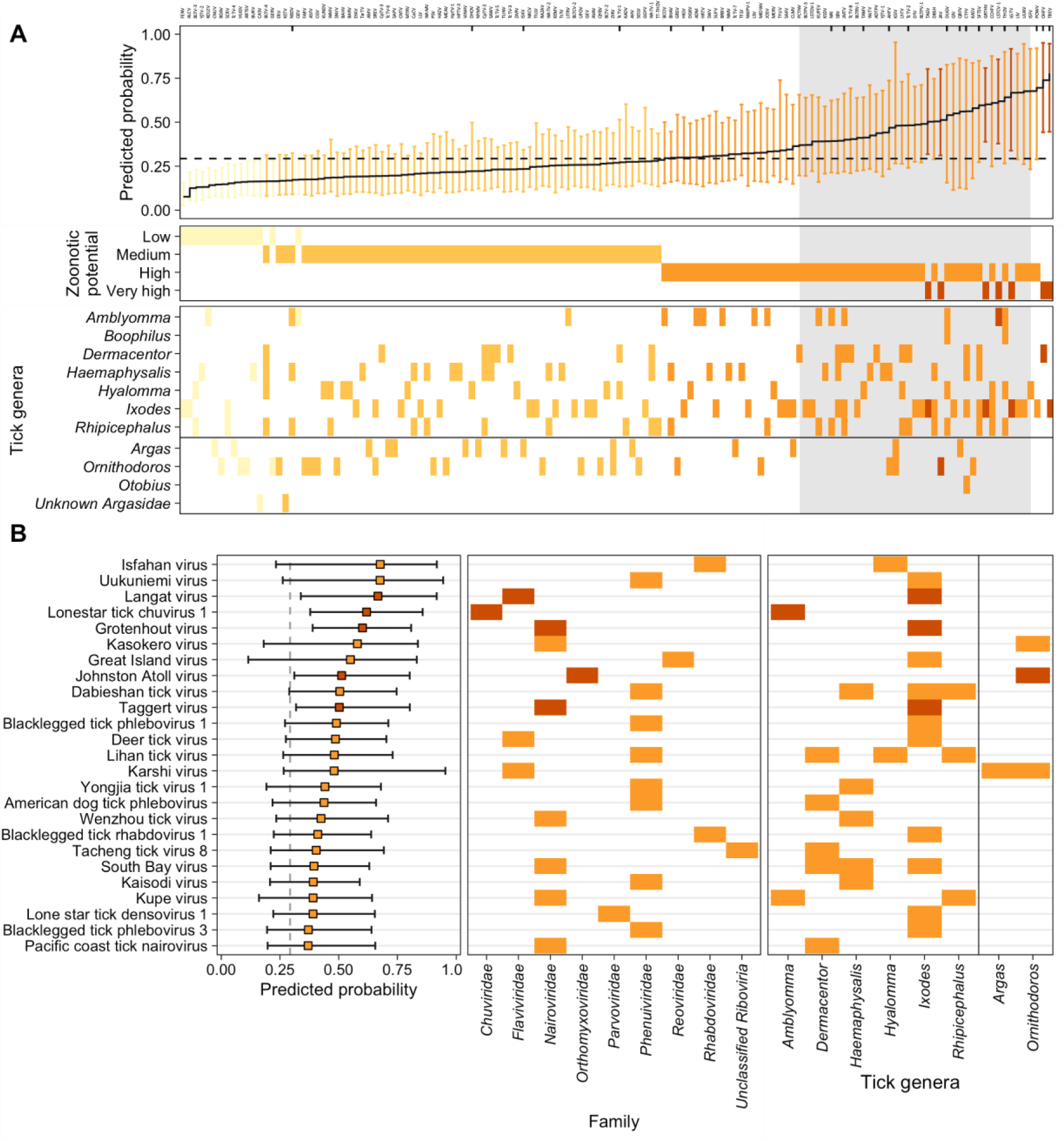
Predicted probability of human infection for tick viruses based on genome composition-based model. (**A**) Predicted probability of human infection for 136 tick viruses with complete genomes. Colour scale show the assigned zoonotic potential categories. Tick marks along the top edge of the first panel show the location of viruses known to infect humans, while a dashed line shows the cut-off 0.293 that balanced sensitivity and specificity according to Mollentze et al. (31). The top 25 viruses that have no known human infection (contained within the grey box) are illustrated in more detail in (**B**) Viruses that have no known human infection are shown in red. Points show the mean calibrated score, with lines indicating 95% confidence intervals. Figures were drawn in R v4.1.1. Zoonotic ranking results can be found in **Appendix Table 6**. Numeric data underlying this figure can be found at https://github.com/ytlin2021/TickVirus.git.

We noted that the high and very high zoonotic potential categories were dominated by *Nairoviridae* (31.0%) and *Phenuiviridae* (28.6%), consistent with the recent findings of emerging human-infecting tick viruses in these two viral families. For example, among members of the *Nairoviridae* family, two emerging orthonairoviruses called Songling virus and Yezo virus have been recently detected in *Hyalomma* spp. ticks and isolated in patients with the acute febrile disease (4,5). Hospitals in Inner Mongolia and Heilongjiang province, China also reported infections from a novel nairovirus named Beiji nairovirus which mainly circulates in the ixodid ticks *I. crenulatus* and *I. persulcatus* (62).

Similarly, among the *Phenuiviridae*, a novel phlebovirus named Tacheng tick virus 2 (TcTV-2) was first sampled from *D. marginatus* ticks in China (15) and later identified in various countries and tick species, primarily circulating in *D. marginatus* and *H. marginatum* ticks (63–65). The risk for human infection from TcTV-2 was not known until 2021 when Dong and colleagues reported on TcTV-2 infection in a patient in China (66). As recently emerged viruses, the geographic ranges of these viruses are poorly understood, and only partial genomes are available. The wide distribution of these tick species suggests that the geographic limits of the emerging tick-borne viruses in the family *Nairoviridae* and *Phenuiviridae* may be larger than presently assumed.

## Discussion

Here we reanalysed existing tick metagenomic and meta-transcriptomic studies in public datasets to identify potential novel tick-borne viruses, using a tick virus database containing all known tick-borne viruses with genetic sequence available. We then employed the genome composition-based machine learning model of Mollentze et al. (31) to evaluate the zoonotic risk of all published tick viruses with complete genomes.

### Novel virus discovery

The presence of undetected viruses in pre-existing meta-genomic and –transcriptomic data illustrate a valuable underutilised data source. A combination of factors, ranging from studies being focused on specific taxa, limitations of the bioinformatic tools being used and incompleteness of reference datasets used within these tools, result the generation of large amounts of unannotated or incorrectly annotated sequence. This provides an opportunity for small studies such as this one, or much larger studies considering wider ranges of or more extensively studied taxa (e.g. (16,35,67)), reanalysing this historical data to detect new viruses (or, equally, other organisms of interest). As a pedagogical aside, as the resource requirements for these studies are low, the scope of the projects limited, the outputs of general scientific interest, and the skills required of particular value to employers, these studies make excellent short projects for late-stage undergraduate or early-stage postgraduate students on ecology or ecology-adjacent courses.

As with all studies that base viral detection around similarity to known viruses, our ability to detect novel viruses was limited by our reference dataset. Hence, some putative novel tick-borne viruses may have been missed due to the absence of similar viruses for comparison.

### Zoonotic risk assessment

The model applied showed promising results in distinguishing the zoonotic potential of closely related viruses within a genus. For instance, *Mivirus boleense* (15) was ranked as low zoonotic potential, while *Mivirus suffolkense* (20) was deemed high zoonotic potential. Given the validation of the methodology previously performed, this adds to the evidence suggesting the use of such models in providing targets for further research in a quick and low-cost way. However, caution must be exercised when interpreting the results of the genome composition-based model, as demonstrated in Figure 4B. The predicted probability interval was often large, leading to low confidence in the actual zoonotic risk of these viruses, and some viruses’ zoonotic risks were inaccurately estimated. For example, Issyk-kul virus had low predicted probability and associated “medium” risk, although it has been indicated as a likely human pathogen (68). Moreover, the model training did not include any virus in the recently described viral family *Chuviridae*, reminding that while these tools are powerful, predictions of zoonotic risks from viral families with limited data should be treated with an abundance of caution and greater weight should be given to other clinical or epidemiological sources of evidence. Despite these limitations, good performance was observed in the family *Nairoviridae*, which included only 13 viruses in the dataset with the model predicting high risks of zoonotic potential for viruses with the group, consistent with recent findings of new and emerging tick-borne viral diseases associated with nairoviruses and orthonairoviruses (4,5,62,69).

A significant proportion of tick-borne viruses with high or very high zoonotic potential were sampled from ixodid ticks, which may reflect sampling bias towards viruses infecting hard ticks (family *Ixodidae*) in previous studies or be true representation of differences in intrinsic zoonotic risk between the viral communities of argasid and ixodid ticks. The ecology of argasid and ixodid ticks are different, with ixodid ticks generally having longer blood meals (70). The longer feeding time of ixodid ticks may make them on average more favourable for viral transmission, and thus more likely to harbour communities of viruses capable of infecting mammals. However, it also means that ixodid ticks are more likely to be detected on a host, and thus more likely to be associated with resulting viral infections (i.e. a virus that might cause a fever of unknown origin if transmitted by an argasid tick may be able to be explicitly linked to a tick bite given the greater detection probability if transmitted by an ixodid tick). A more systematic approach that samples both hard and soft ticks would increase our knowledge of the diversity of tick-borne viruses carried by each type and allow us to assess whether they truly do differ in human infection risk.

As the model was trained on complete genomes (31), it is inappropriate for the use on viruses for which only partial genomes are available. This is a particularly important limitation for viruses found purely through bioinformatic means, where only single contigs or disconnected viral segments may be available. We attempted to assess the zoonotic risk of the putative novel viruses detected in this study by using the genomes of the closest related virus with a complete genome as a proxy. However, while this was unavoidable in this case, it does fall prey to one of the same biases the use of genome-based composition methods was originally intended to avoid, the assumption that zoonotic risk is conserved across the phylogeny. Hence, the zoonotic risks of these viruses ought to be reassessed should full genomes become available.

## Conclusion

Despite ticks being important vectors of pathogens, the human infection risks of most tick-borne viruses remain uncertain (71). In this study we have both identified new tick viruses in previously collected data and assessed the zoonotic potential of known tick virus diversity, identifying certain high risk viral families deserving of further study and, potentially, surveillance. Cataloging viruses and assessing their human infection risk, as we have done here, represents only a first step in the risk management of zoonoses. Further research into the ecological processes that underlie the geographical distribution and interspecific transmission of ticks and their viruses is necessary to gain a comprehensive understanding of the ecology of this system. It is this eventual ecological knowledge that may allow us to formulate more effective strategies for managing human exposure to vectors, preventing zoonoses at the source.

## Data accessibility statement

All novel viral sequences identified in this study have been submitted to Third Party Annotation Section of the DDBJ/ENA/GenBank databases under the accession numbers TPA: BK062905-BK062914. These sequence records will be held confidential until the data or accession numbers appear in print. Data and scripts used to generate the analyses are provided in at https://github.com/ytlin2021/TickVirus.git.

## Author Contributions

Yuting Lin: Project administration (lead); methodology (equal); data curation (lead); investigation (lead); formal analysis (equal); visualisation (lead); writing – original draft (lead); writing – review and editing (equal). David J Pascall: Conceptualization (lead); methodology (equal); formal analysis (equal); resources (lead); funding acquisition (lead); supervision (lead); writing – review and editing (equal).

## Acknowledgements

We thank members of the Medical Research Council (MRC) Biostatistics Unit at the University of Cambridge (MRC BSU) for scientific input. We are grateful for the technical support of core facilities the Cambridge University High Performance Computing Service.

DJP is funded by UKRI (UK Research & Innovation) through the JUNIPER consortium (MR/V038613/1), and through the MRC Biostatistics Unit core award from the MRC (MC UU 00002/11).

## Appendices

**Appendix Figure 1.**
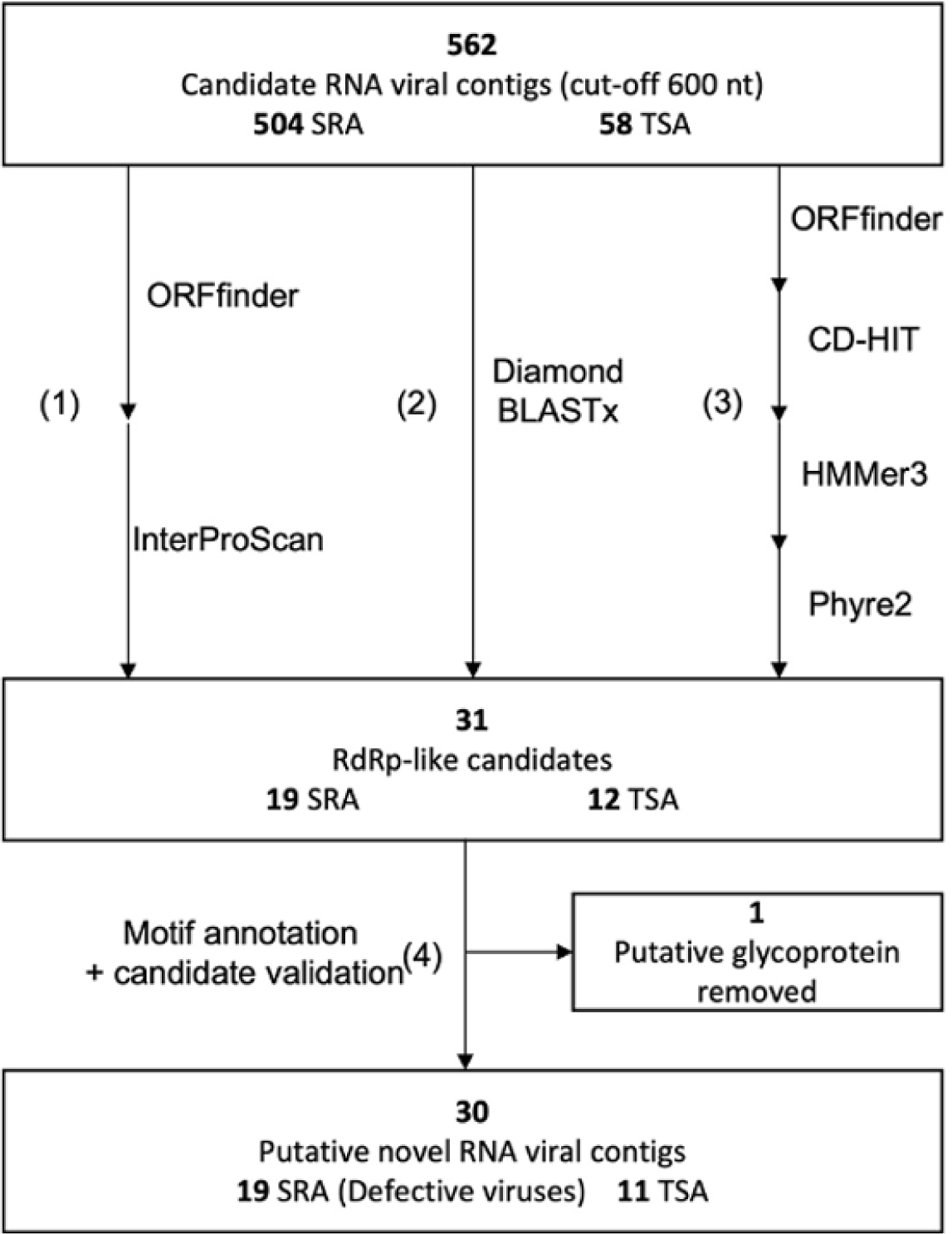
Workflow of detecting viral RNA-dependent RNA polymerase sequences. (1) ORFs were translated using ORFfinder (cut-off 200 aa). The translated sequences were predicted using the InterProScan-5.55-88.0 (--minsize 600; -goterms). (2) BLASTx using Diamond v2.0.14.152 against both non-redundant protein database and RdRp-scan database (-e 1e-5 --min-orf 600 --very-sensitive). (3) ORFs were translated using ORFfinder (cut-off 200 aa). Redundant sequences were then removed using CD-HIT v4.6.8 at 90% level of identity (-c 0.98 sequence identity threshold). Remaining sequences were scanned using HMMer3 v3.2.1 against RdRp HMM profile database (full sequence e-value 1e-06). HMM-RdRp hits were compared against PDB using Phyre2 server. In total, 31 RdRp-like candidates from SRA and TSA databases were found using RdRp-scan and InterProScan after removing duplicates. (4) RdRp motifs were mapped to candidate sequences using Geneious v2022.1. RdRp-scan database, RdRp HMM profile database, RdRp motifs database and RdRp key word lists were obtained from Charon et al. (77) (https://github.com/JustineCharon/RdRp-scan).

**Appendix Figure 2.**
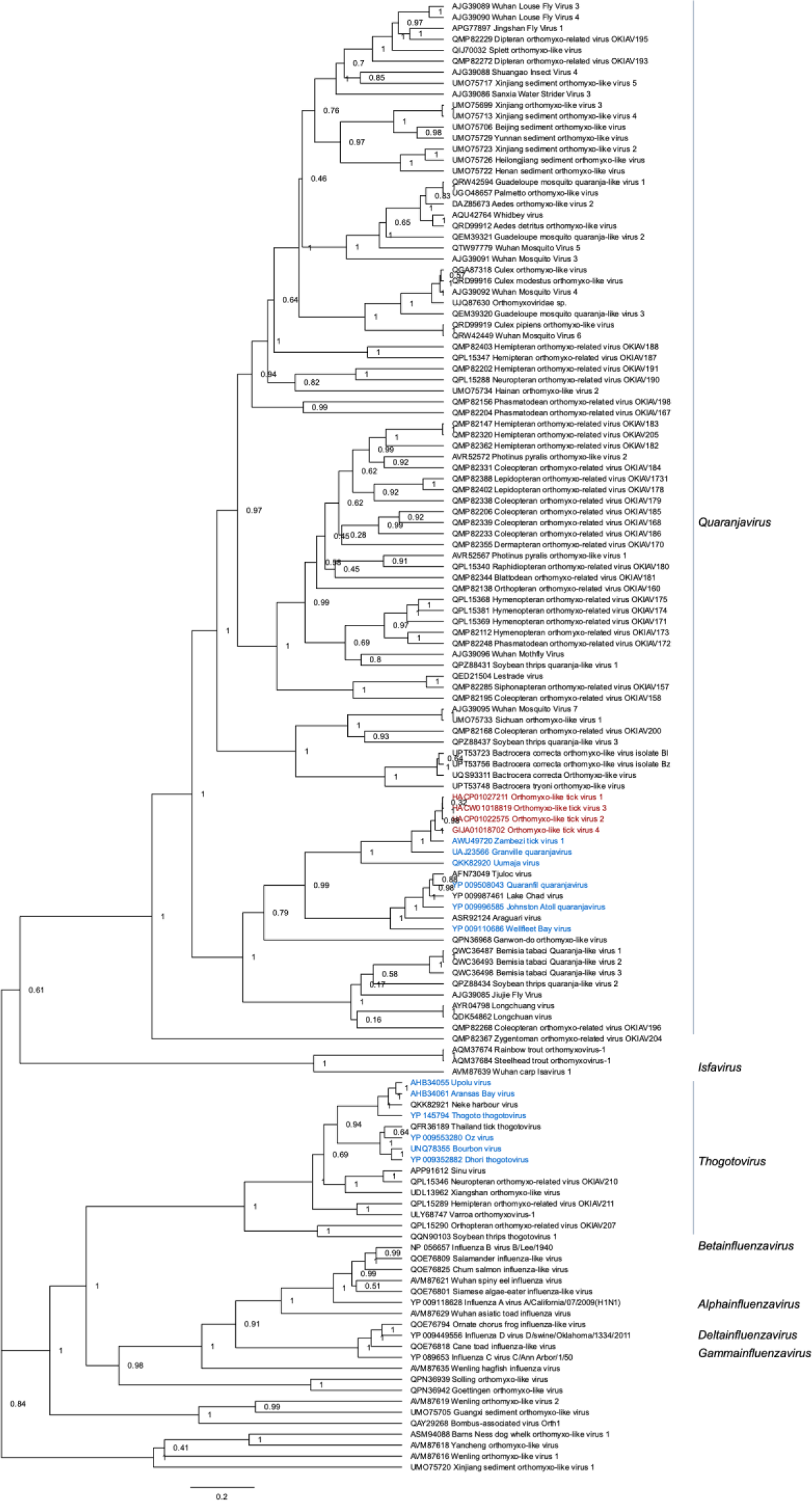
Phylogenetic relationship of the novel Orthomyxo-like viral contigs. Maximum clade credibility tree was inferred from the Bayesian posterior sample. Putative novel viral contigs found in the present study are shown in red, and previously known tick-borne viruses are shown in blue.

**Appendix Figure 3.**
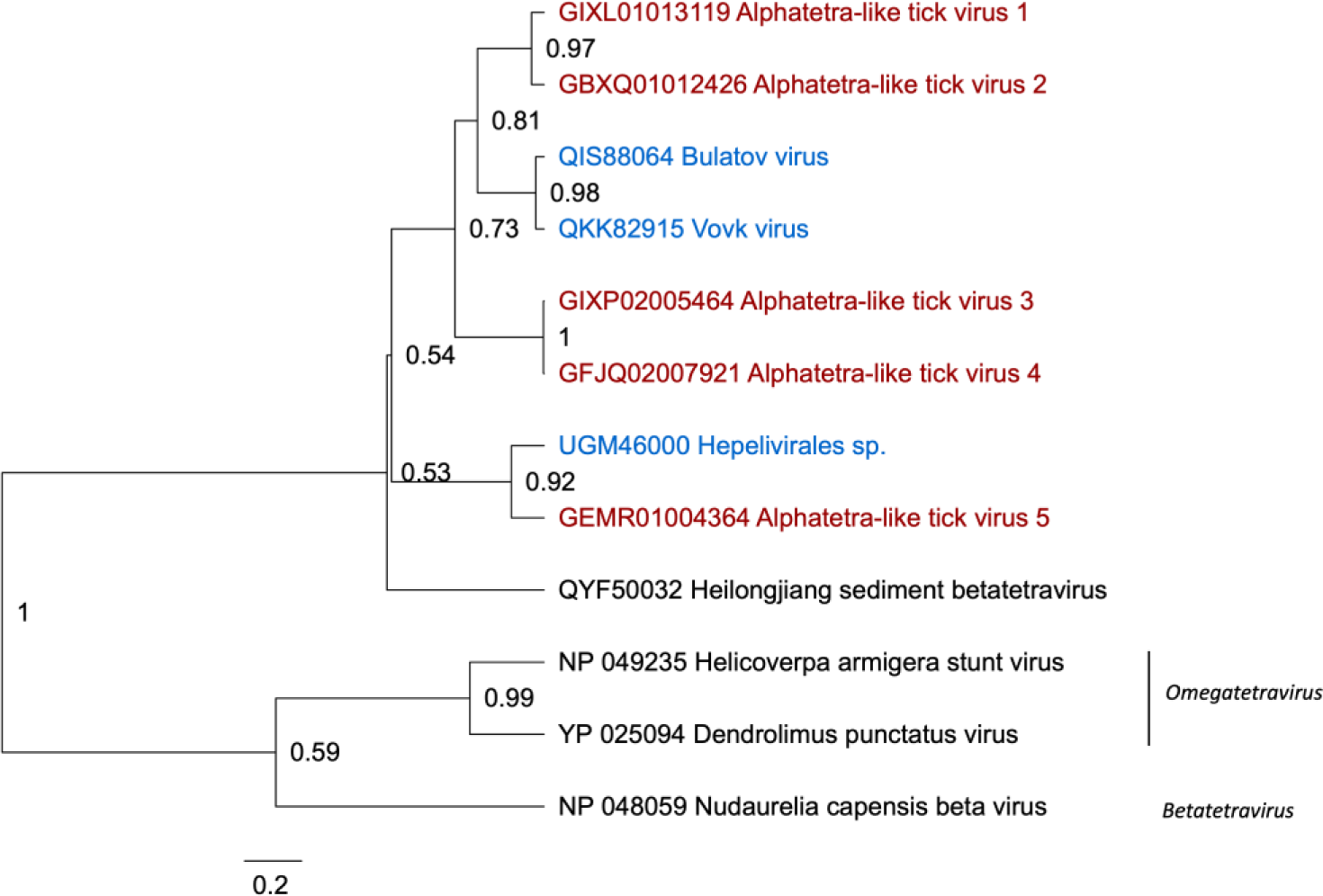
Phylogenetic relationship of the novel Alphatetra-like viral contigs. Maximum clade credibility tree was inferred from the Bayesian posterior sample. Putative novel viral contigs found in the present study are shown in red, and previously known tick-borne viruses are shown in blue.

**Appendix Figure S4.**
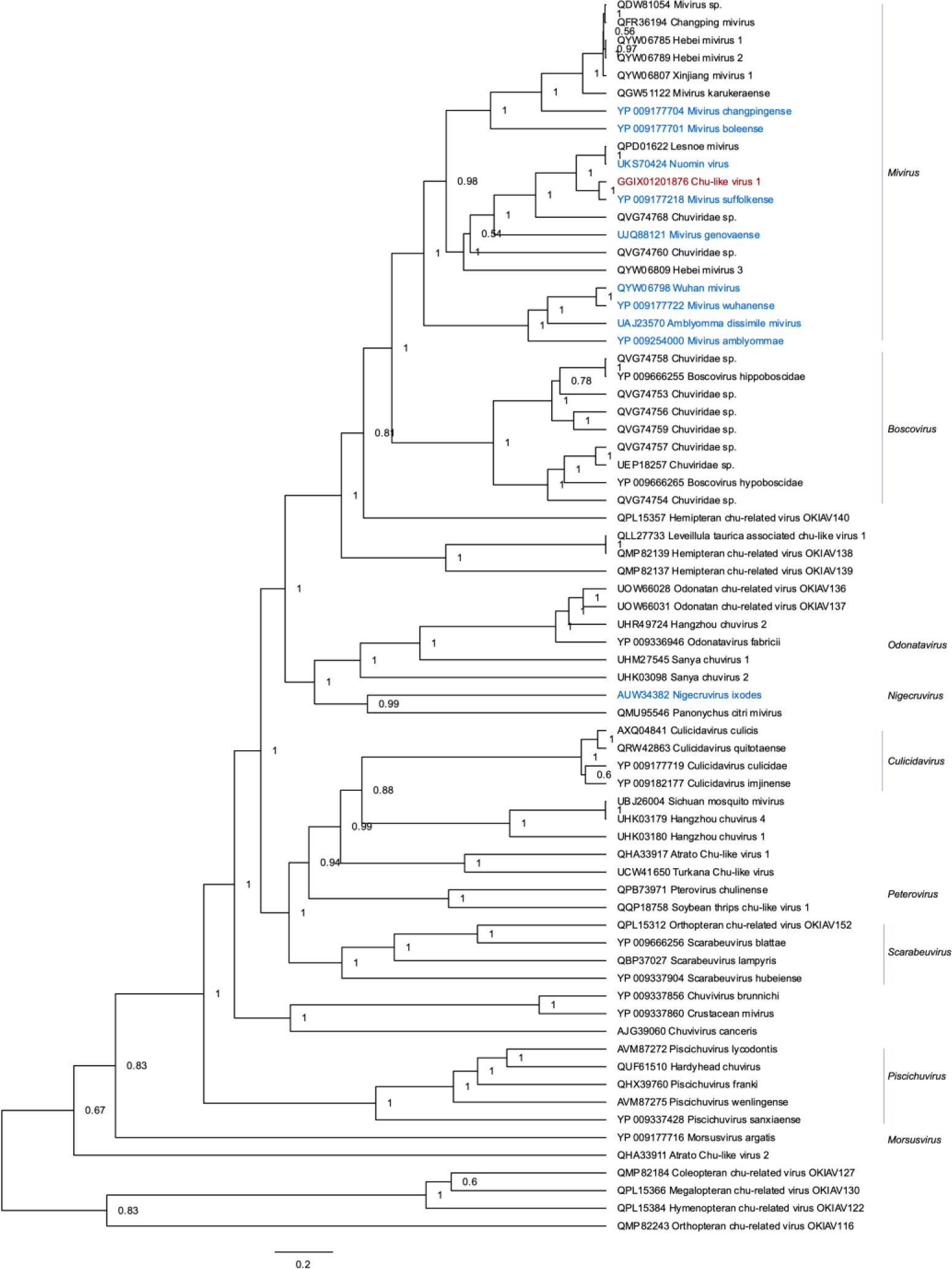
Phylogenetic relationship of the novel Chu-like viral contigs. Maximum clade credibility tree was inferred from the Bayesian posterior sample. Putative novel viral contigs found in the present study are shown in red, and previously known tick-borne viruses are shown in blue.

**Appendix Table 1.**
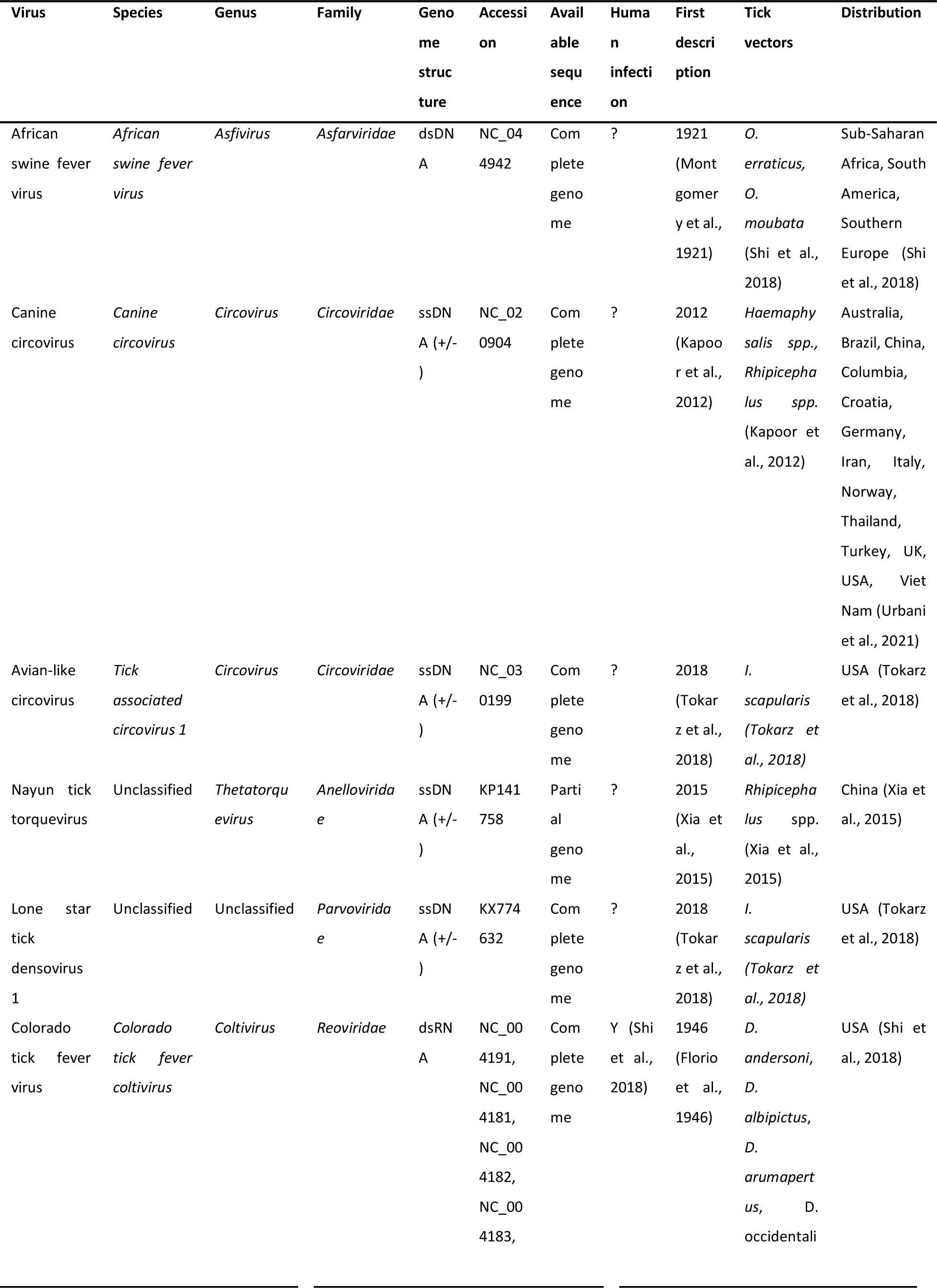
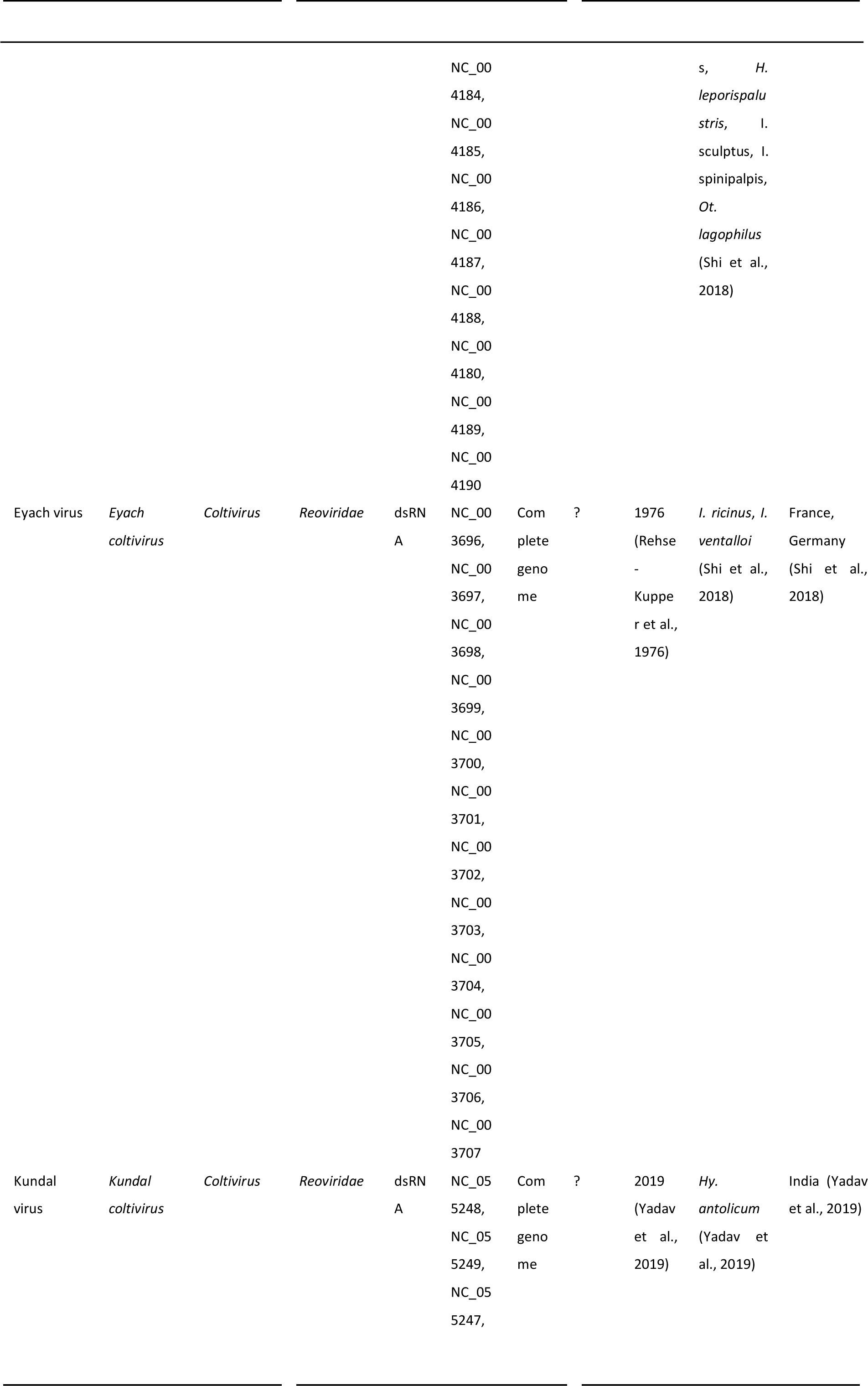
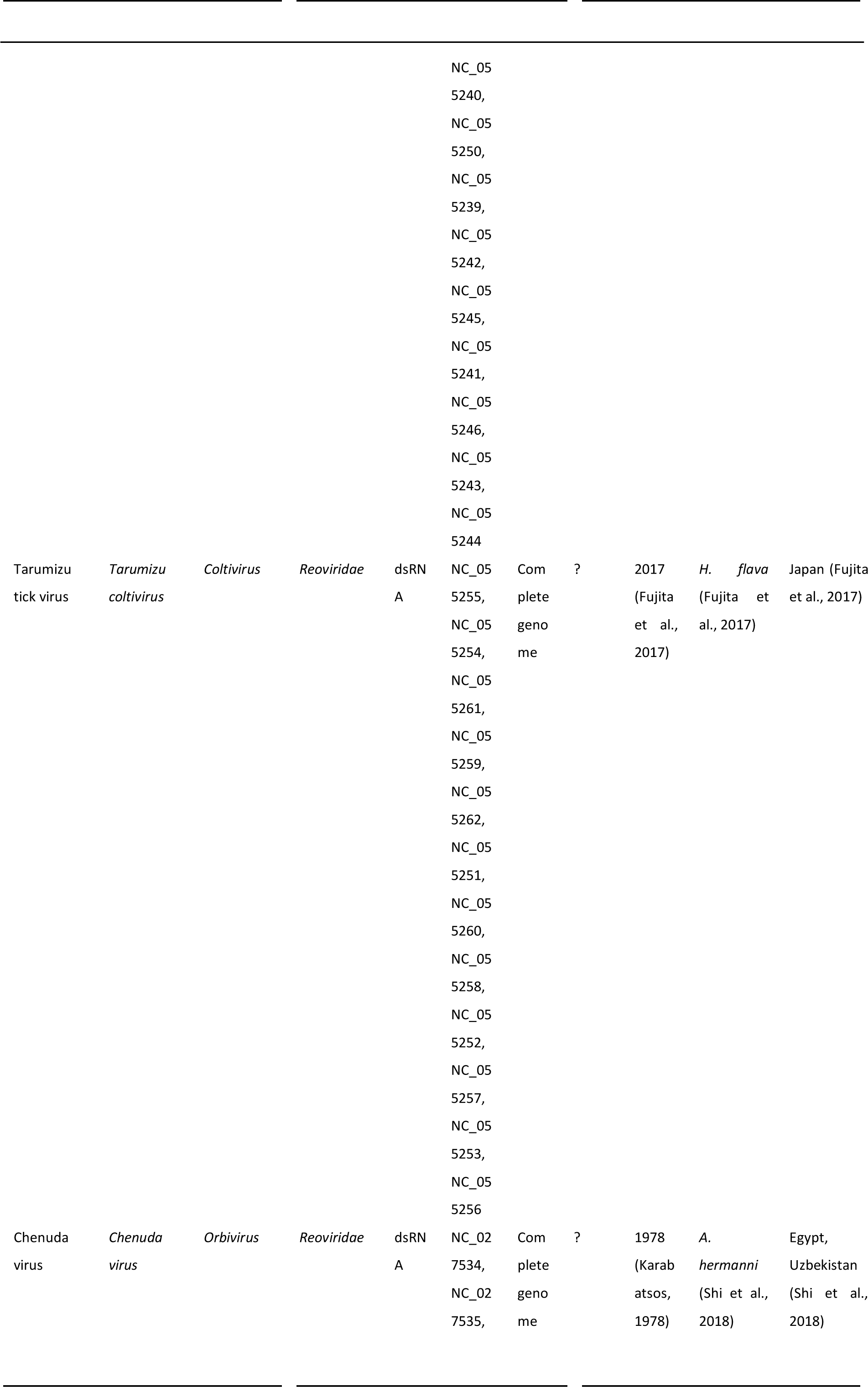
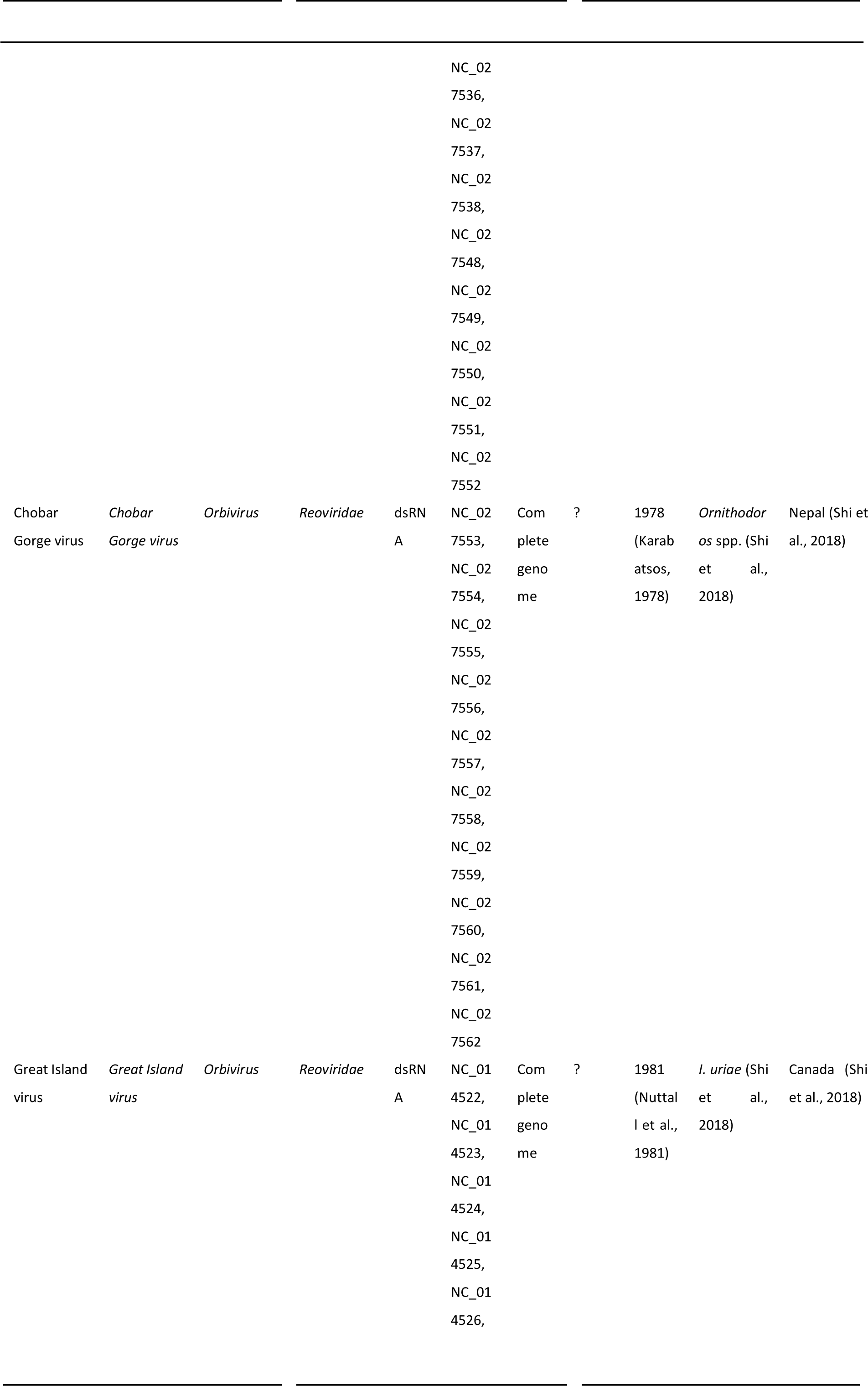
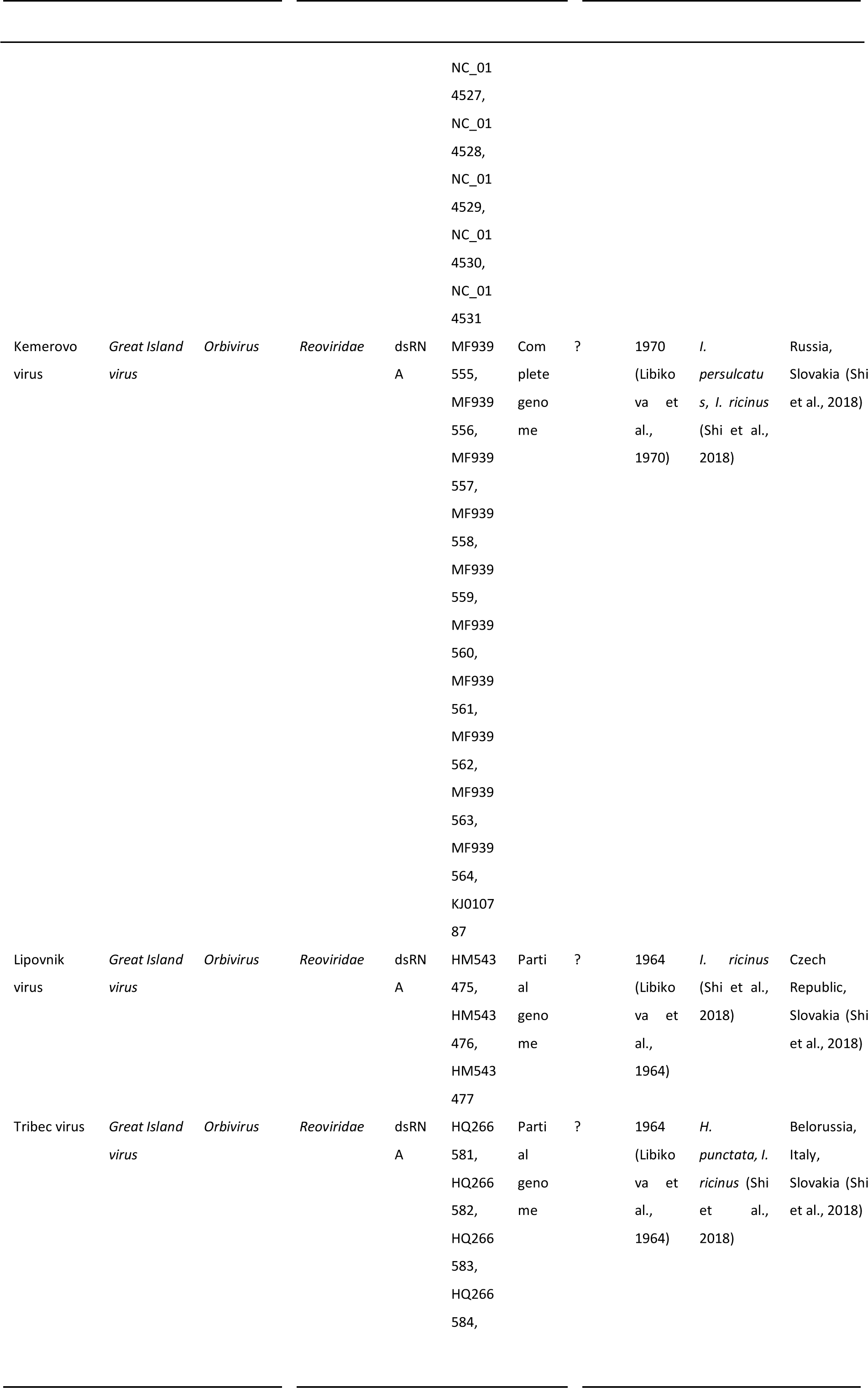
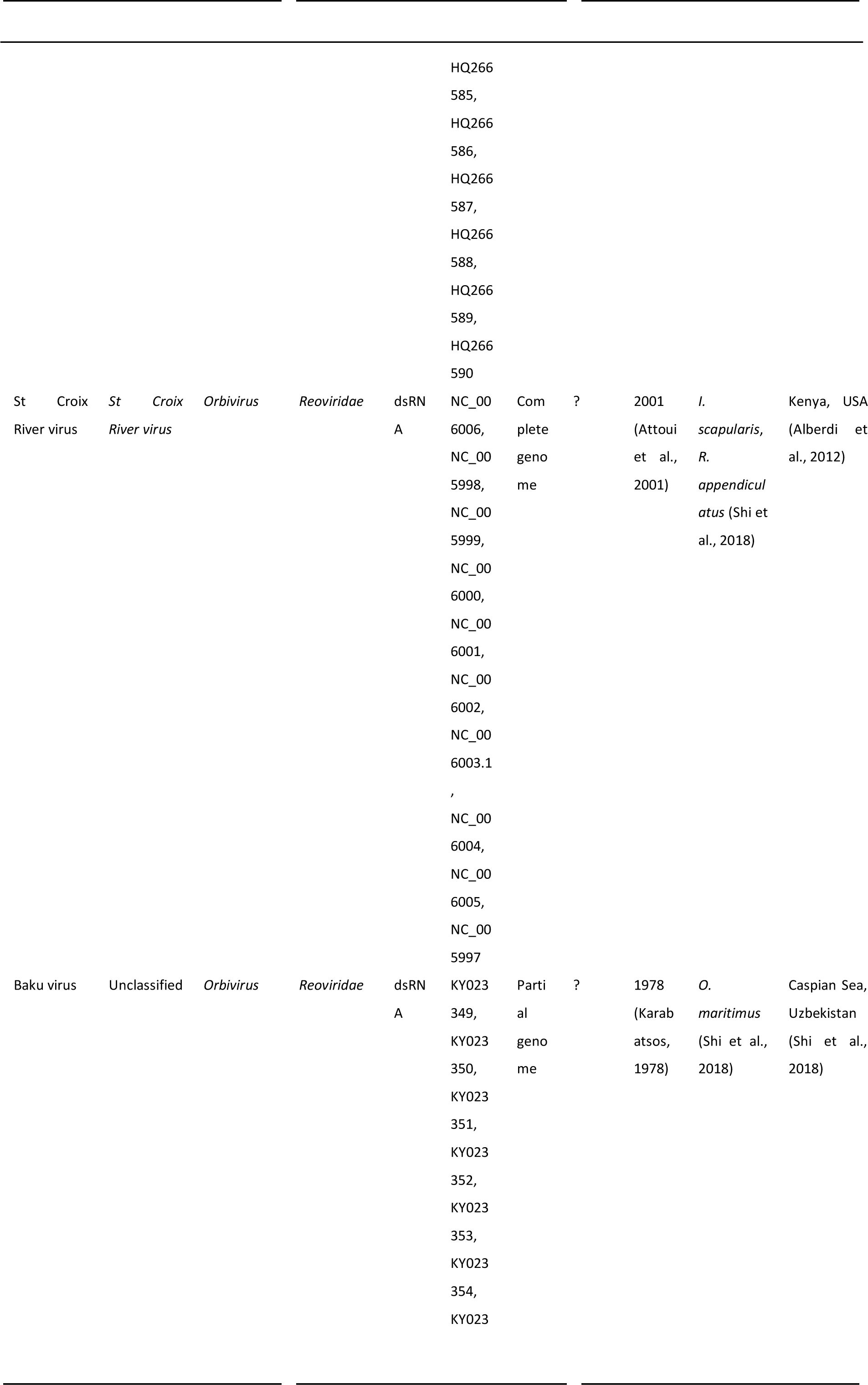
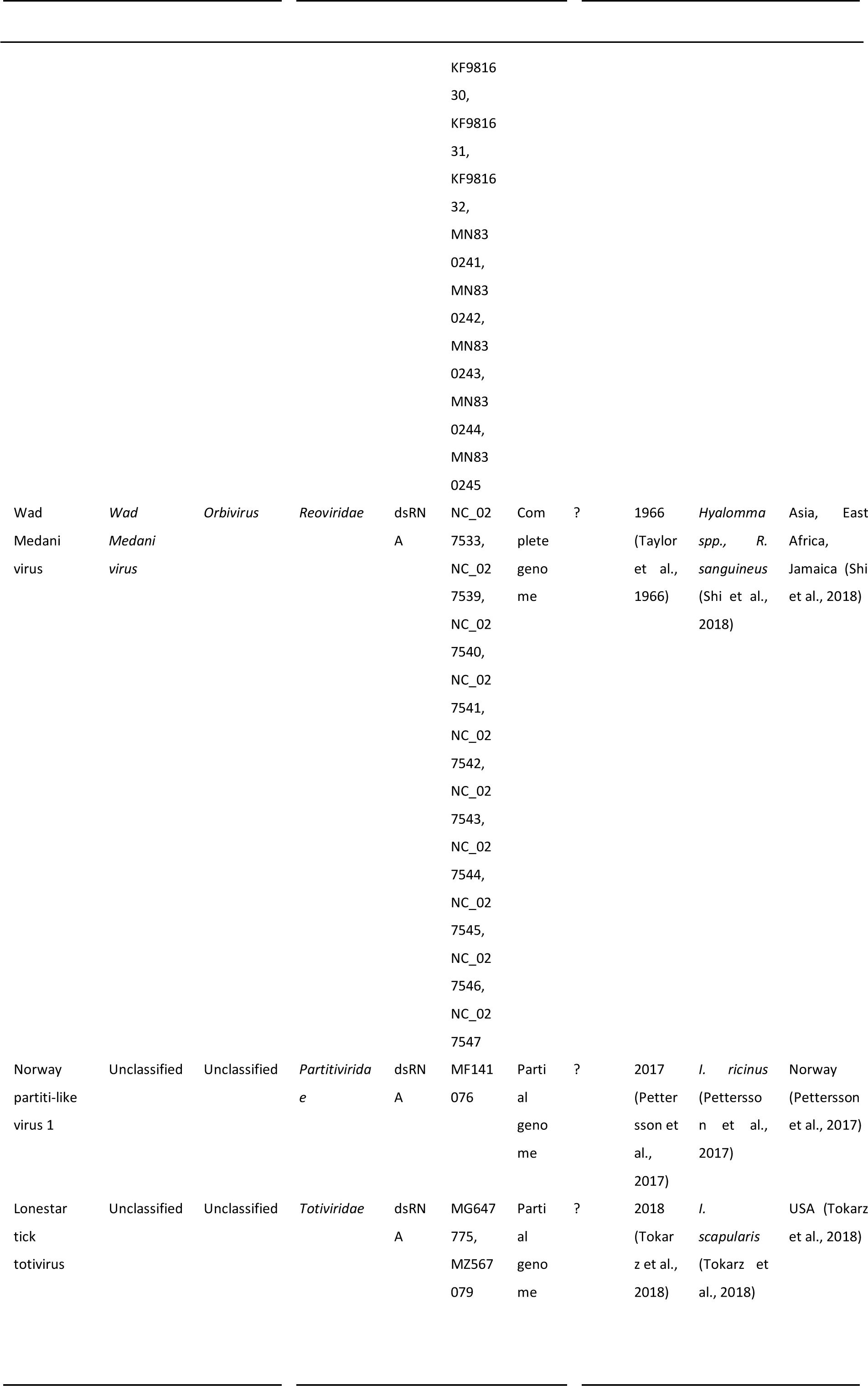
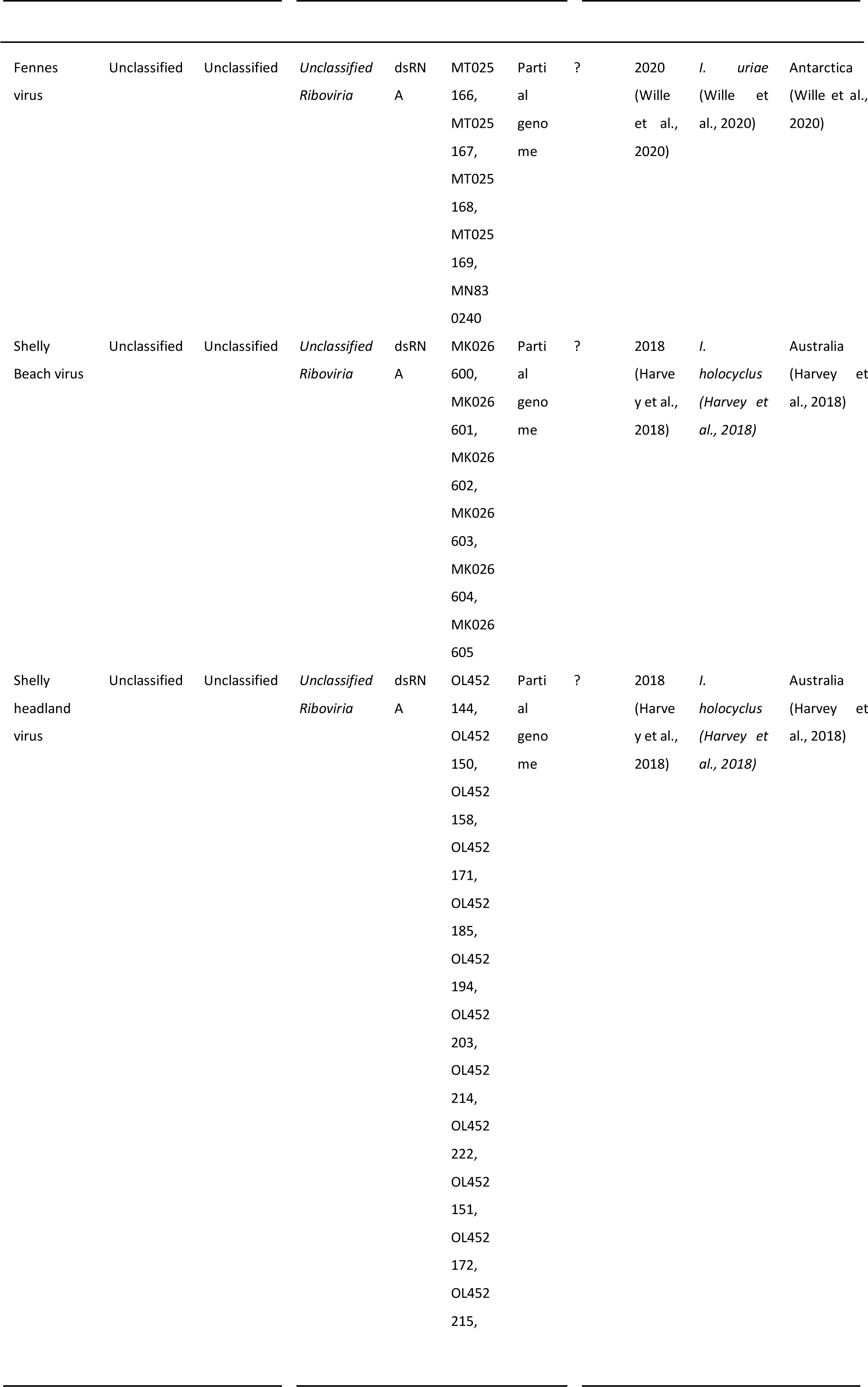
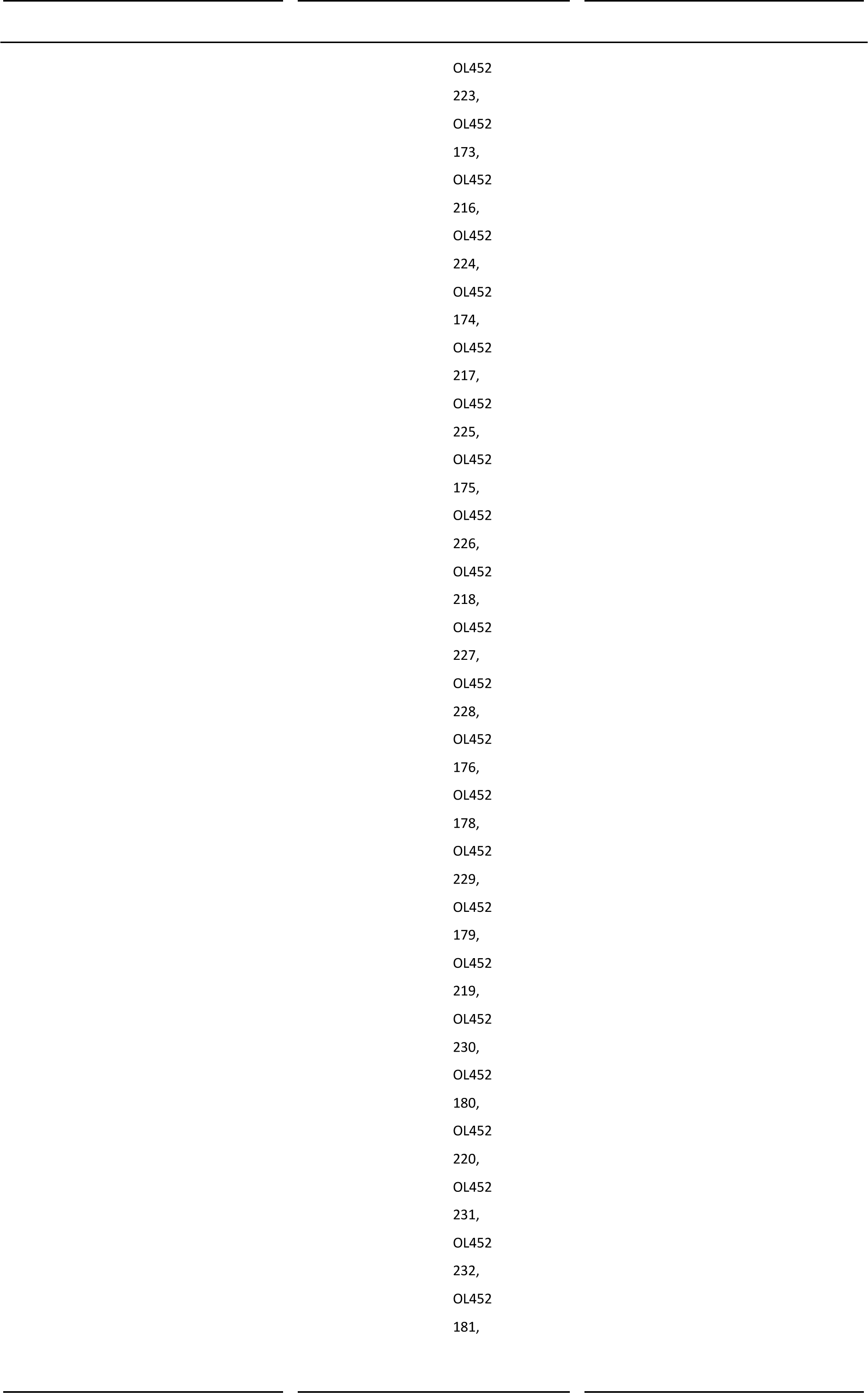
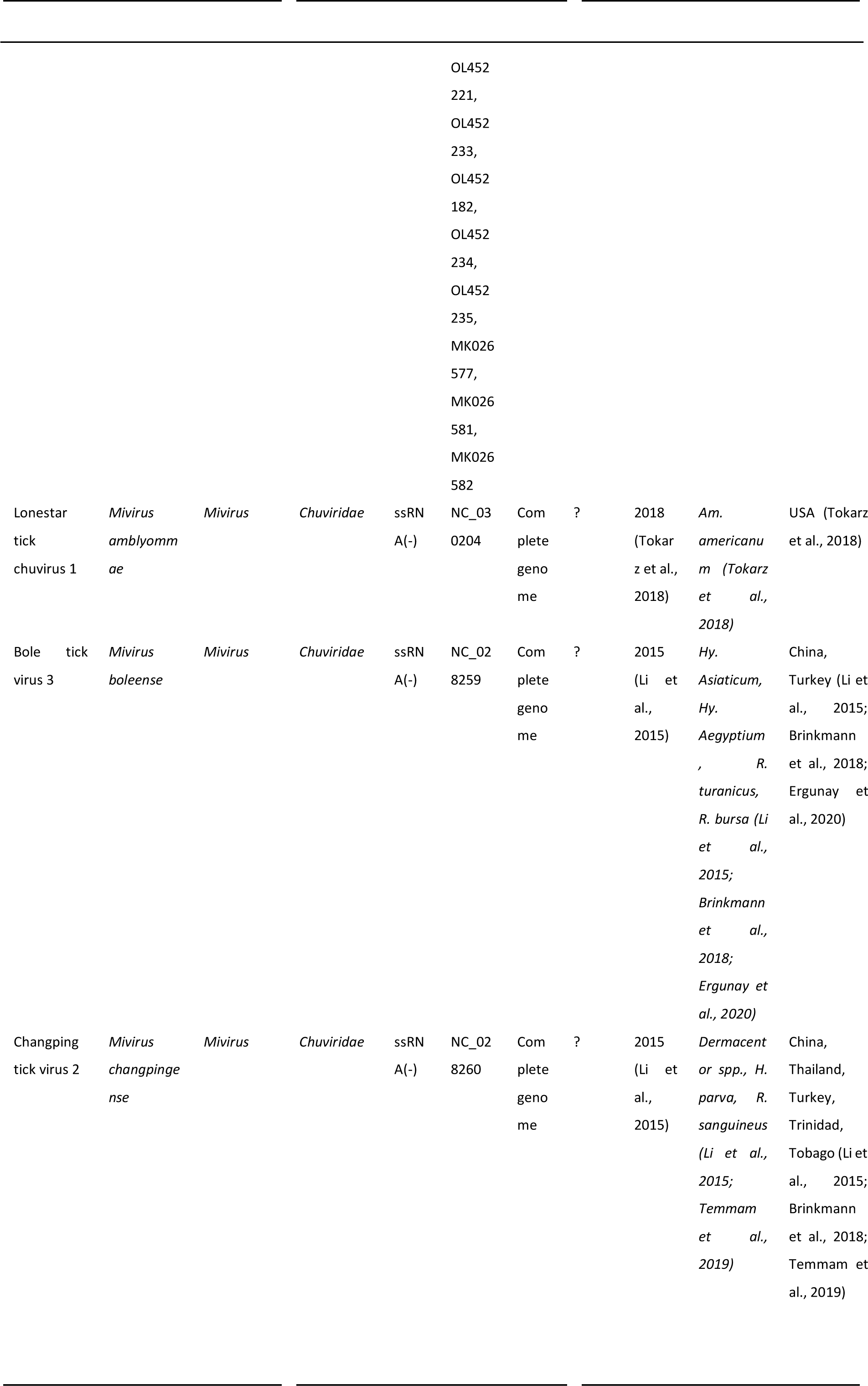
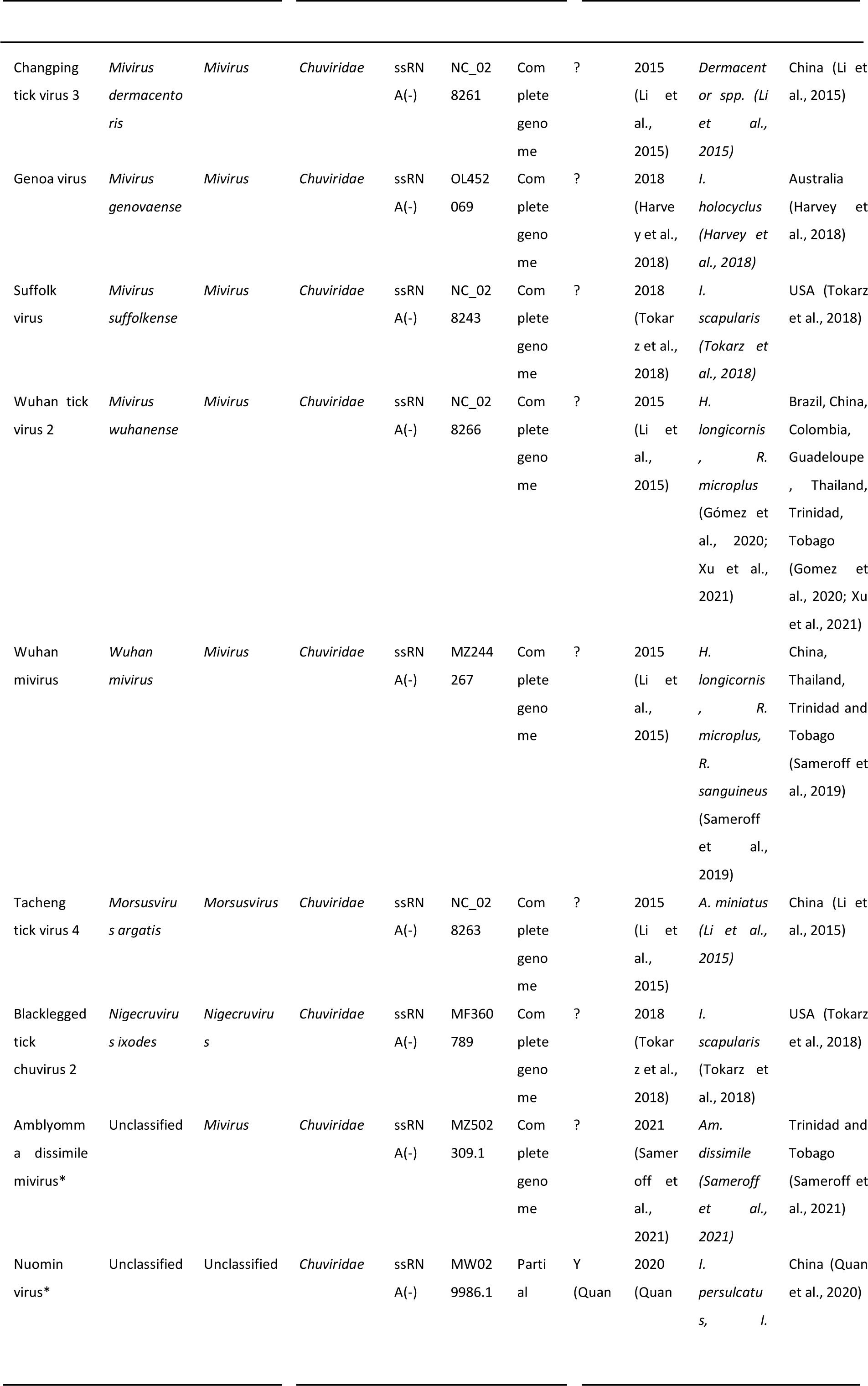
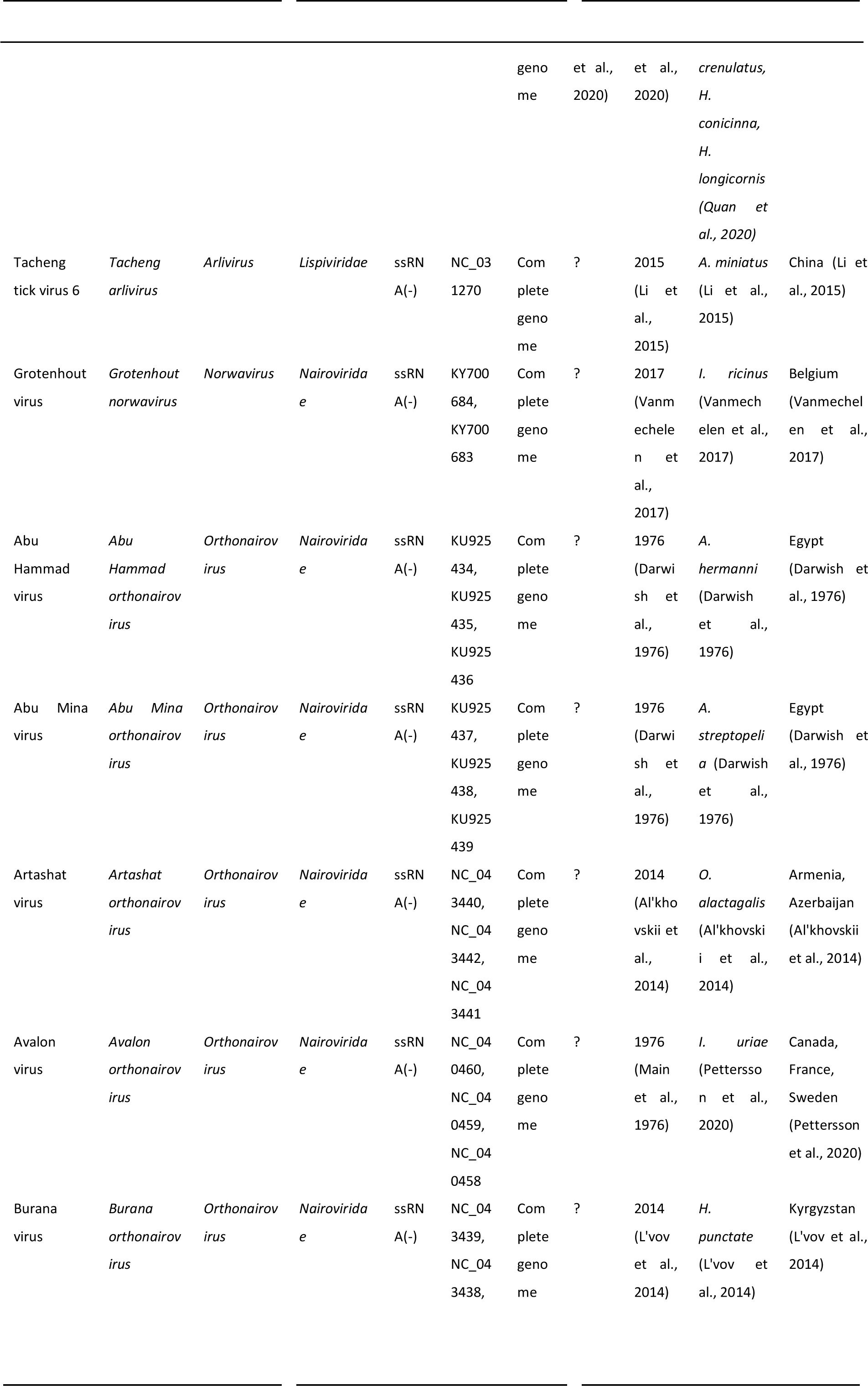
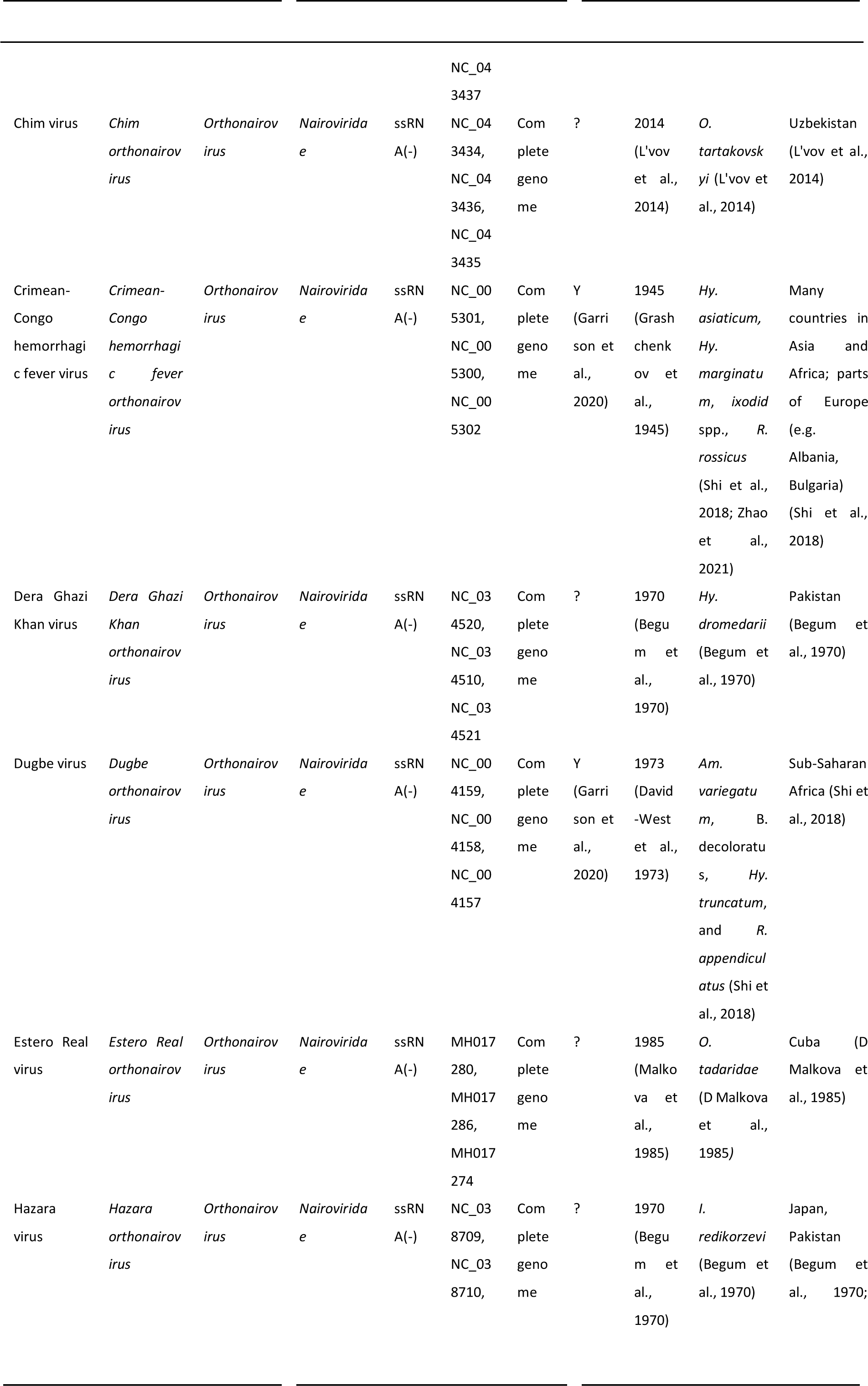
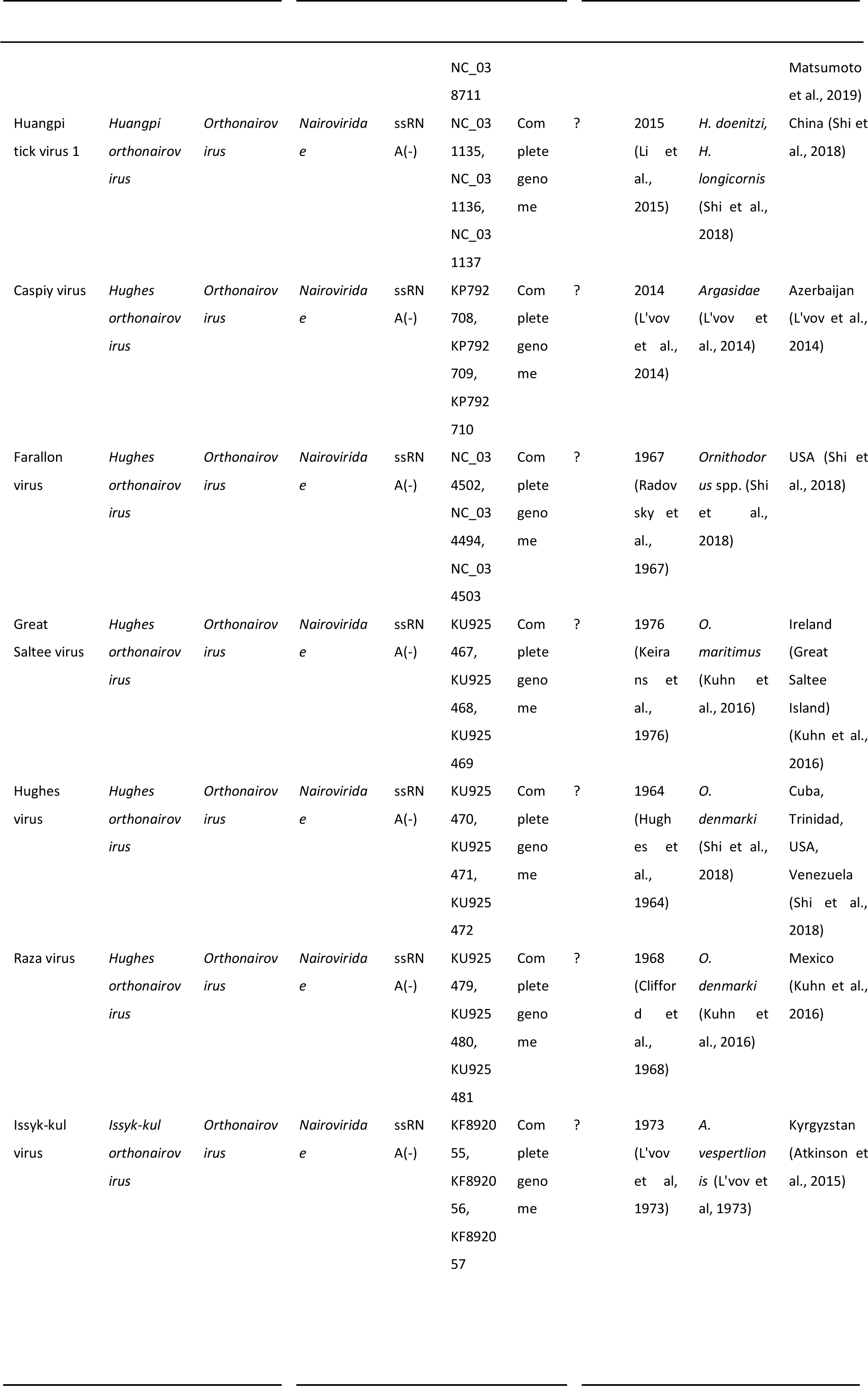
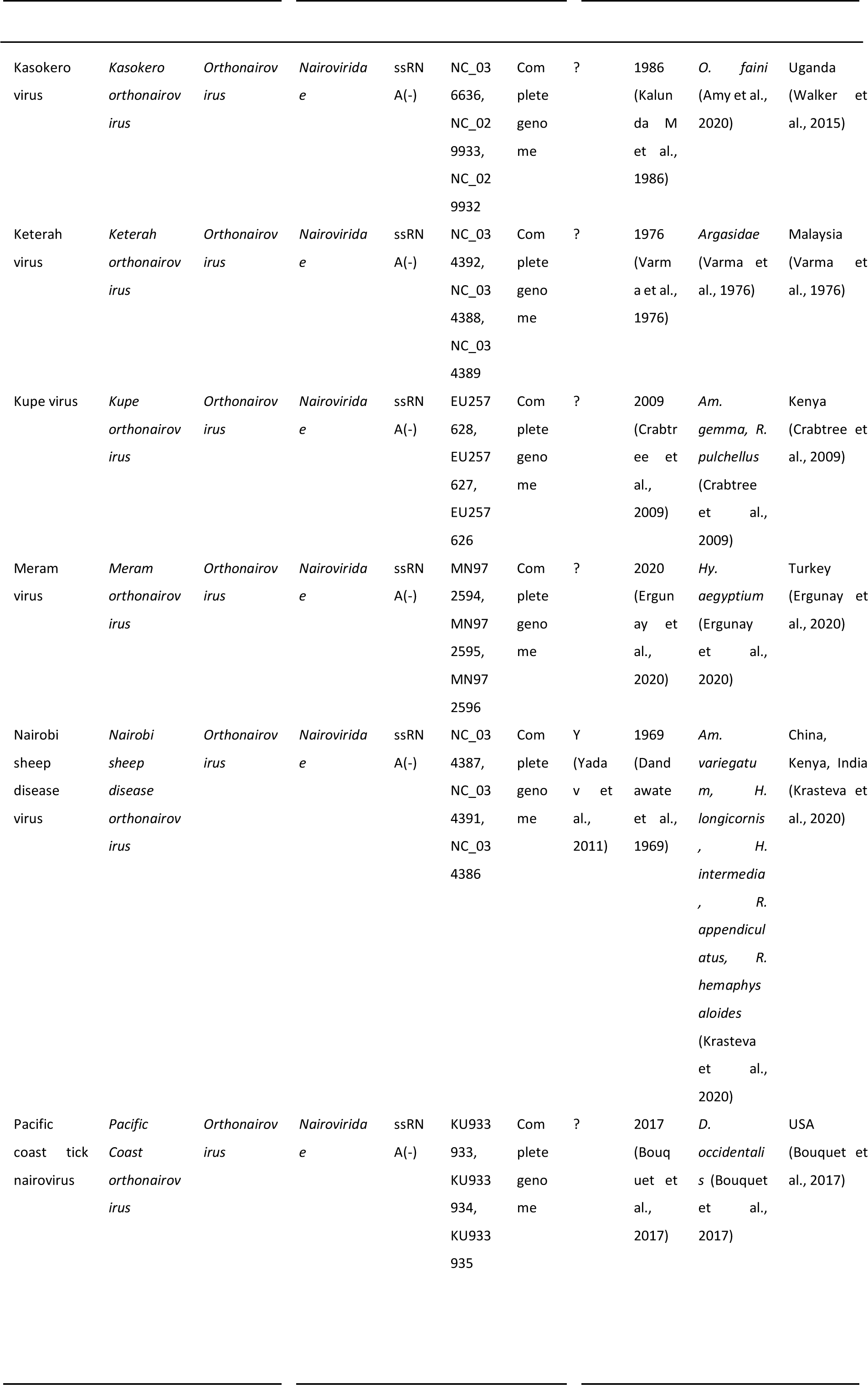
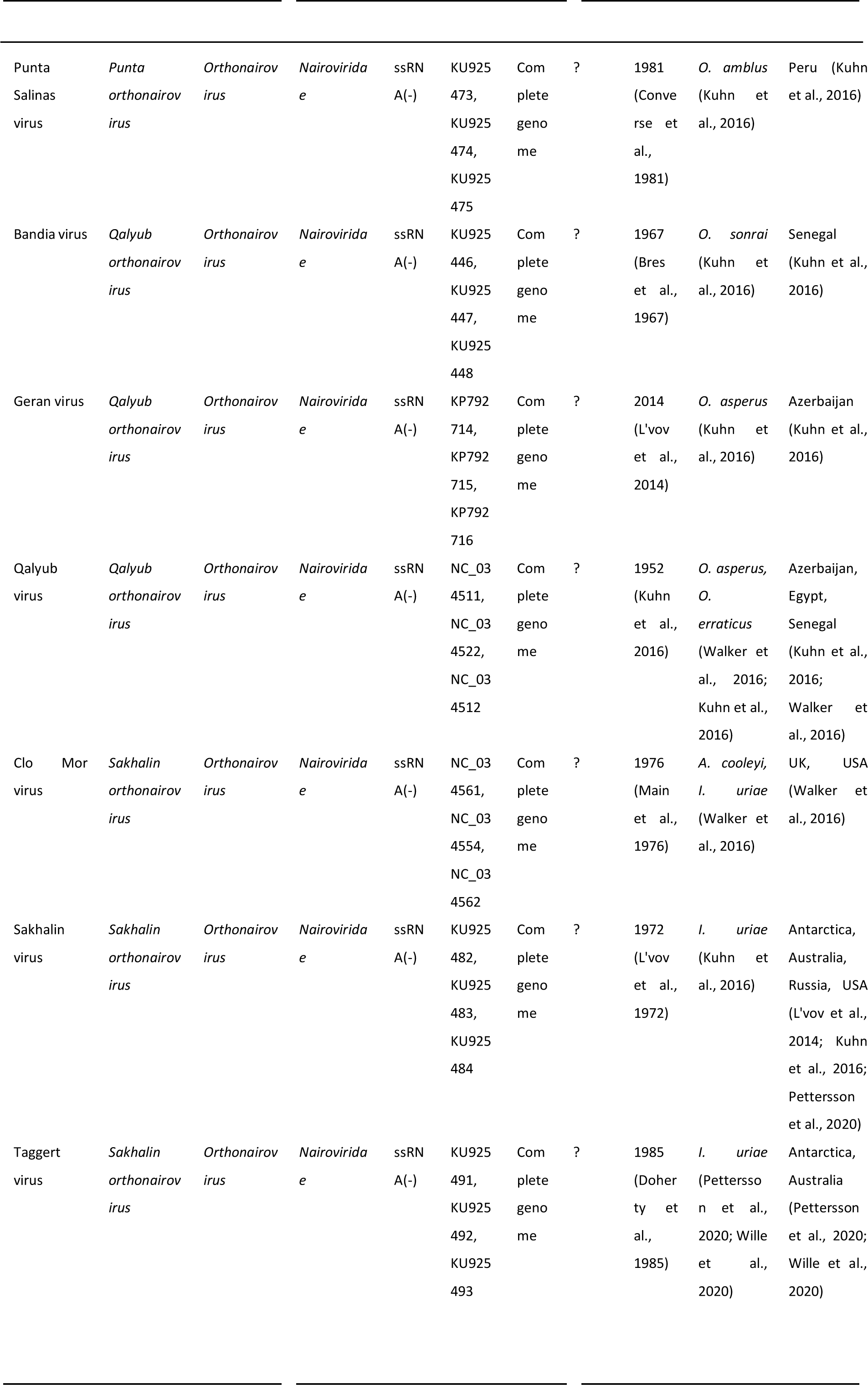
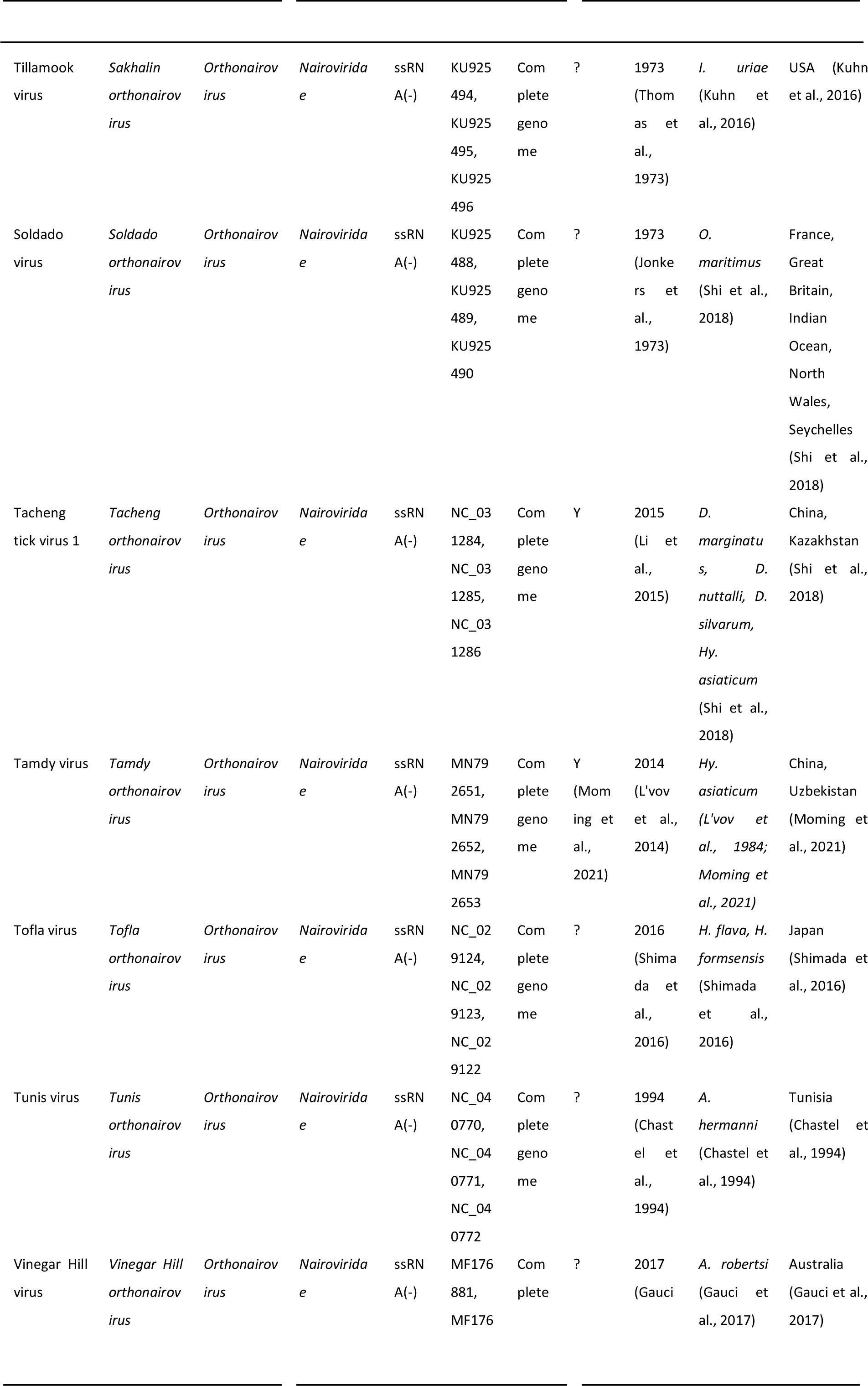
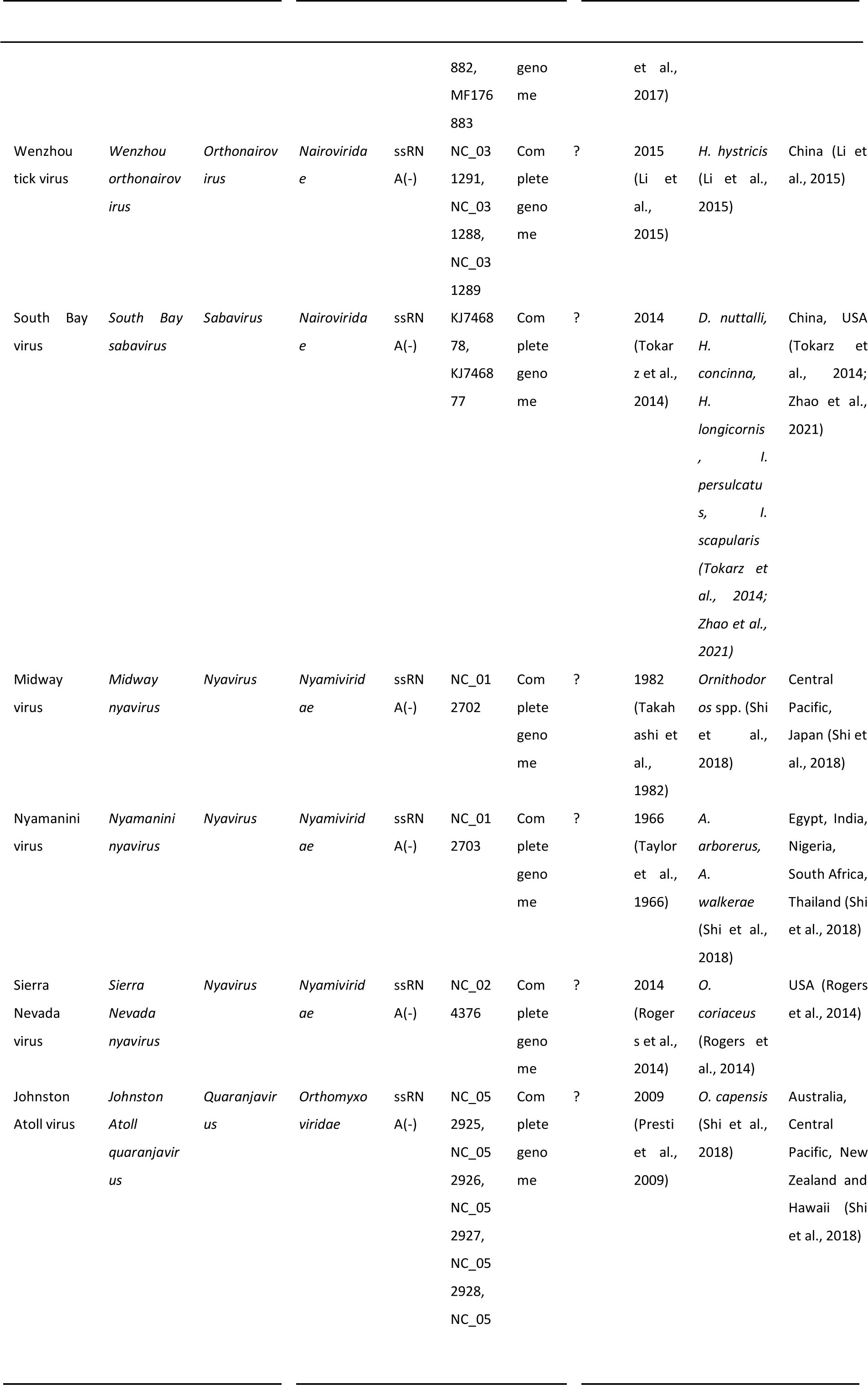
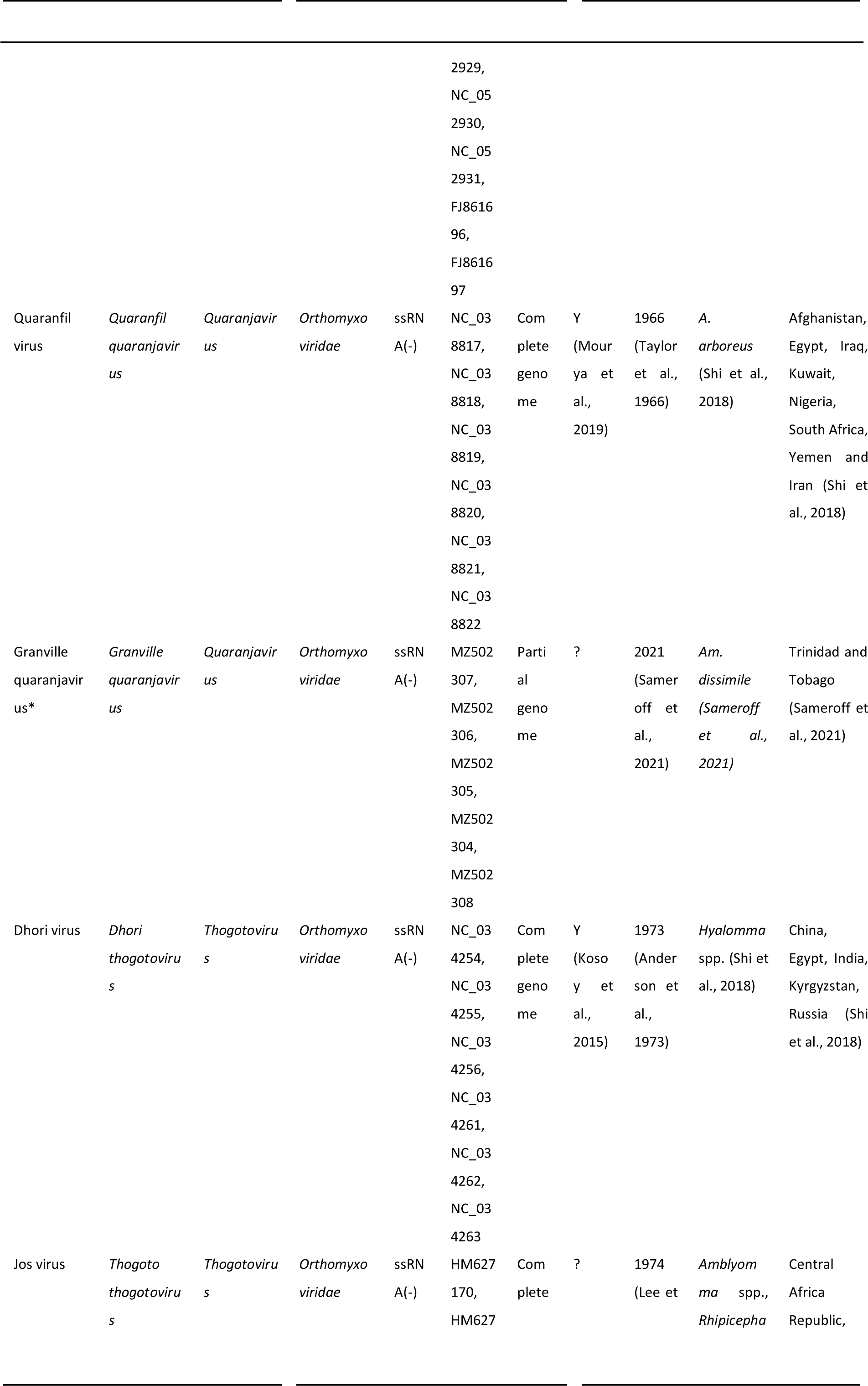
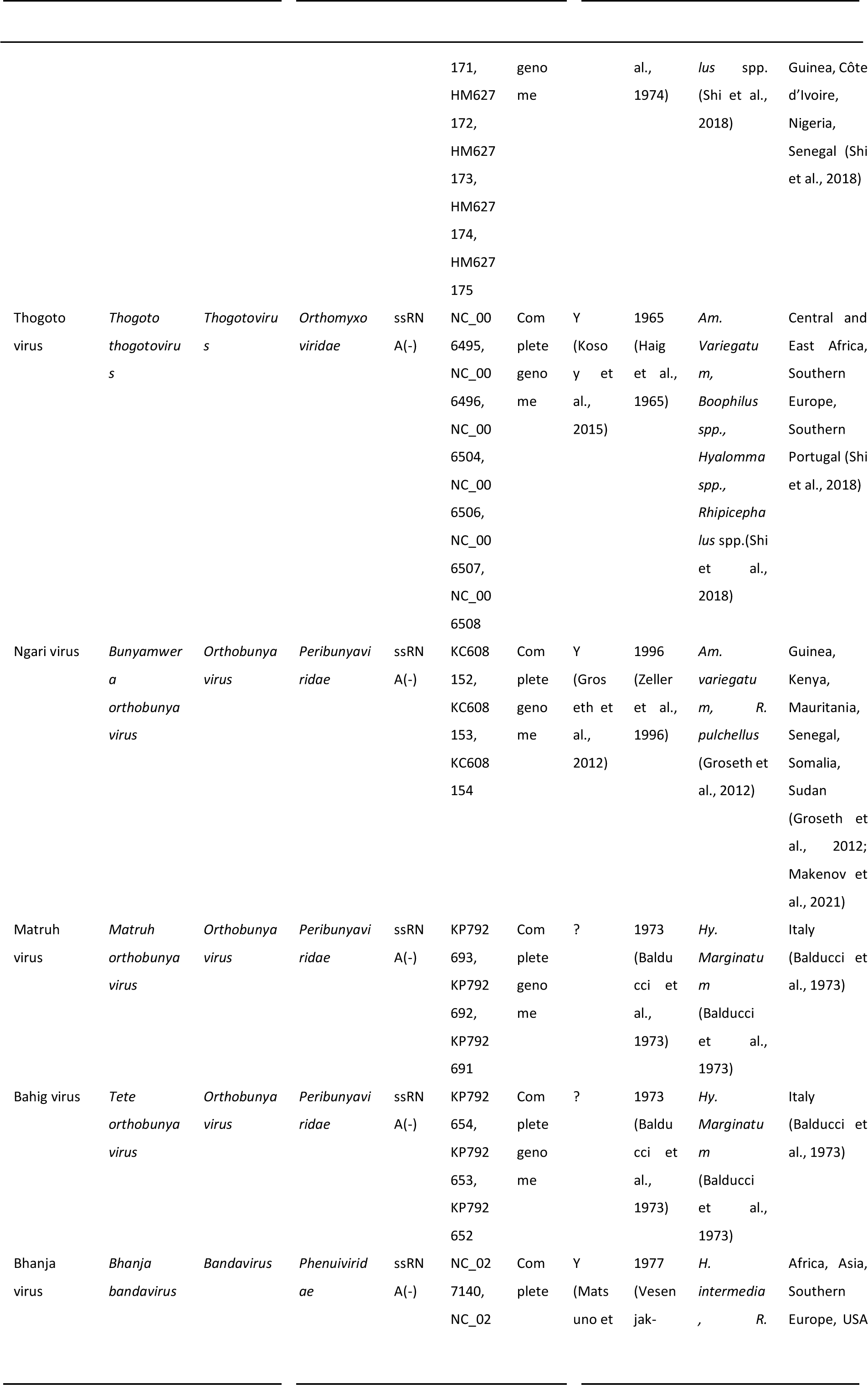
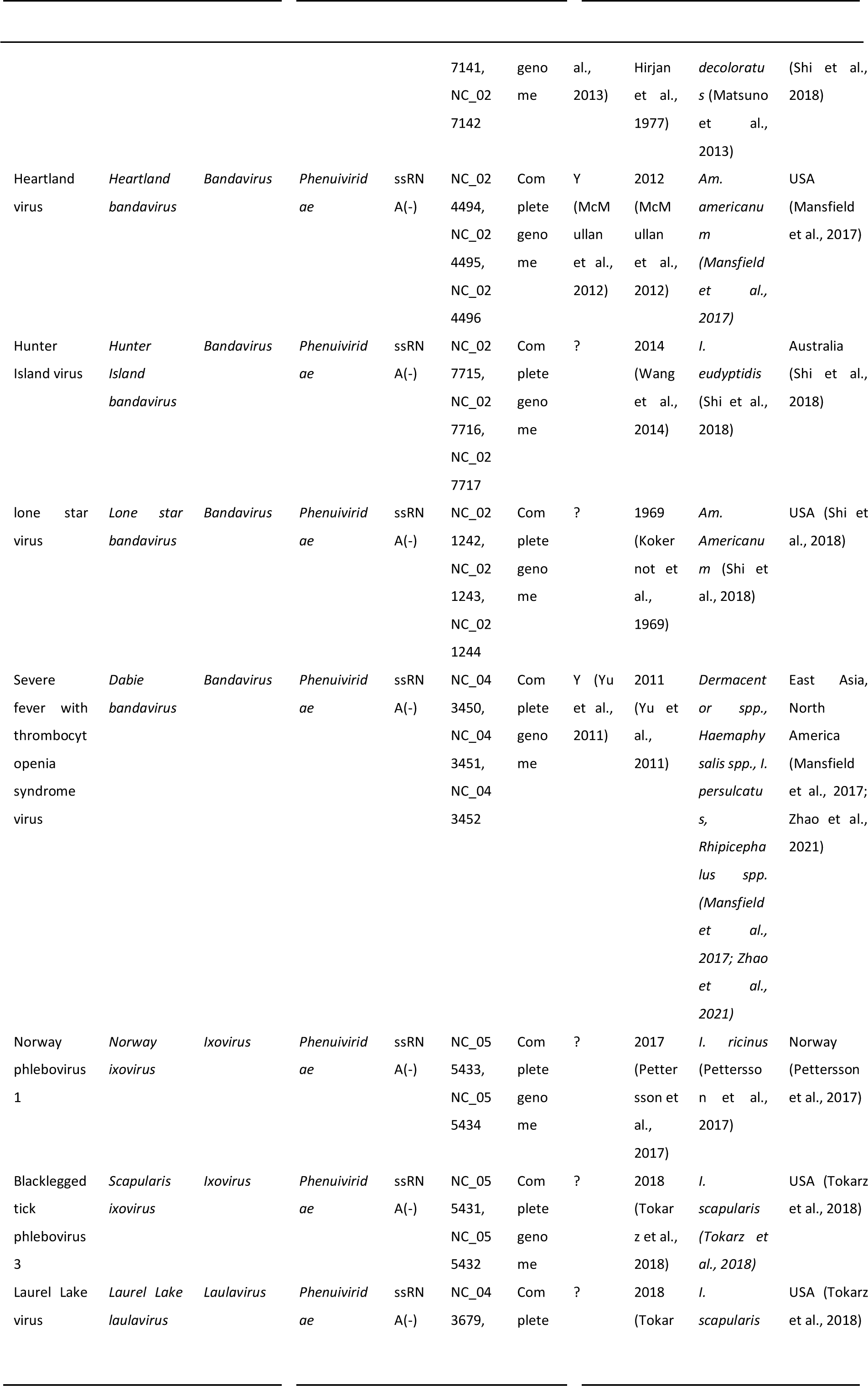
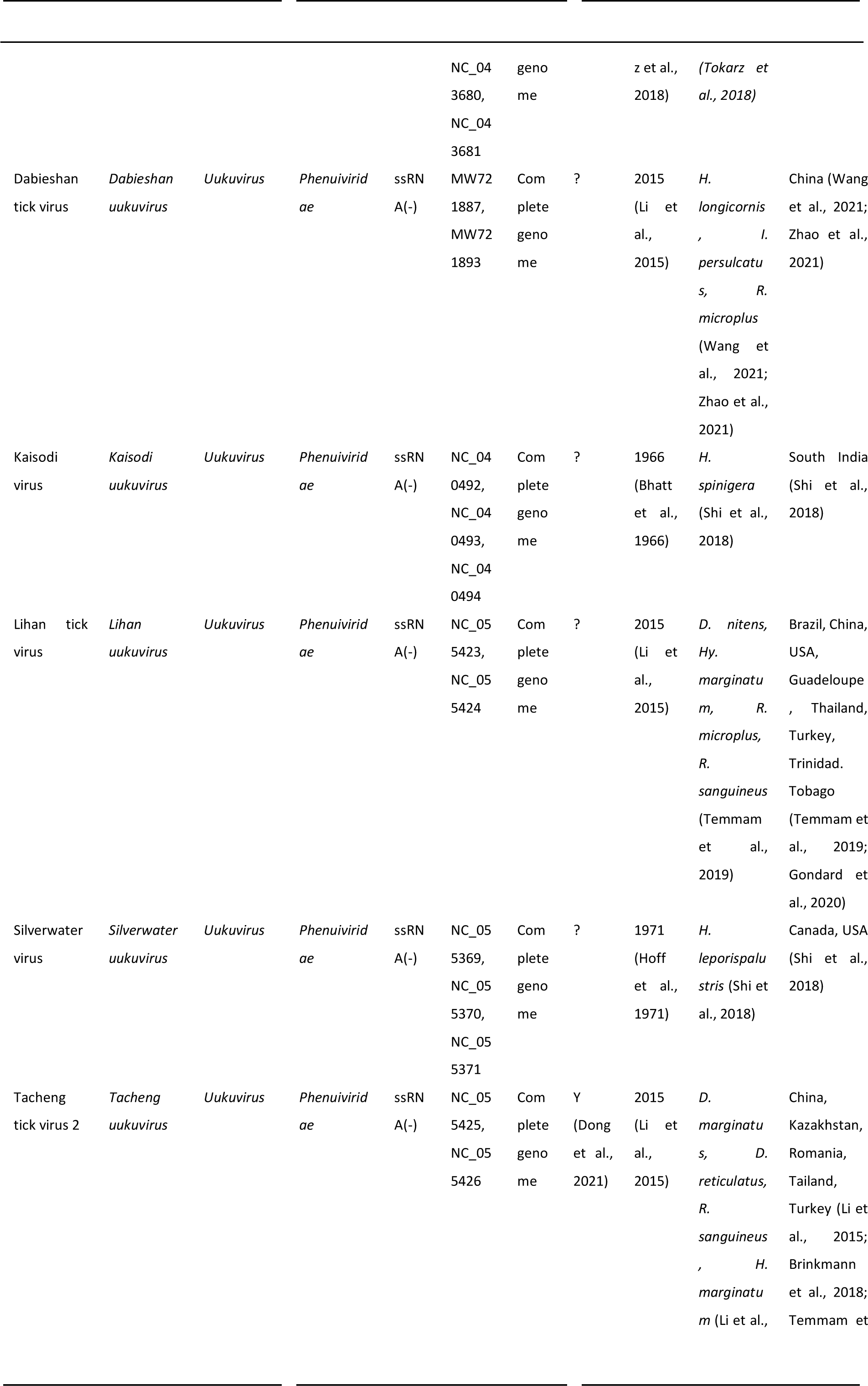
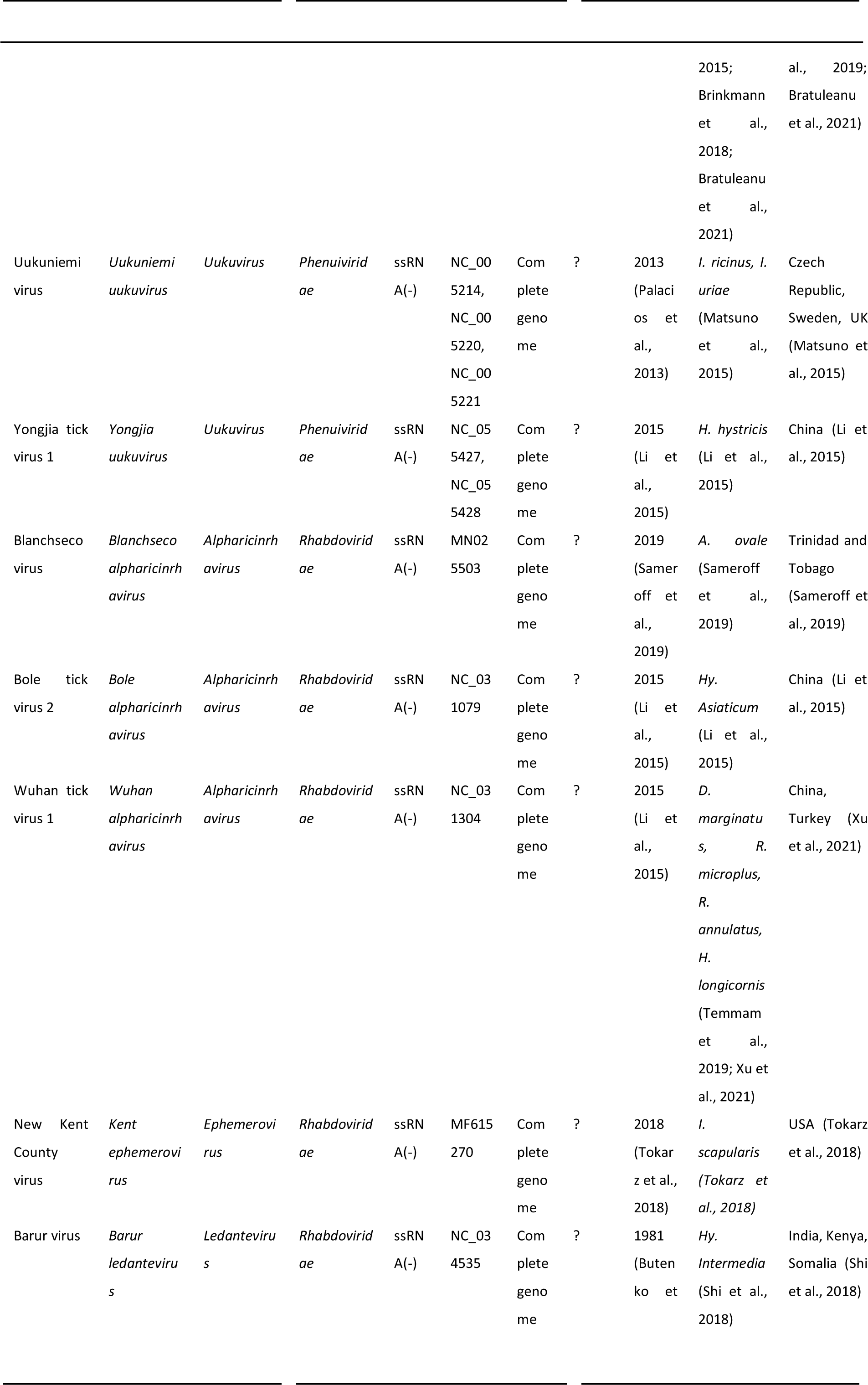
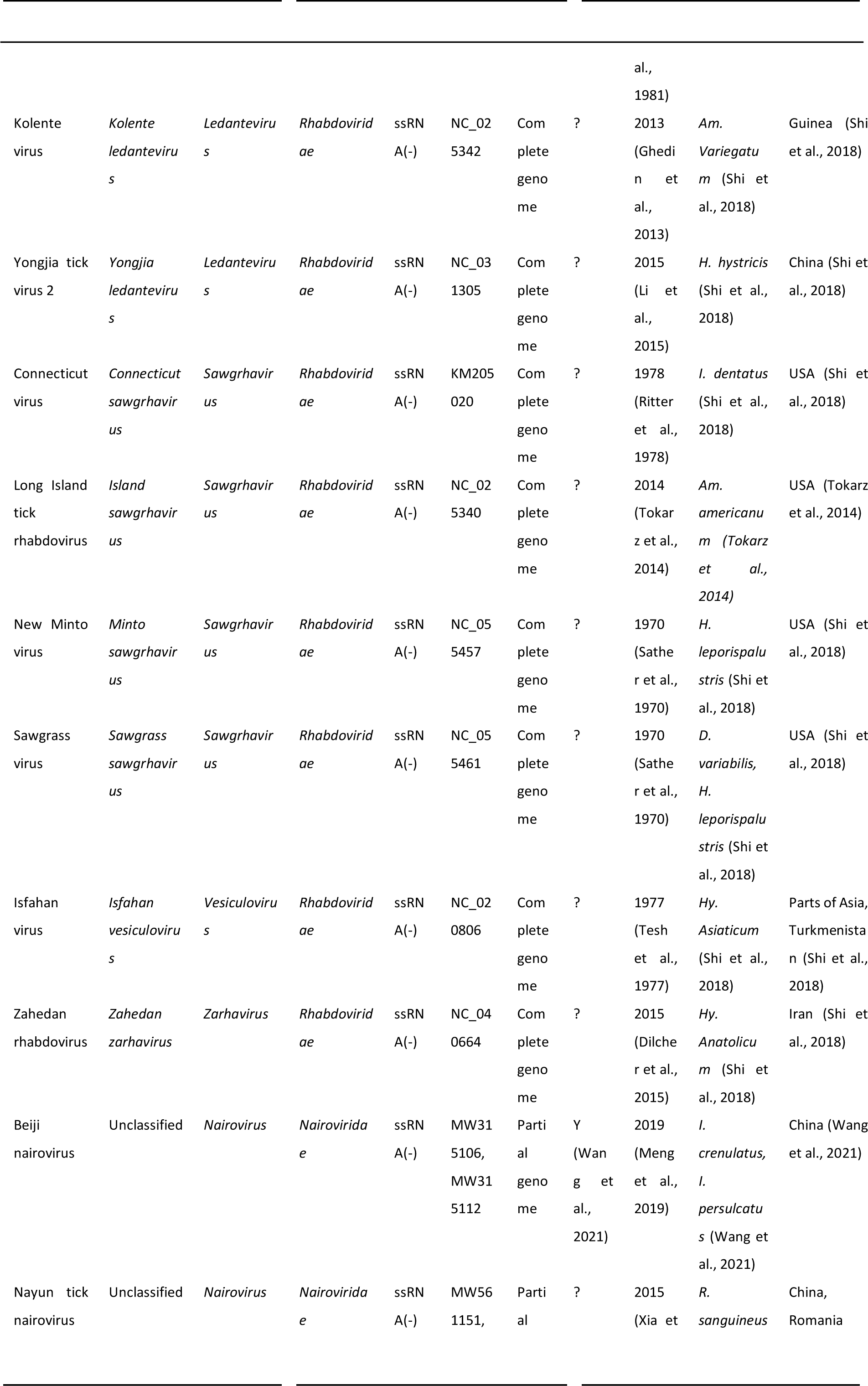
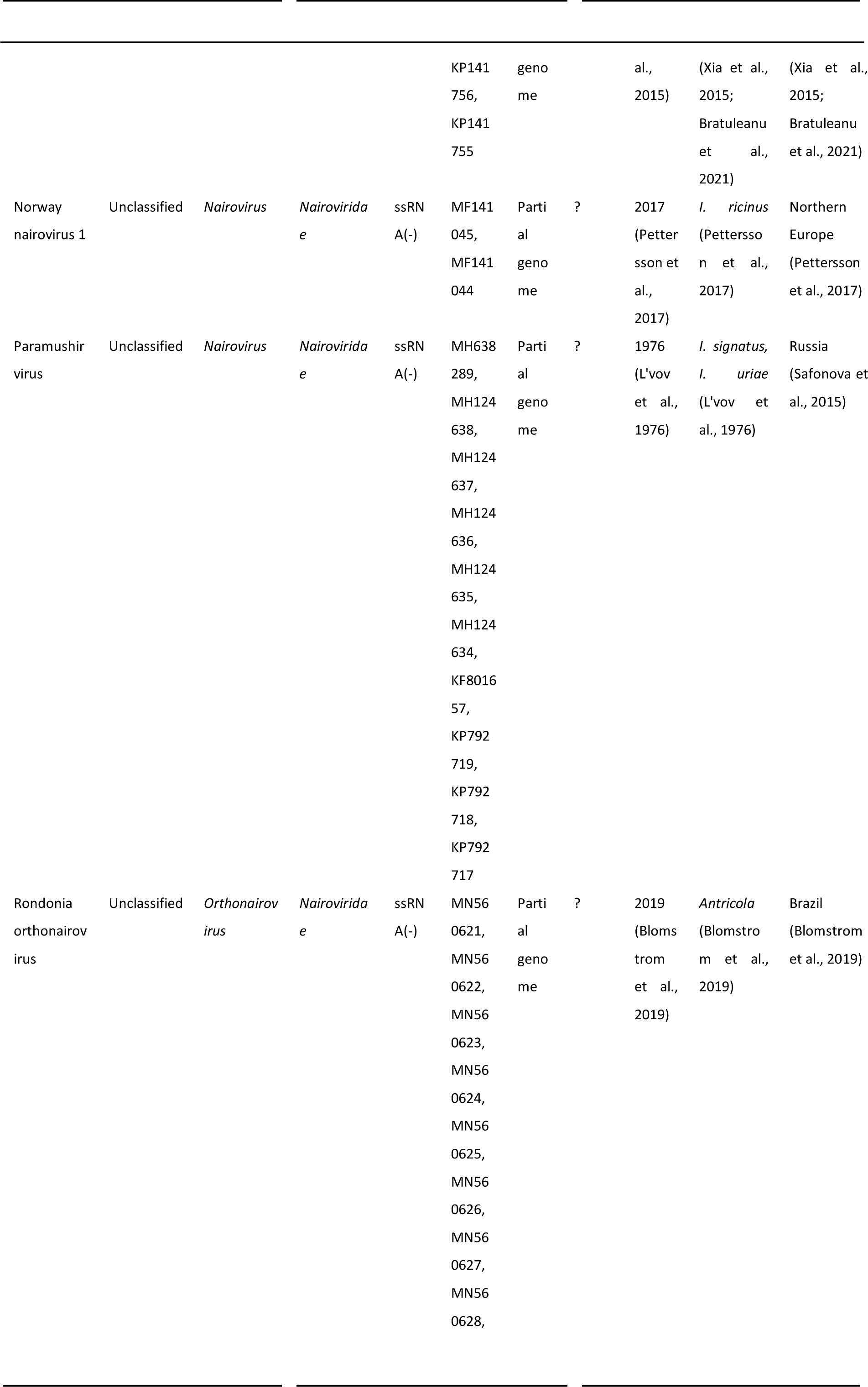
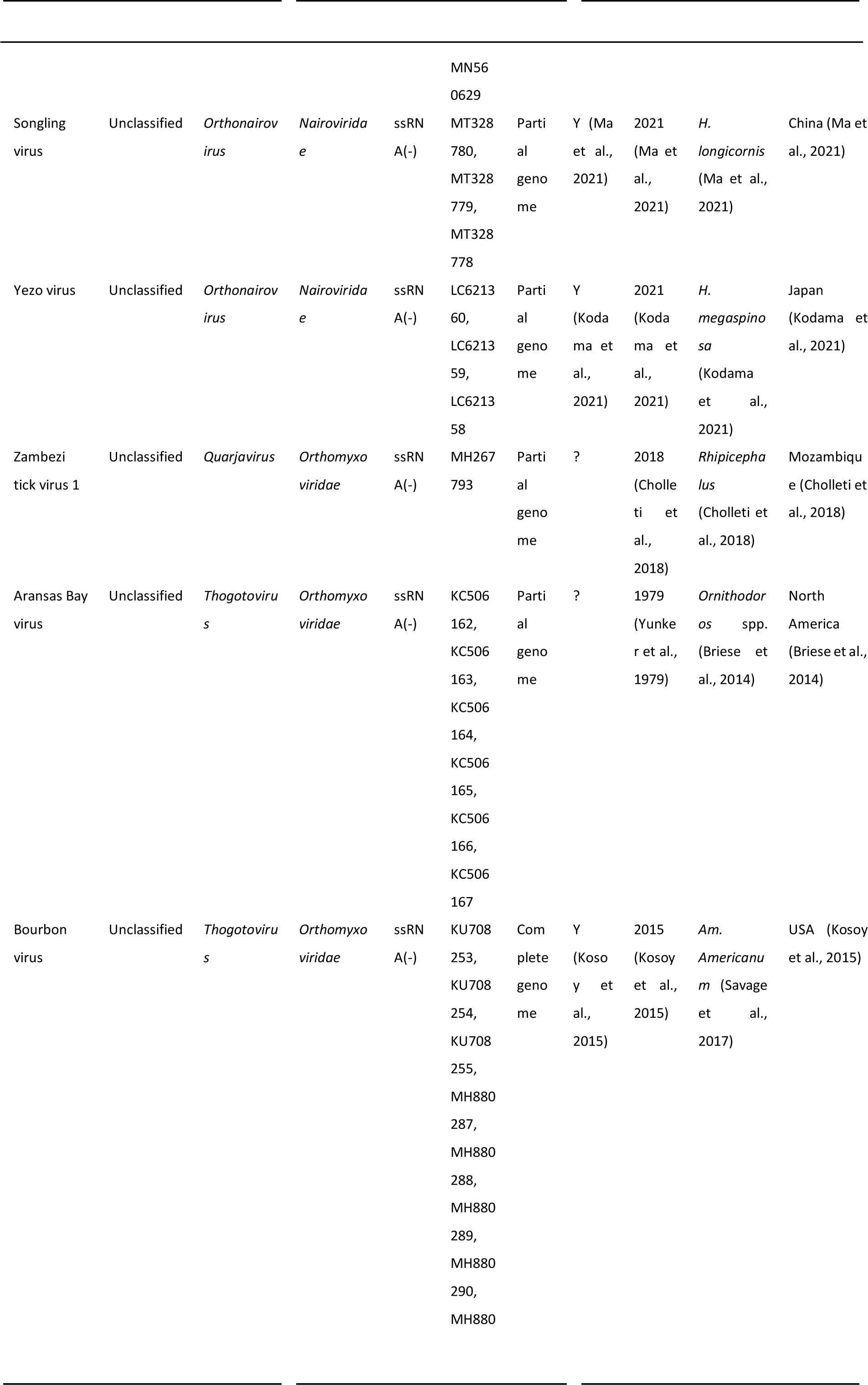
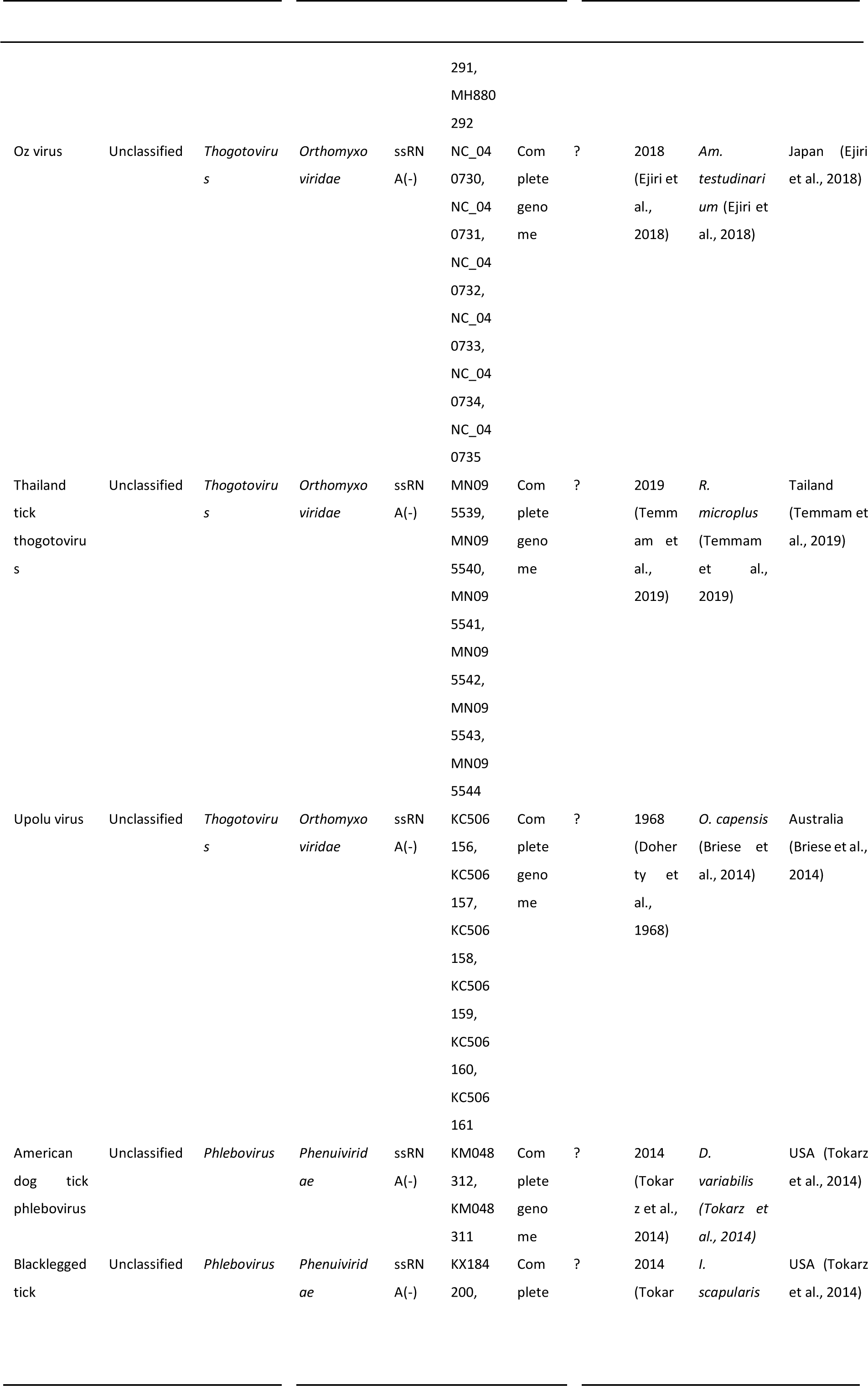
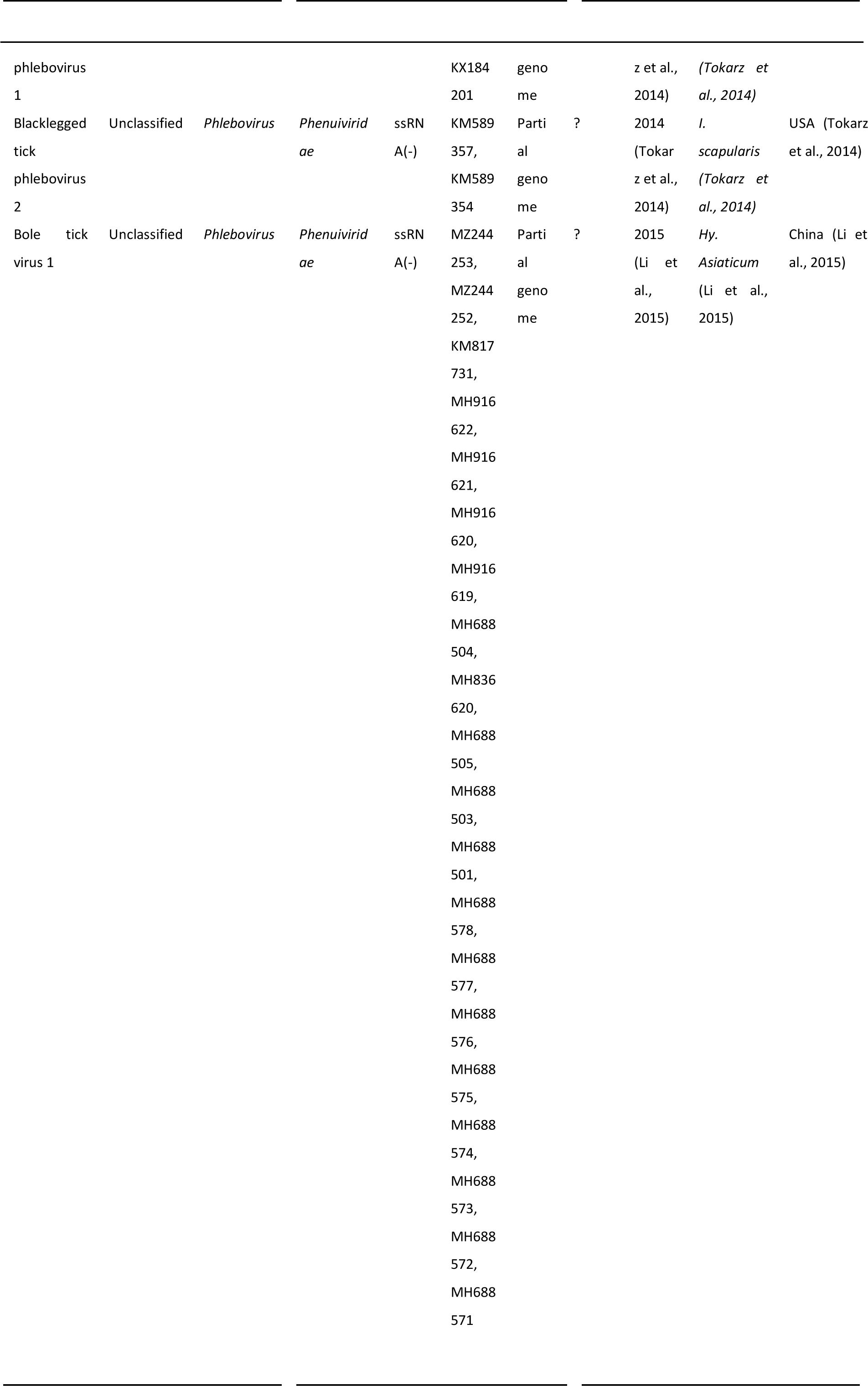
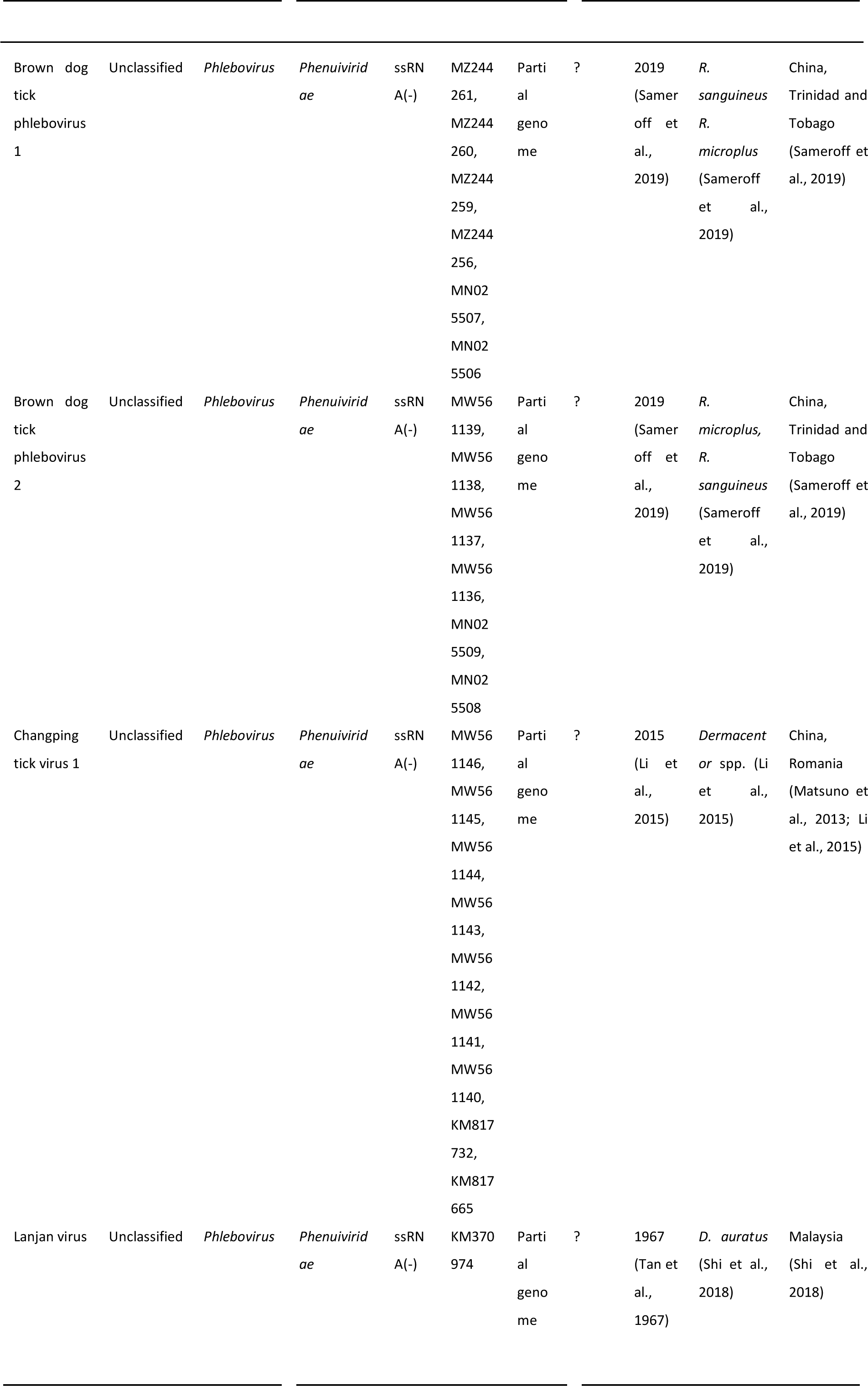
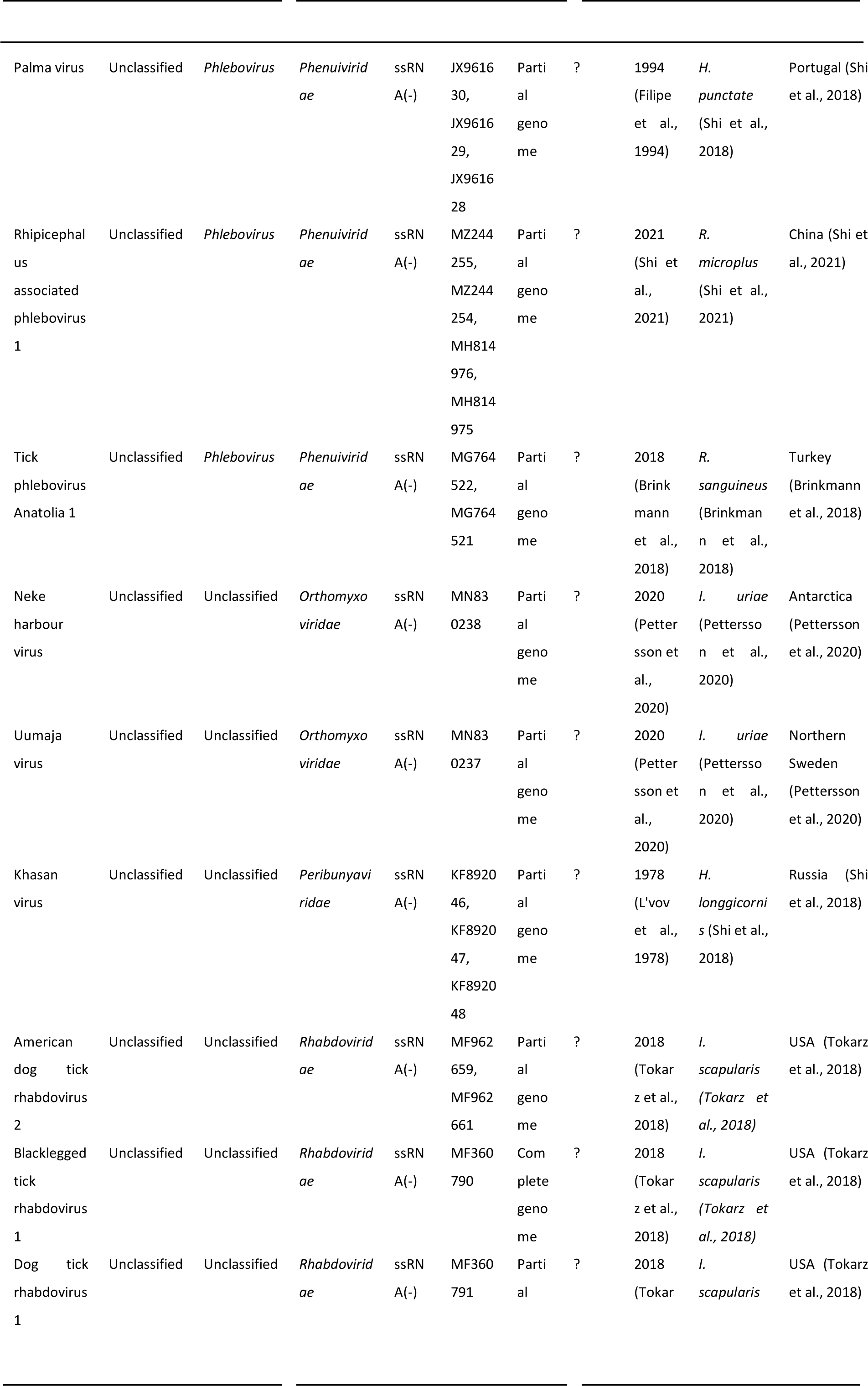
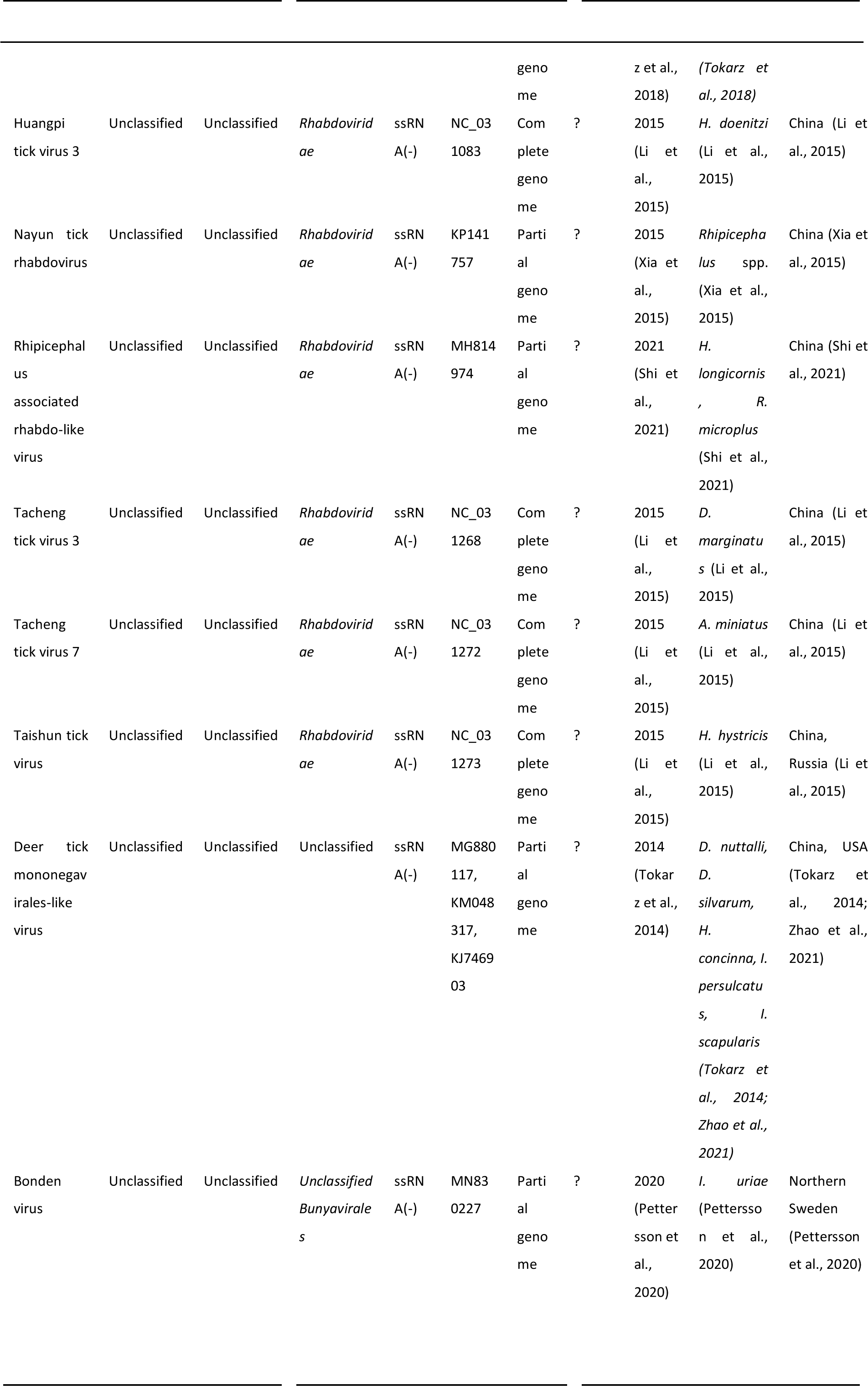
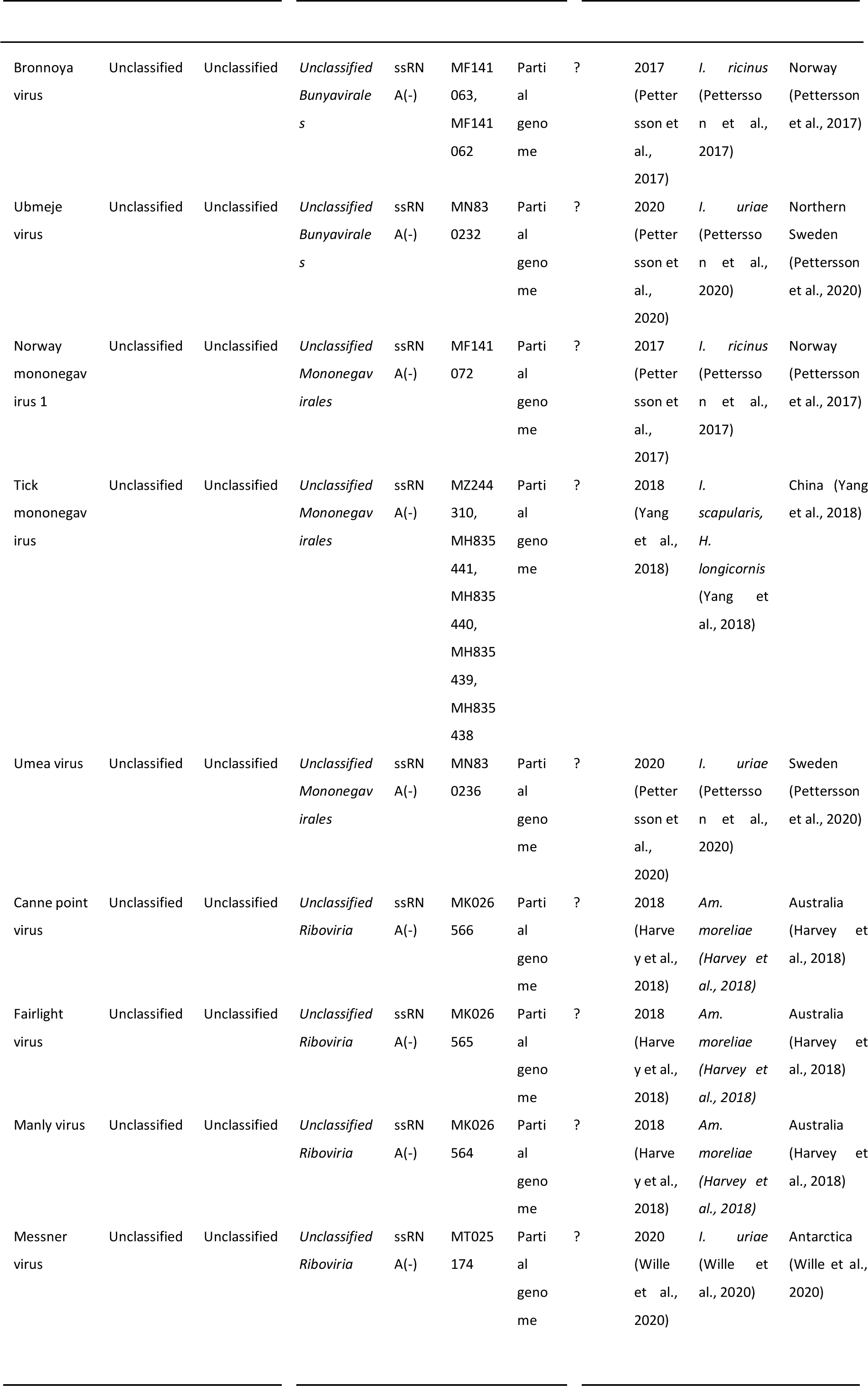
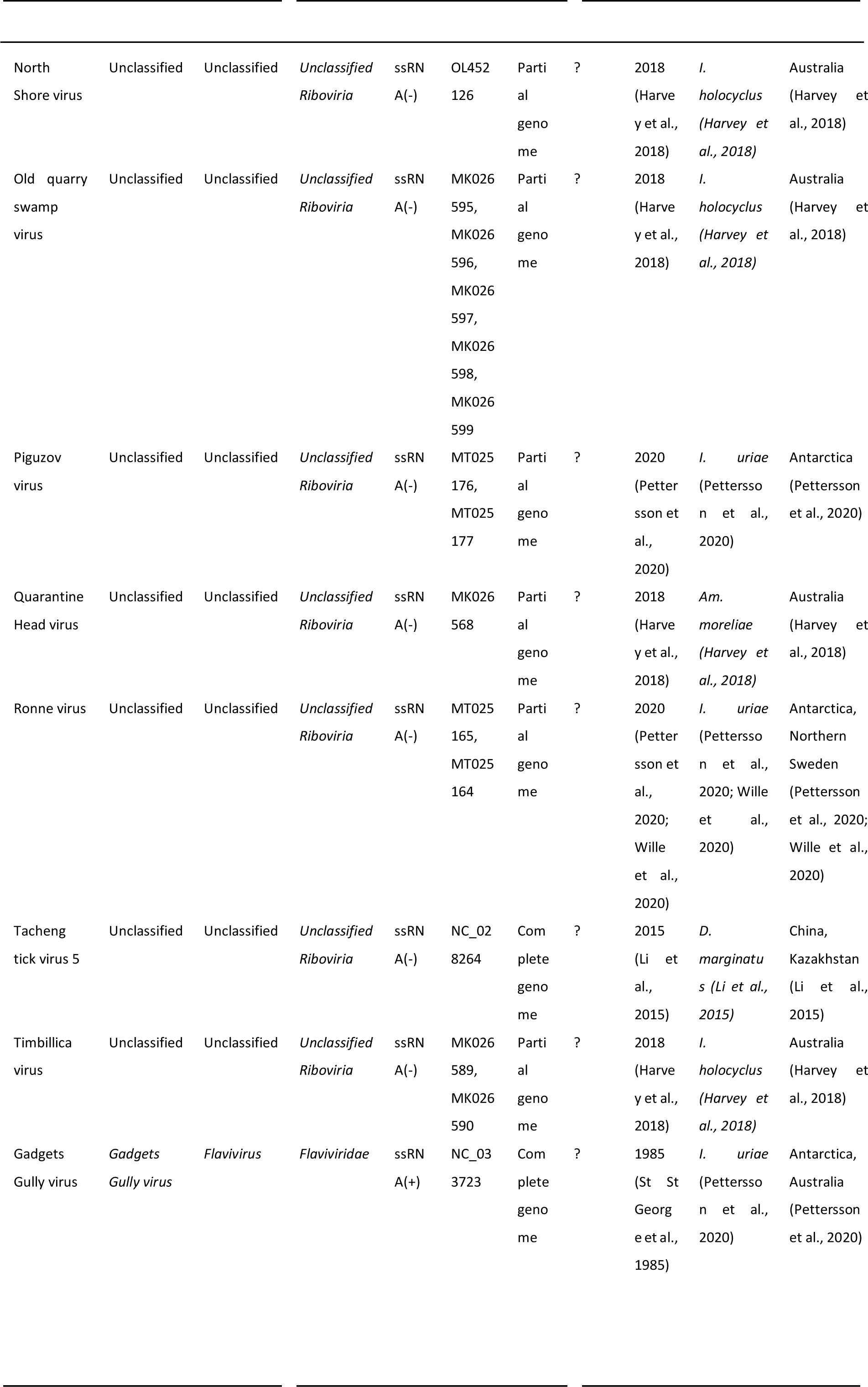
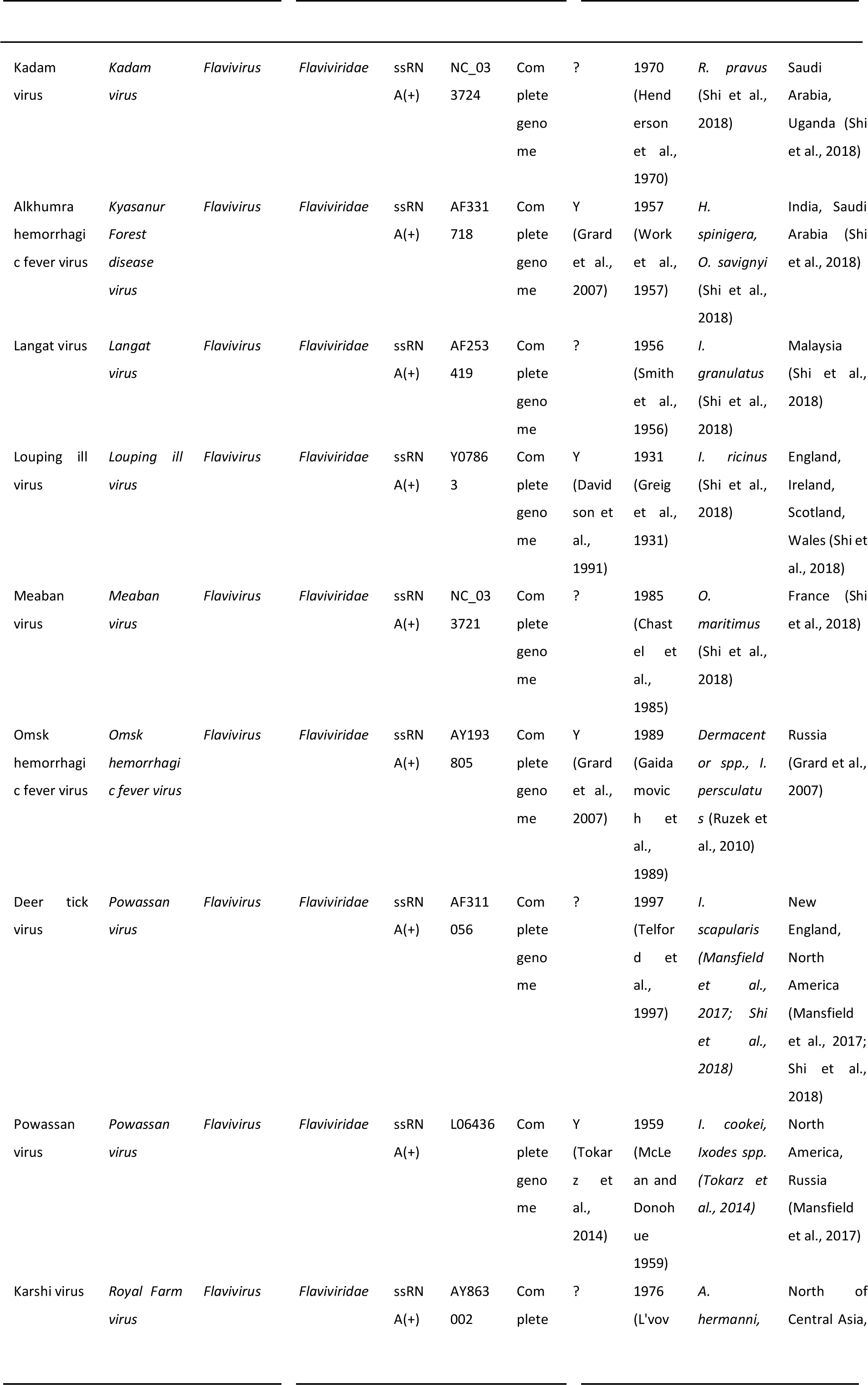
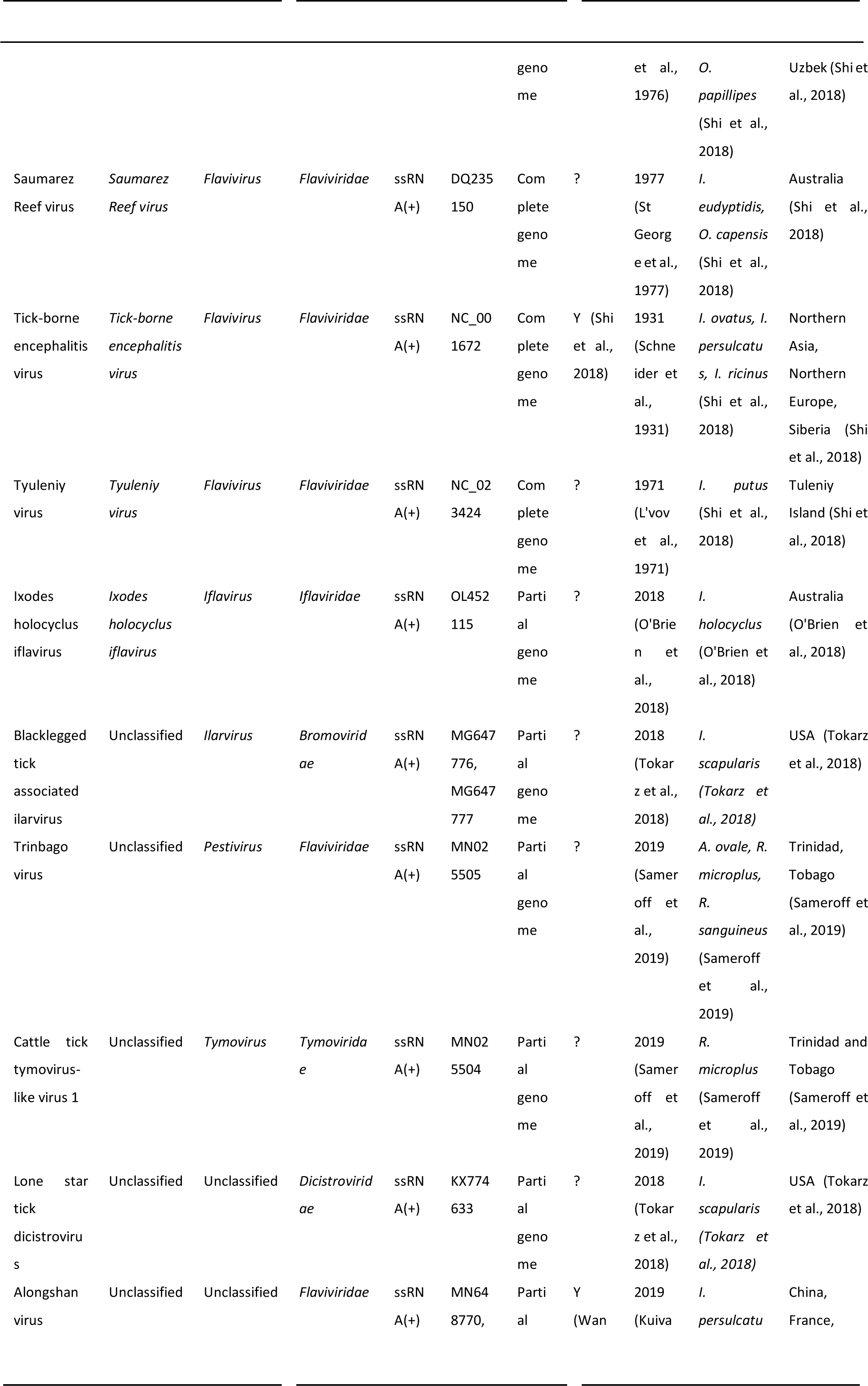
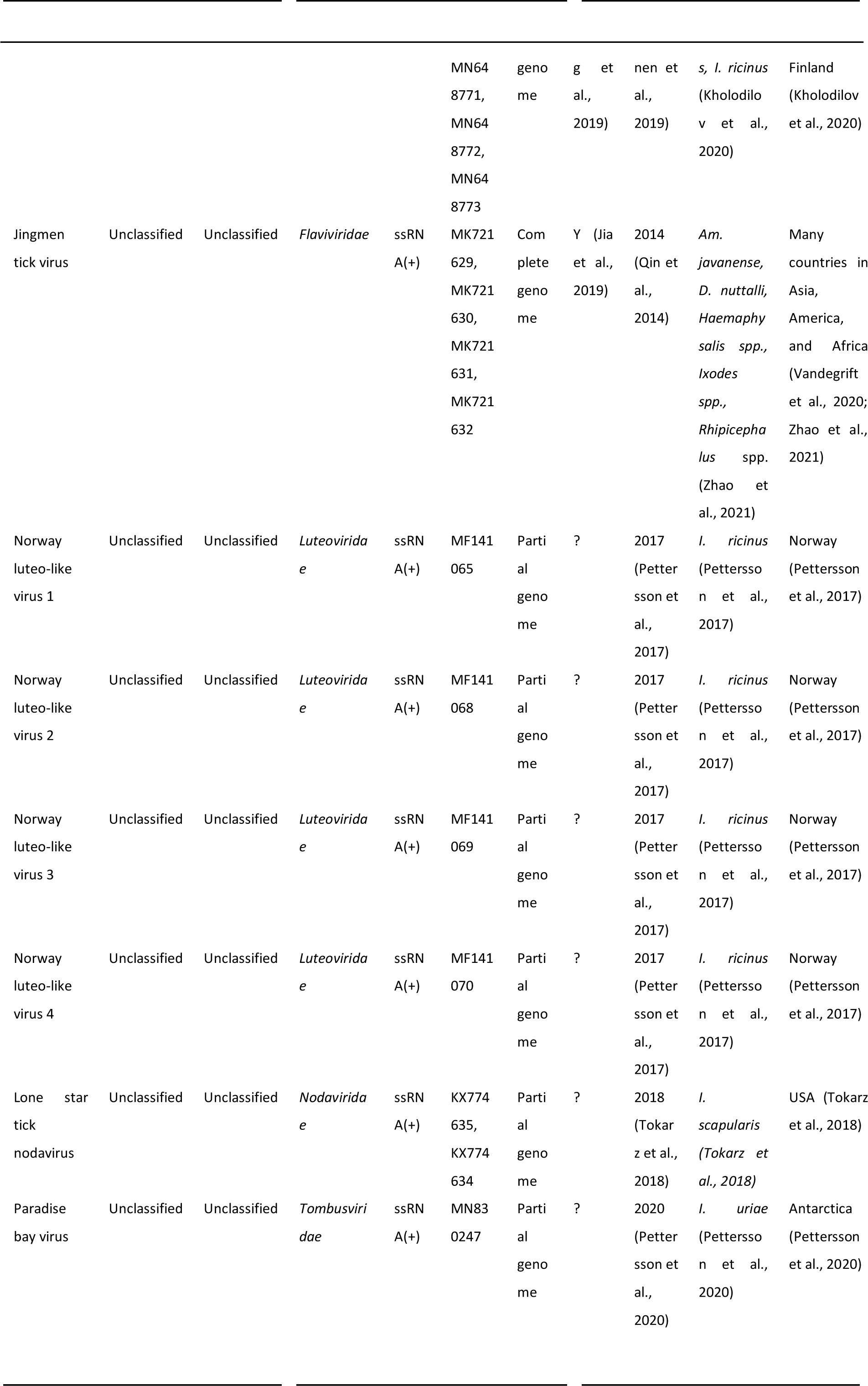
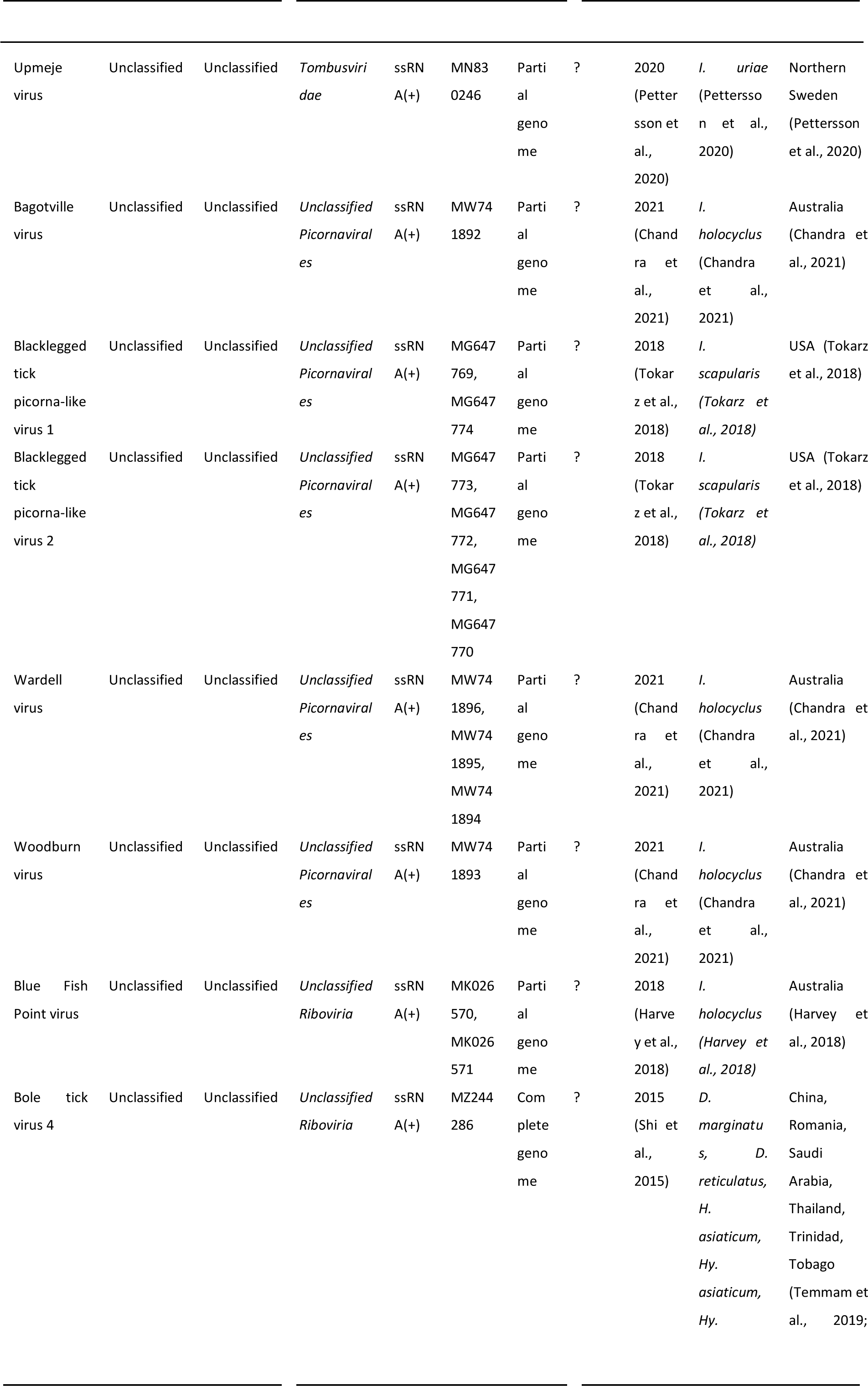
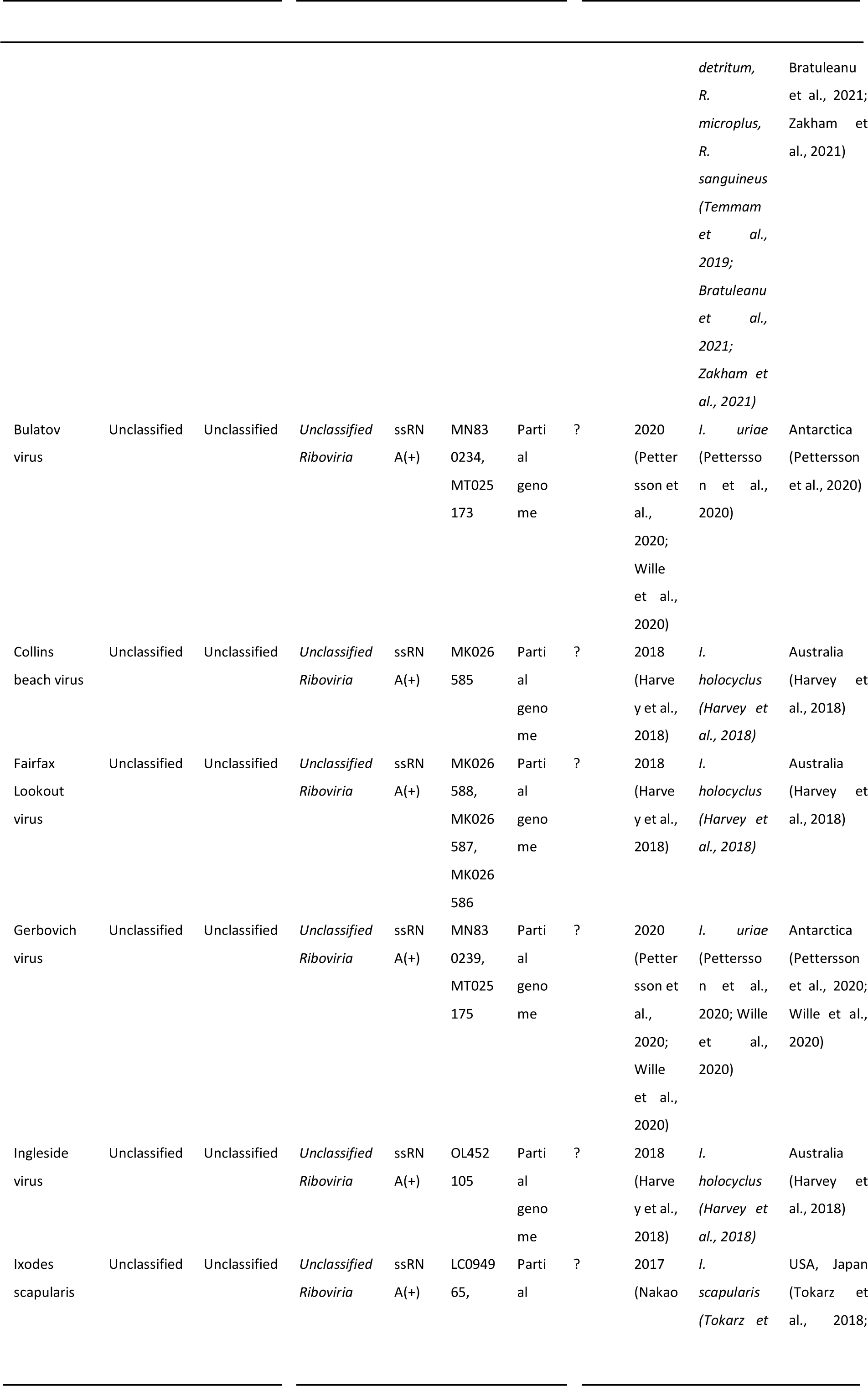
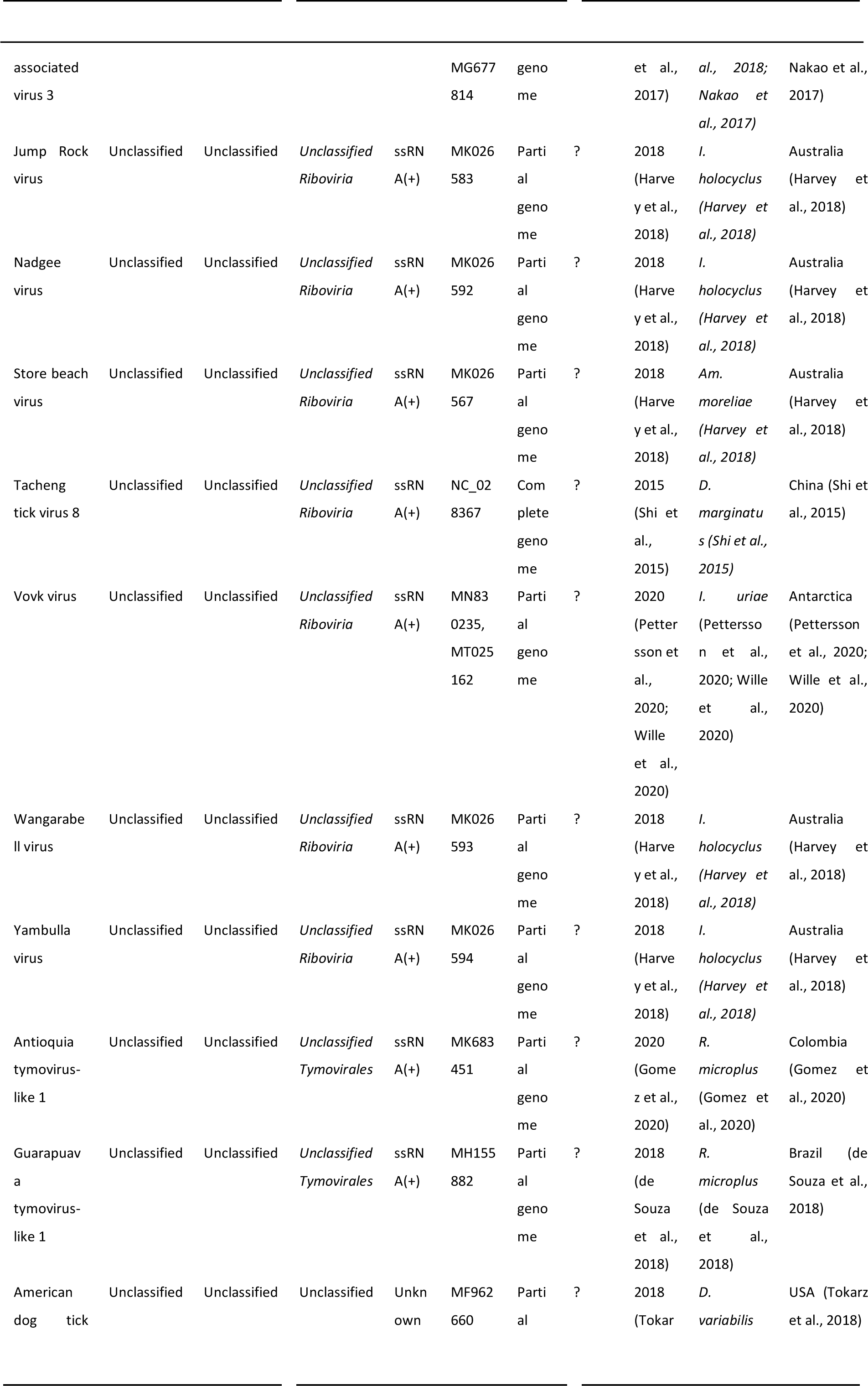
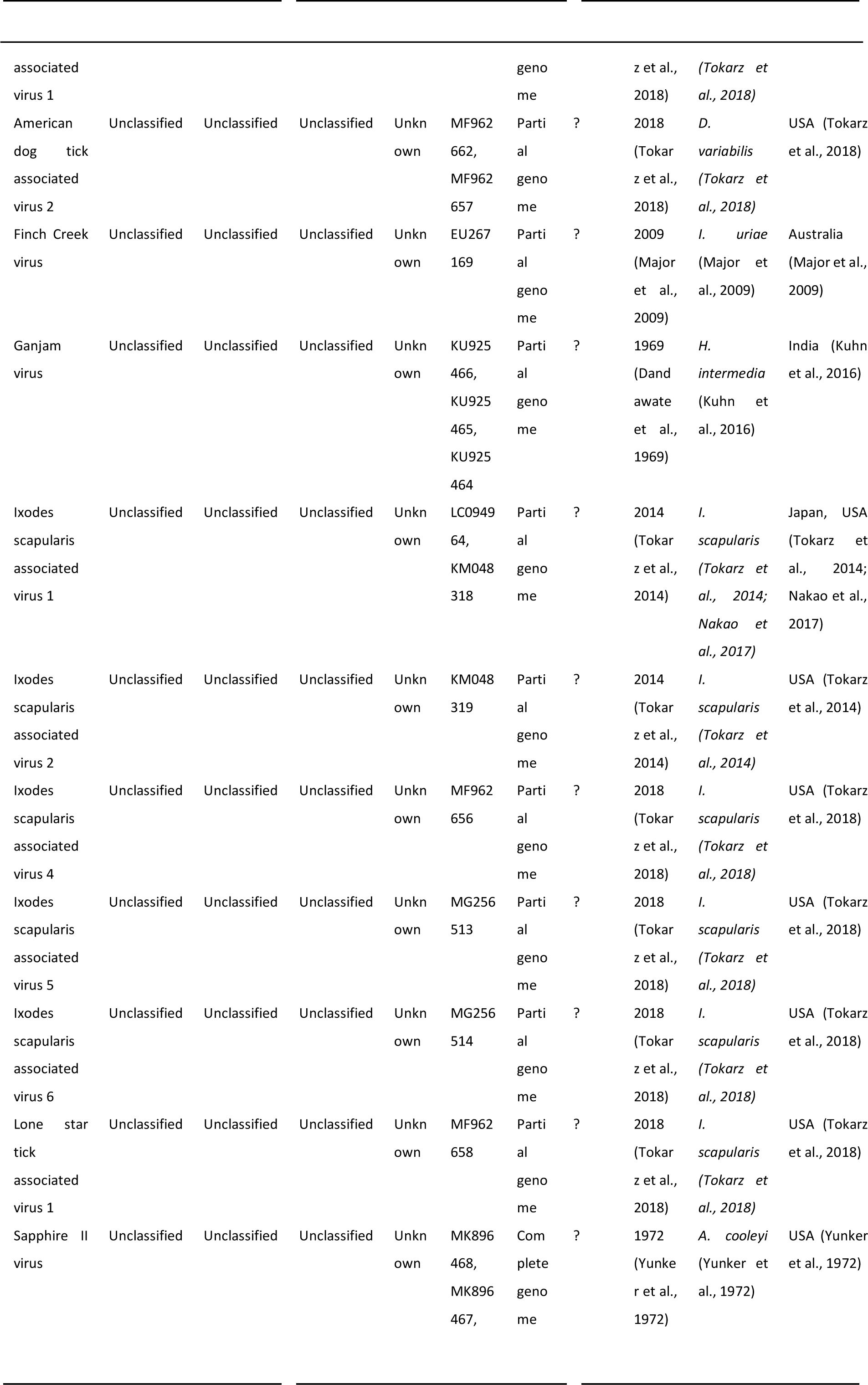
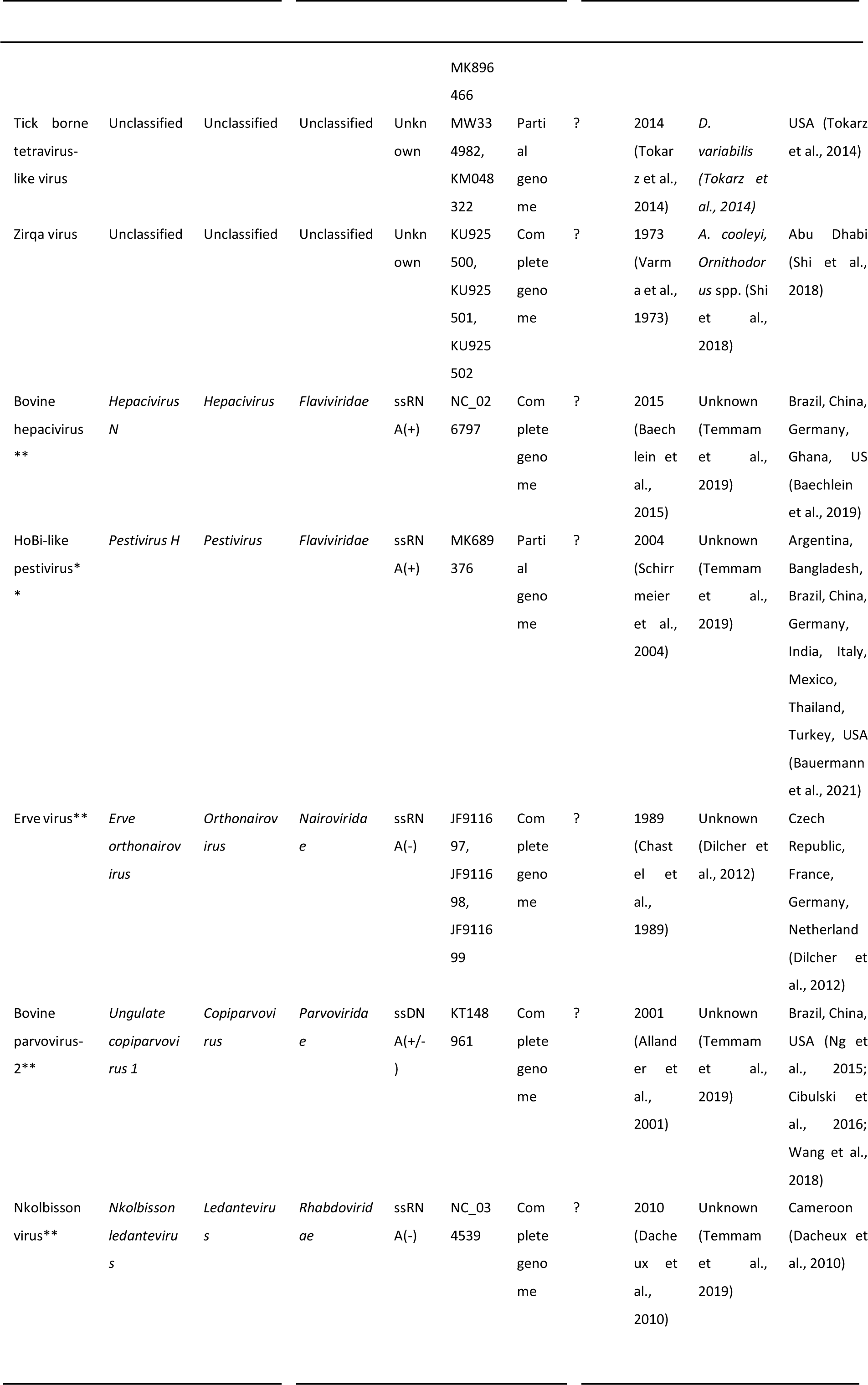
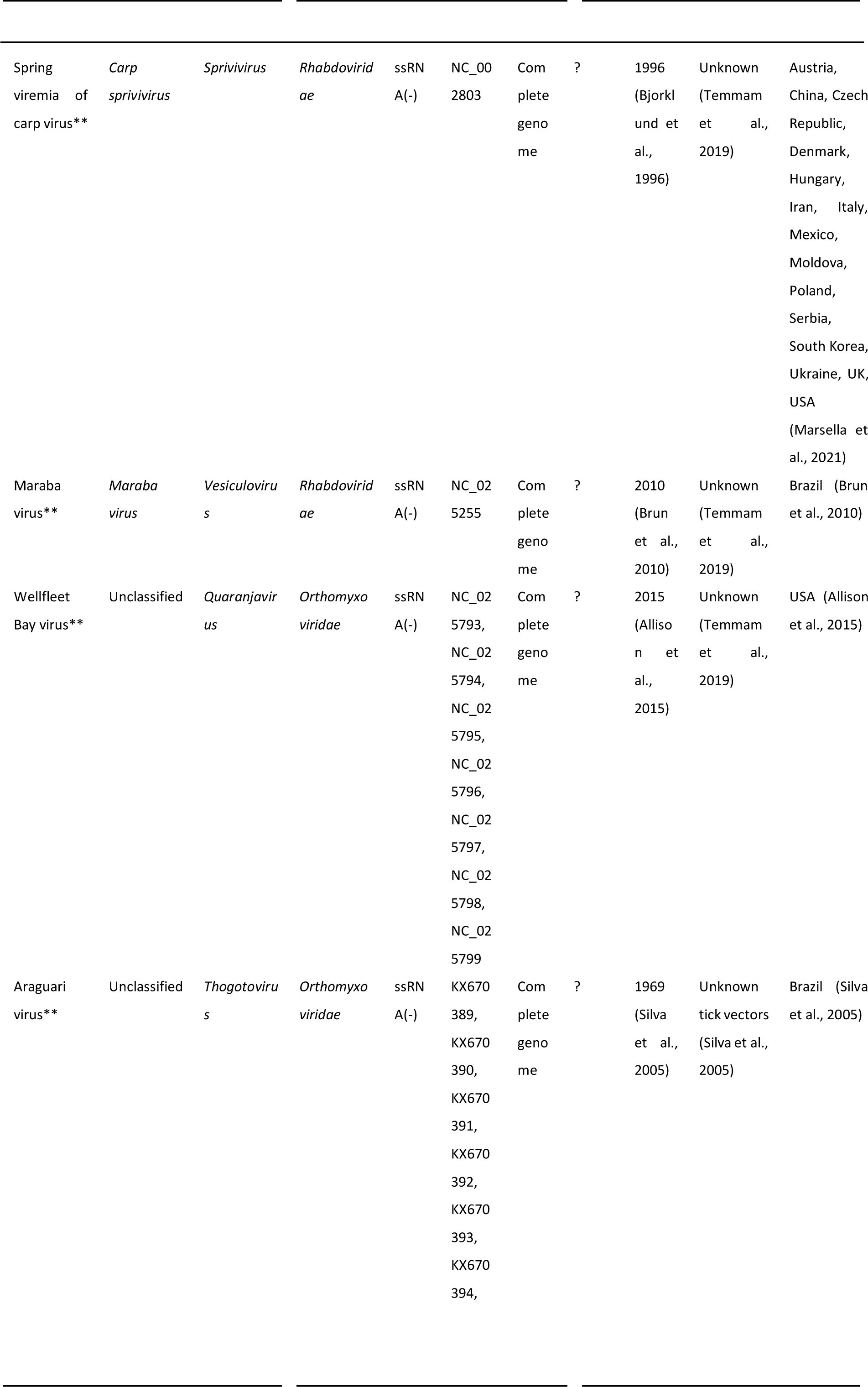
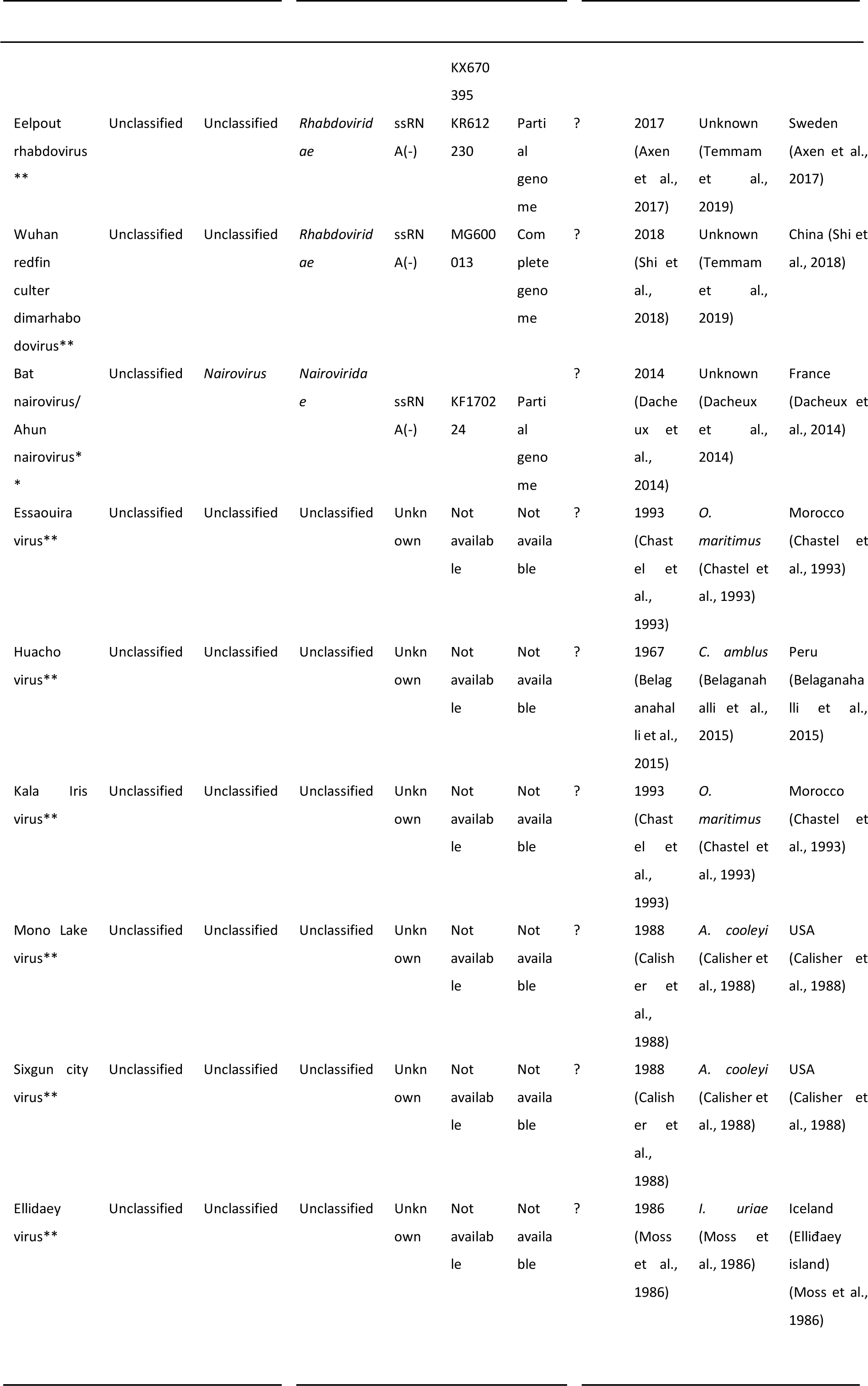
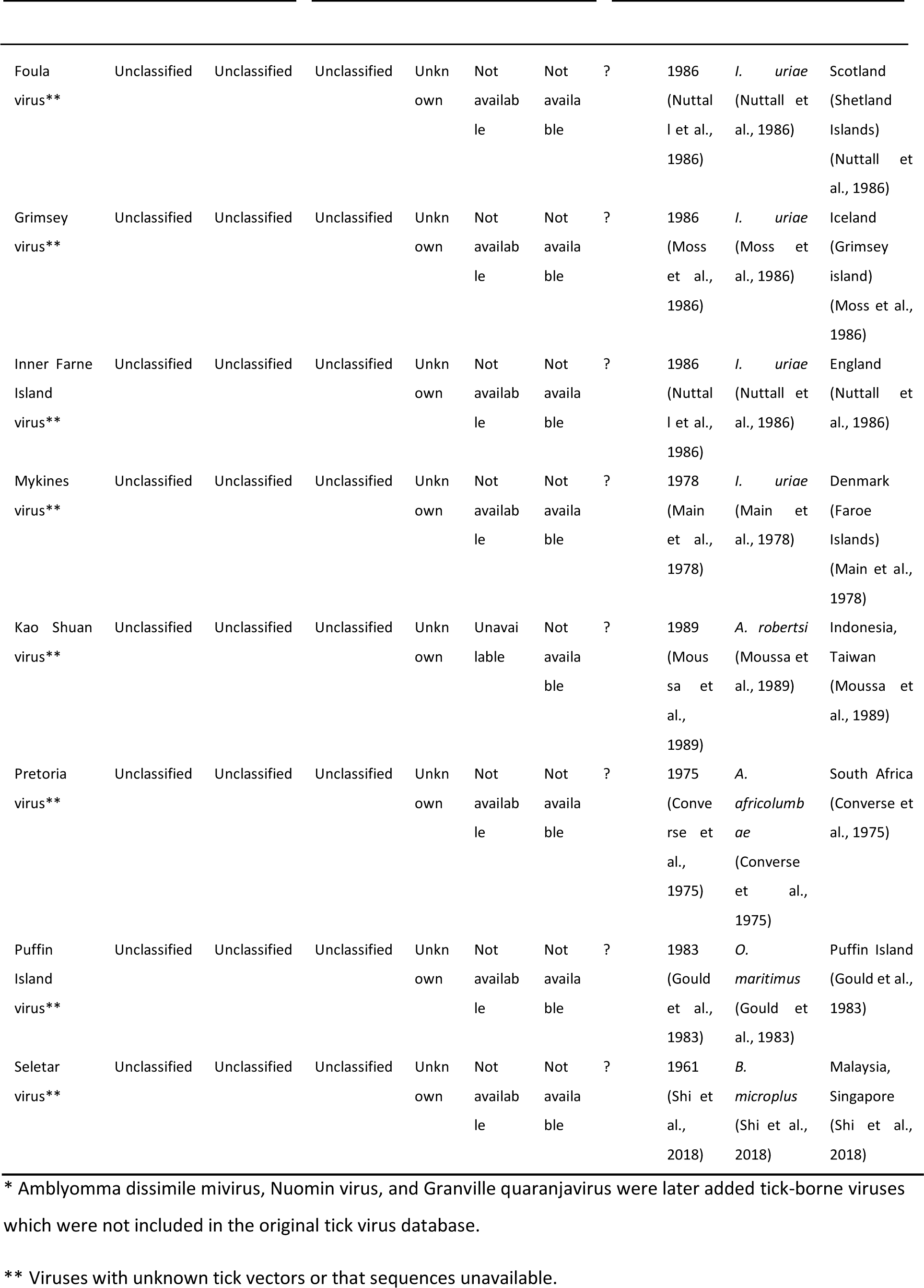
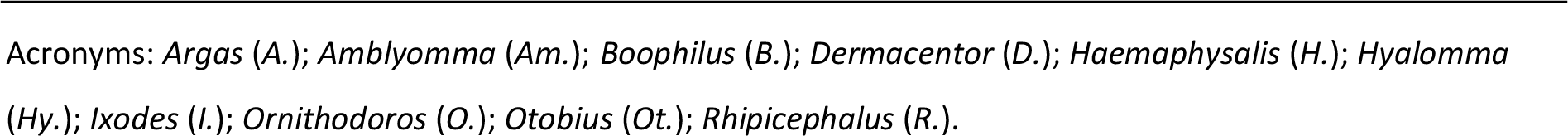
Summary of currently known tick-borne viruses by December 2021. Information from a wide variety of sources, references in the table.

**Appendix Table 2.**
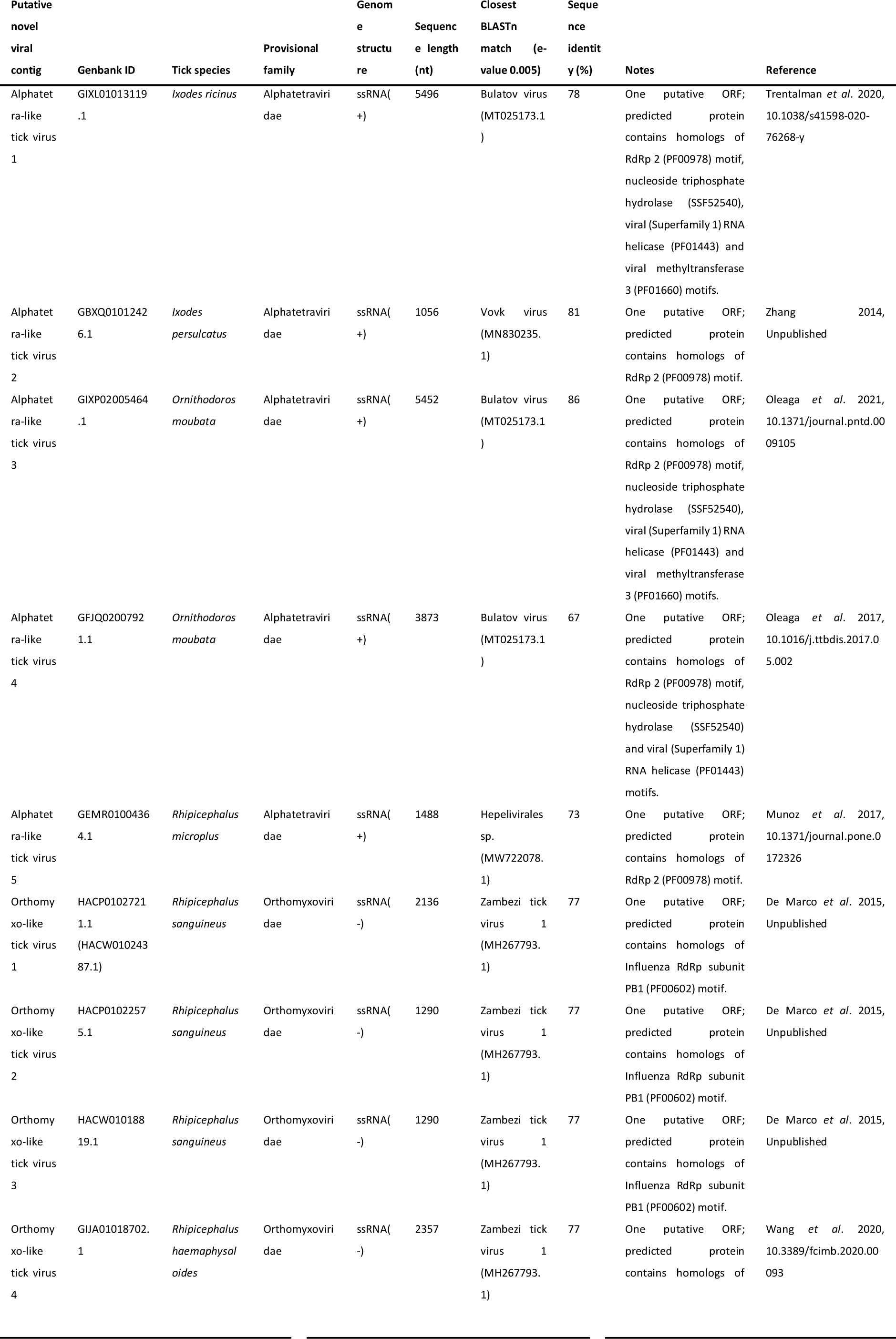
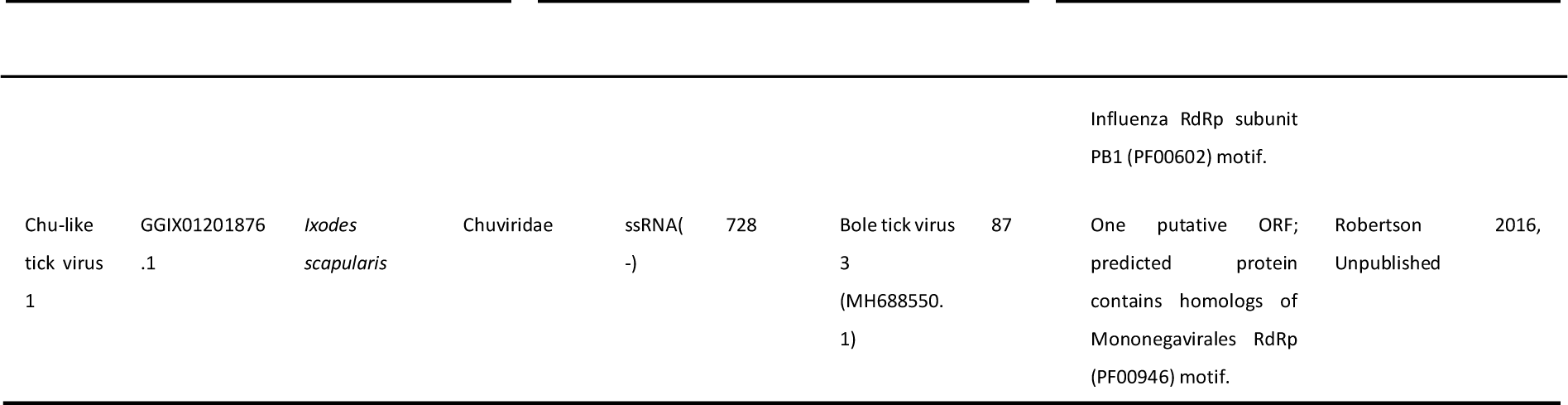
Putative novel RNV viral contigs identified in this study.

**Appendix Table 3.**
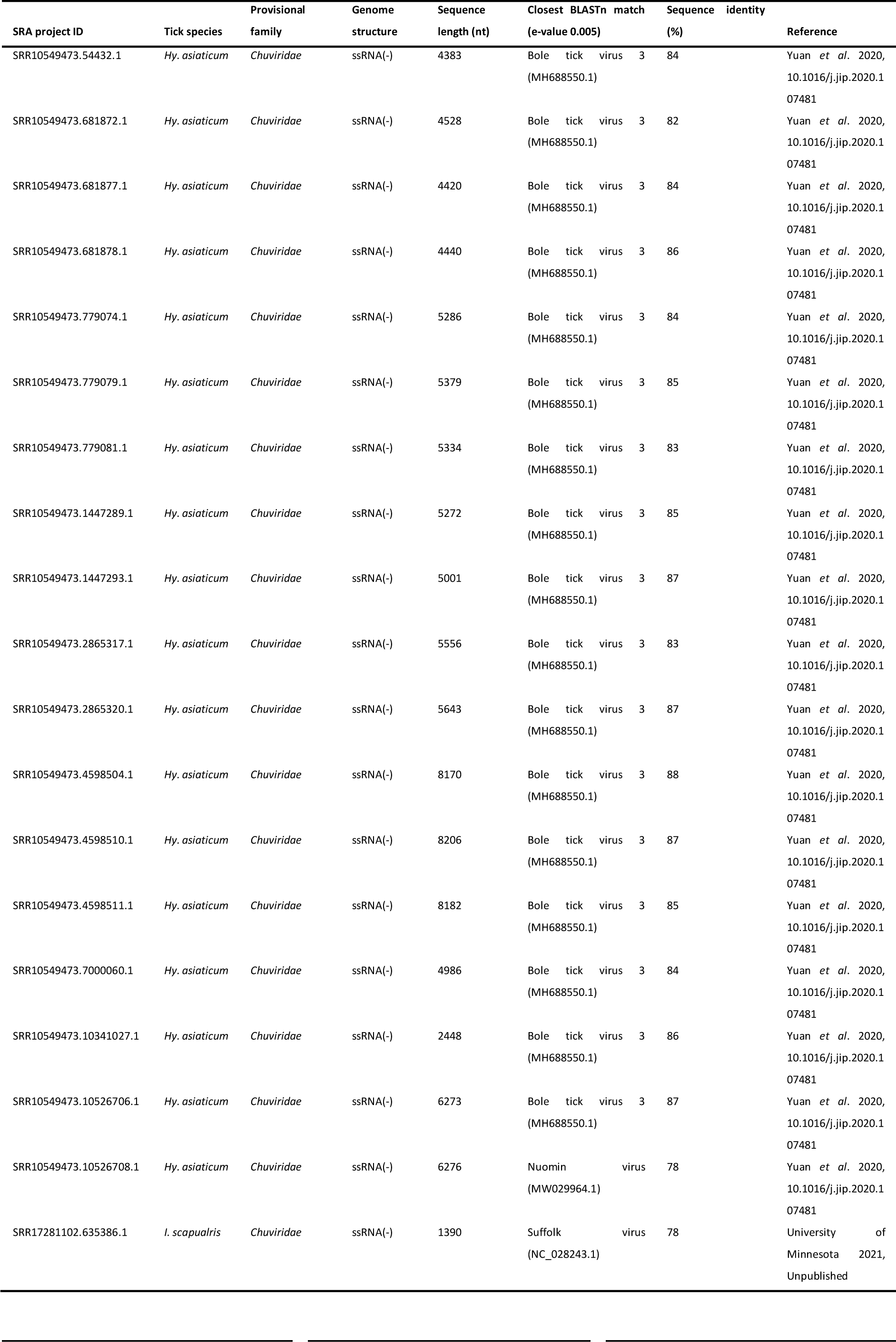
Putative novel RNA viral contigs that represent defective viruses.

**Appendix Table 4.**
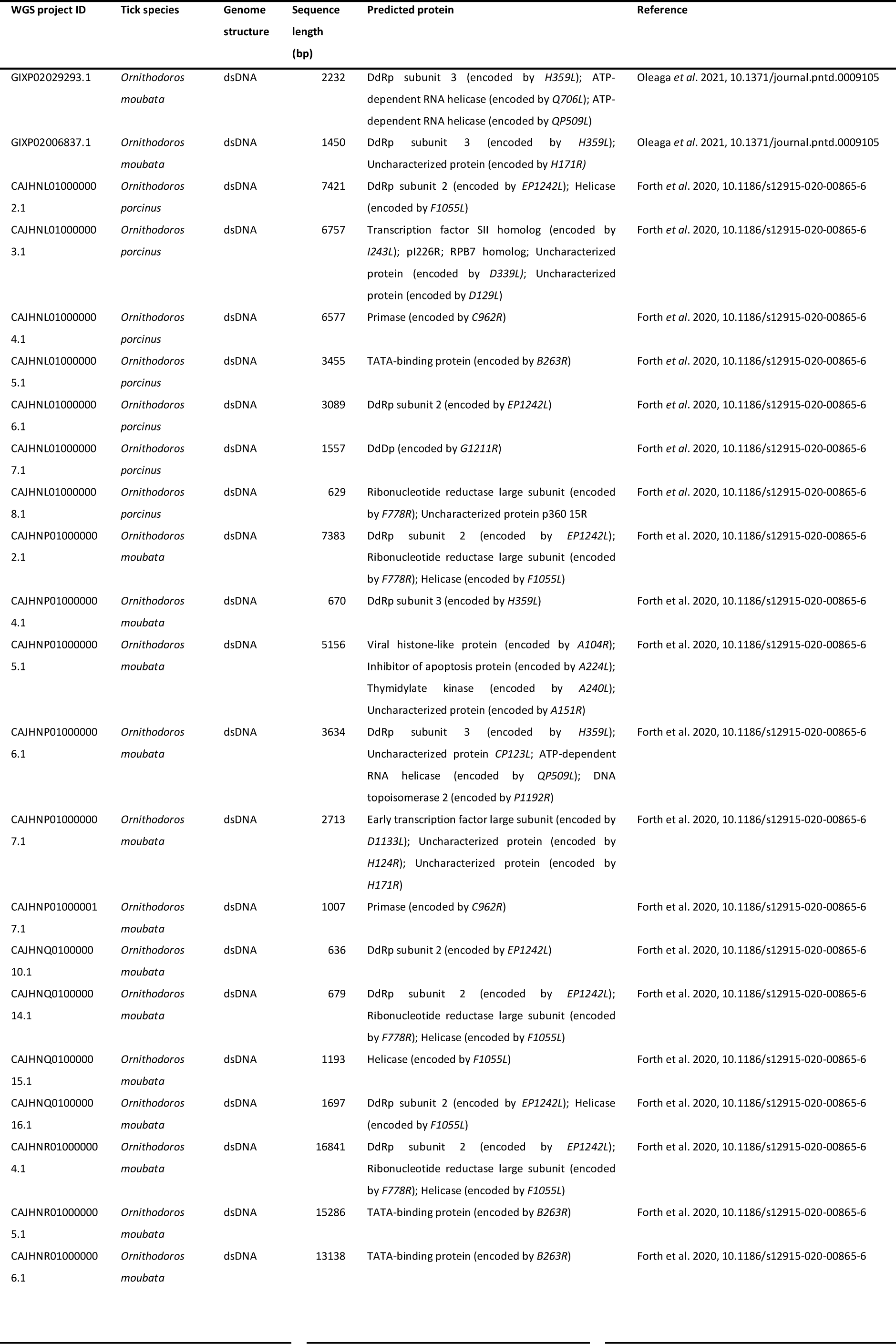
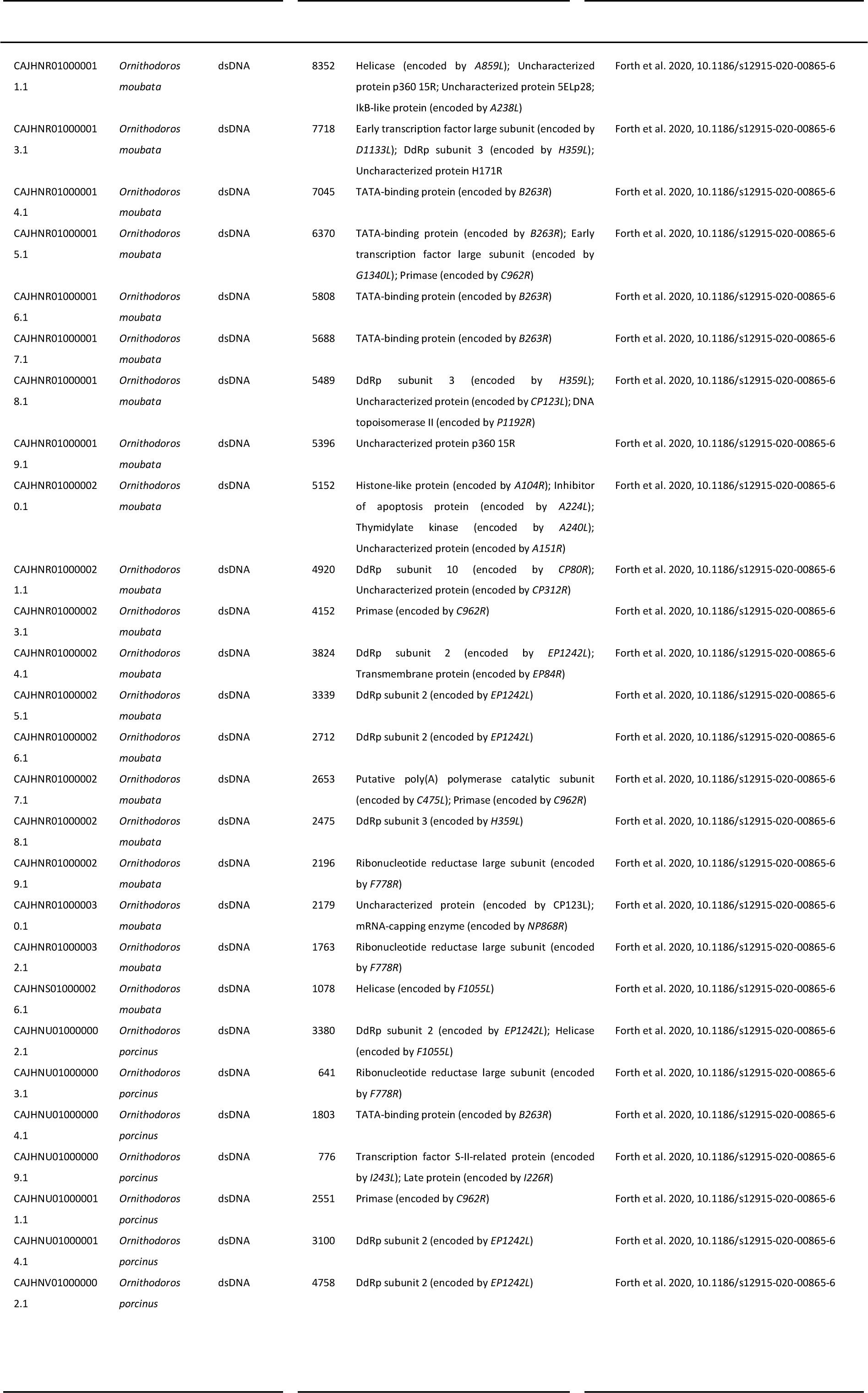
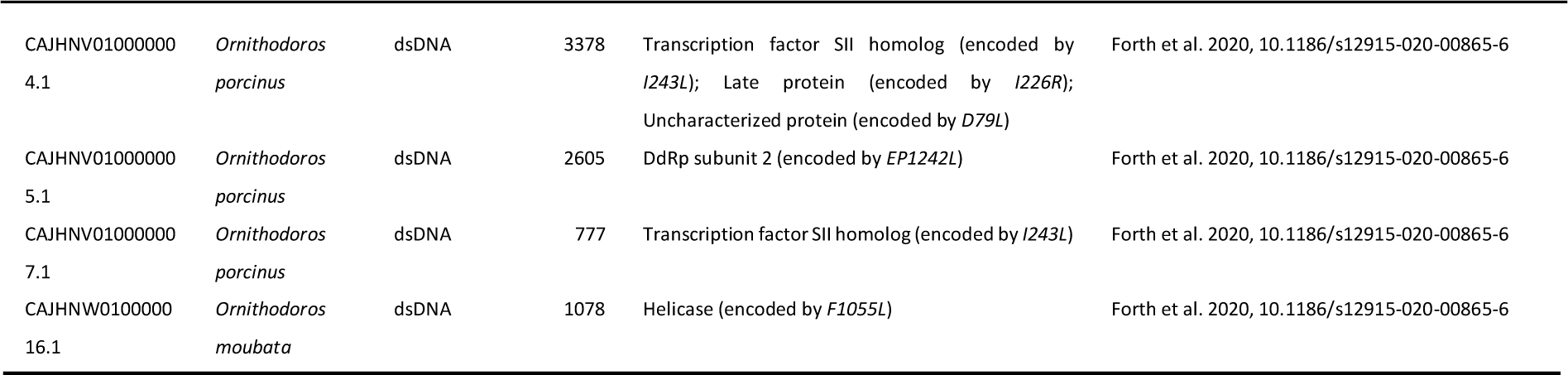
DNA viral contigs found after bioinformatic checking.

**Appendix Table 5.**
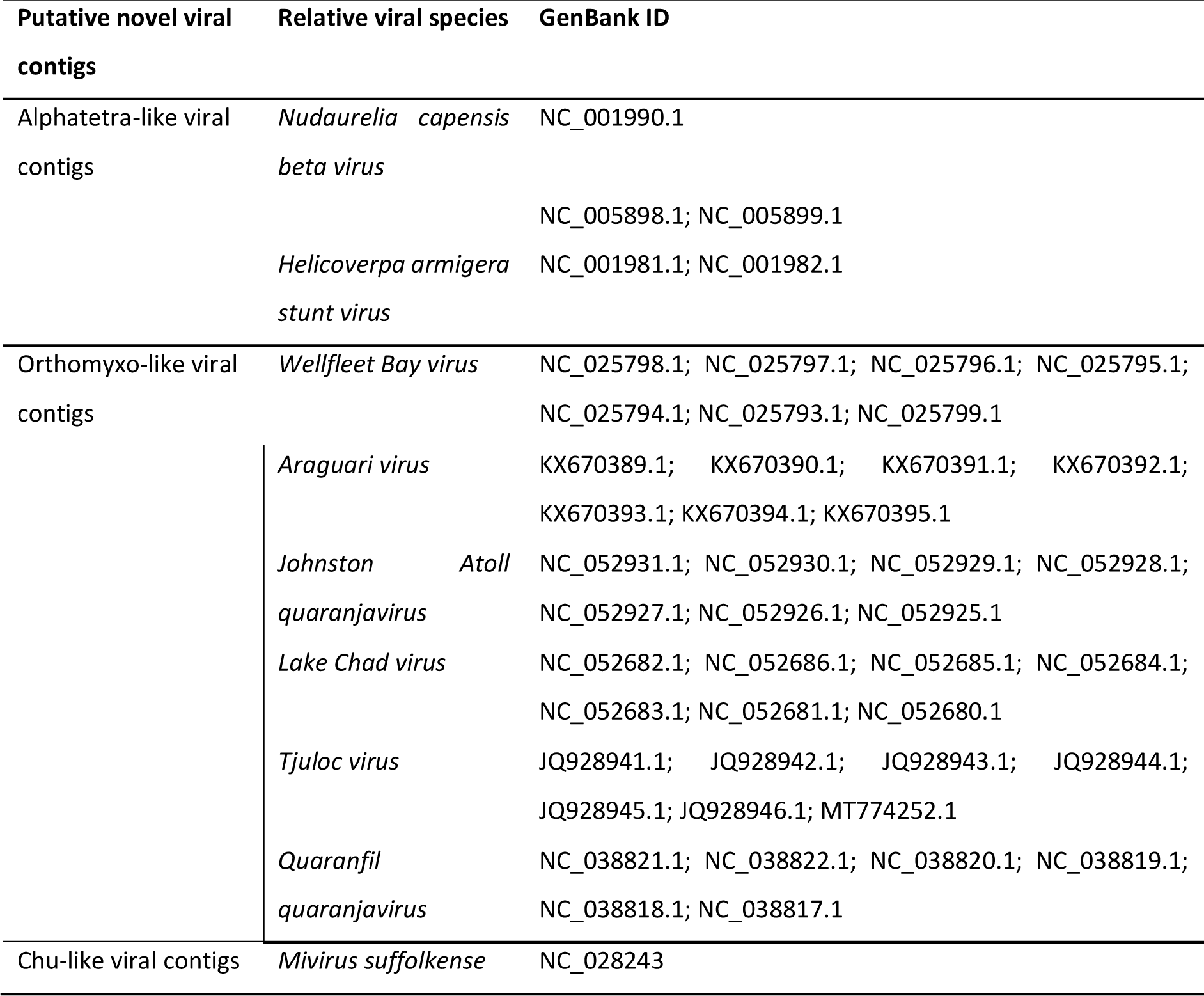
GenBank ID of viruses phylogenetically related to the putative novel viral contigs. Zoonotic risks of virus sequences (in GenBank format) were predicted using the GCB model.

**Appendix Table 6.**
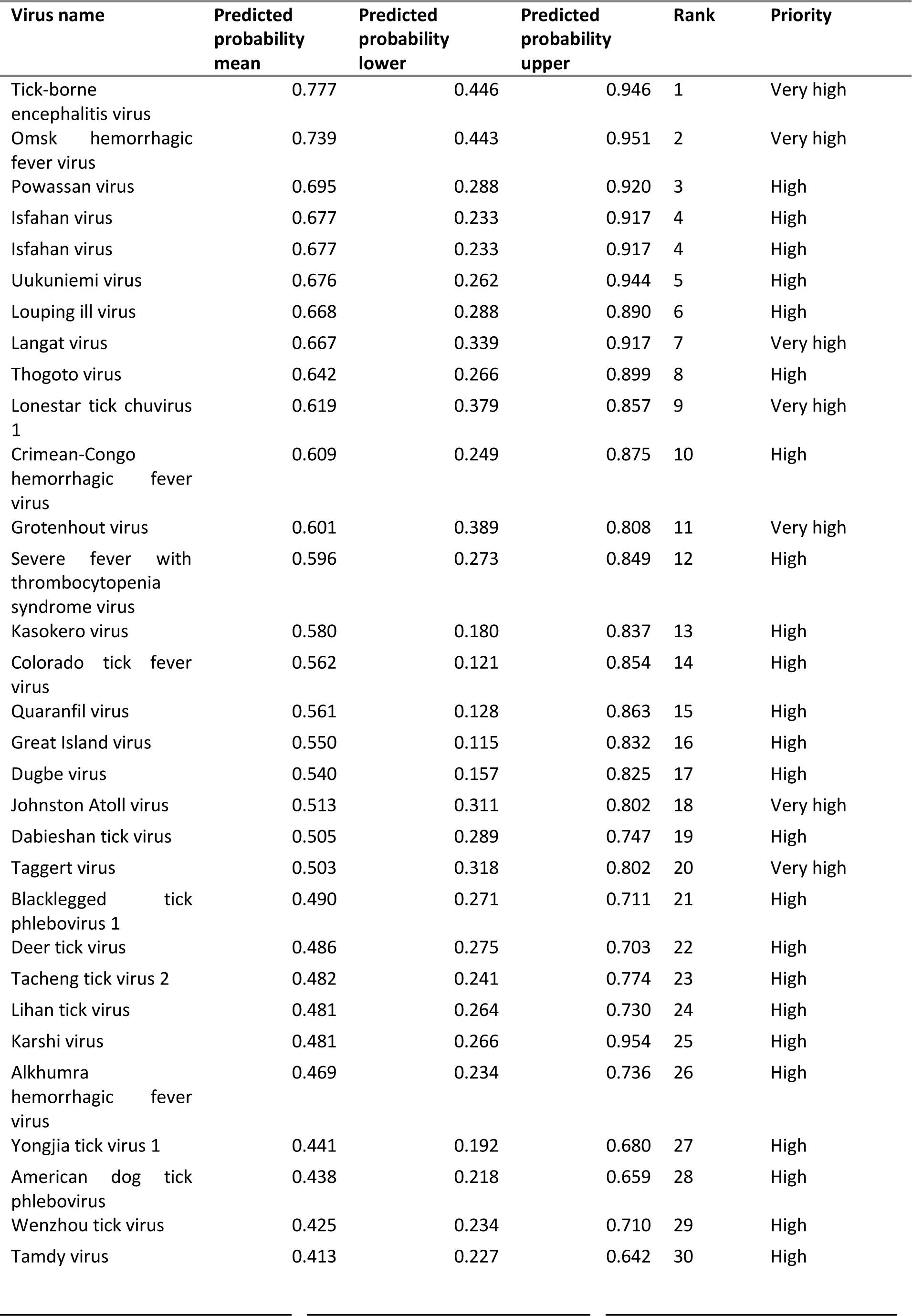
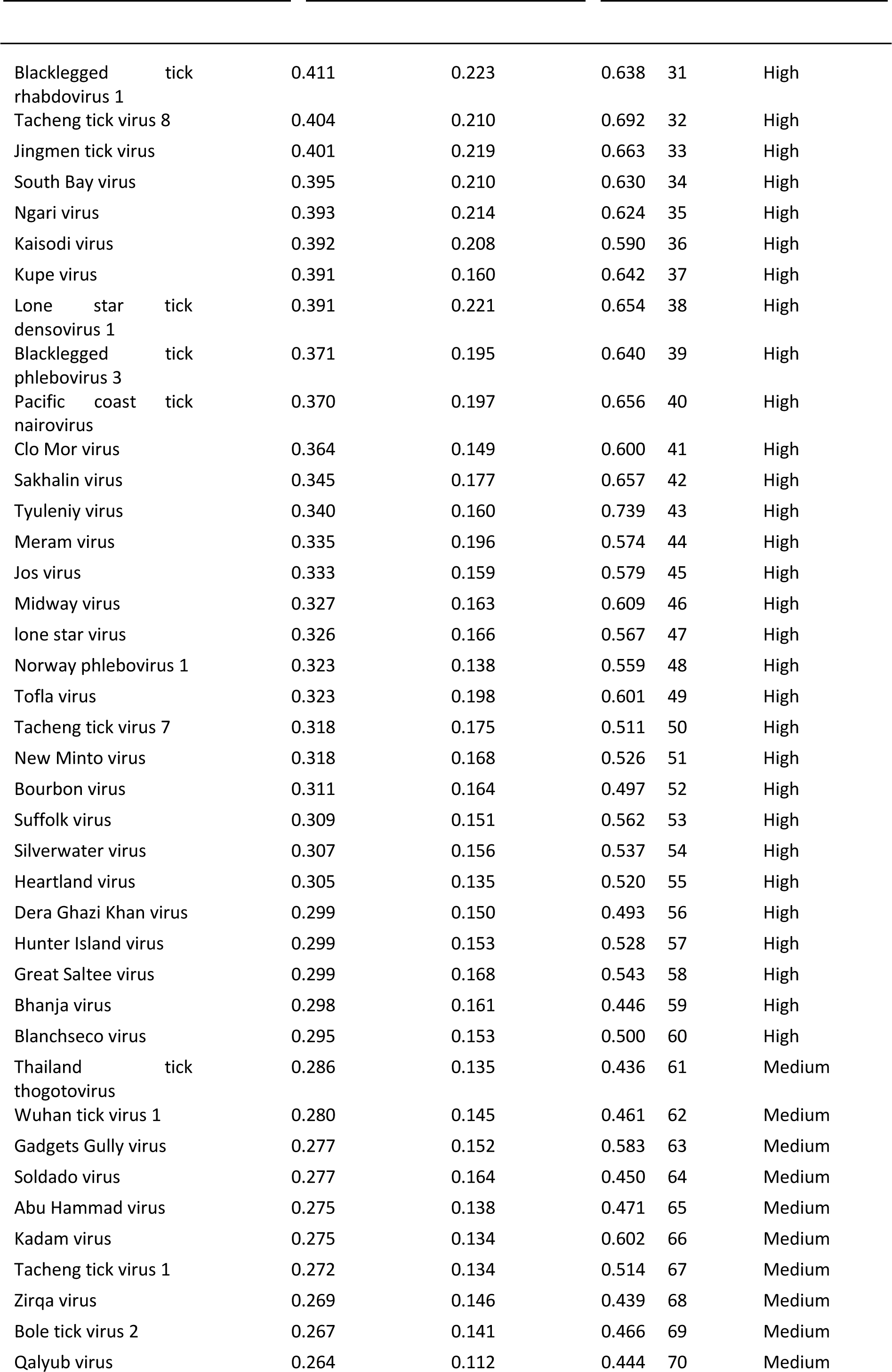
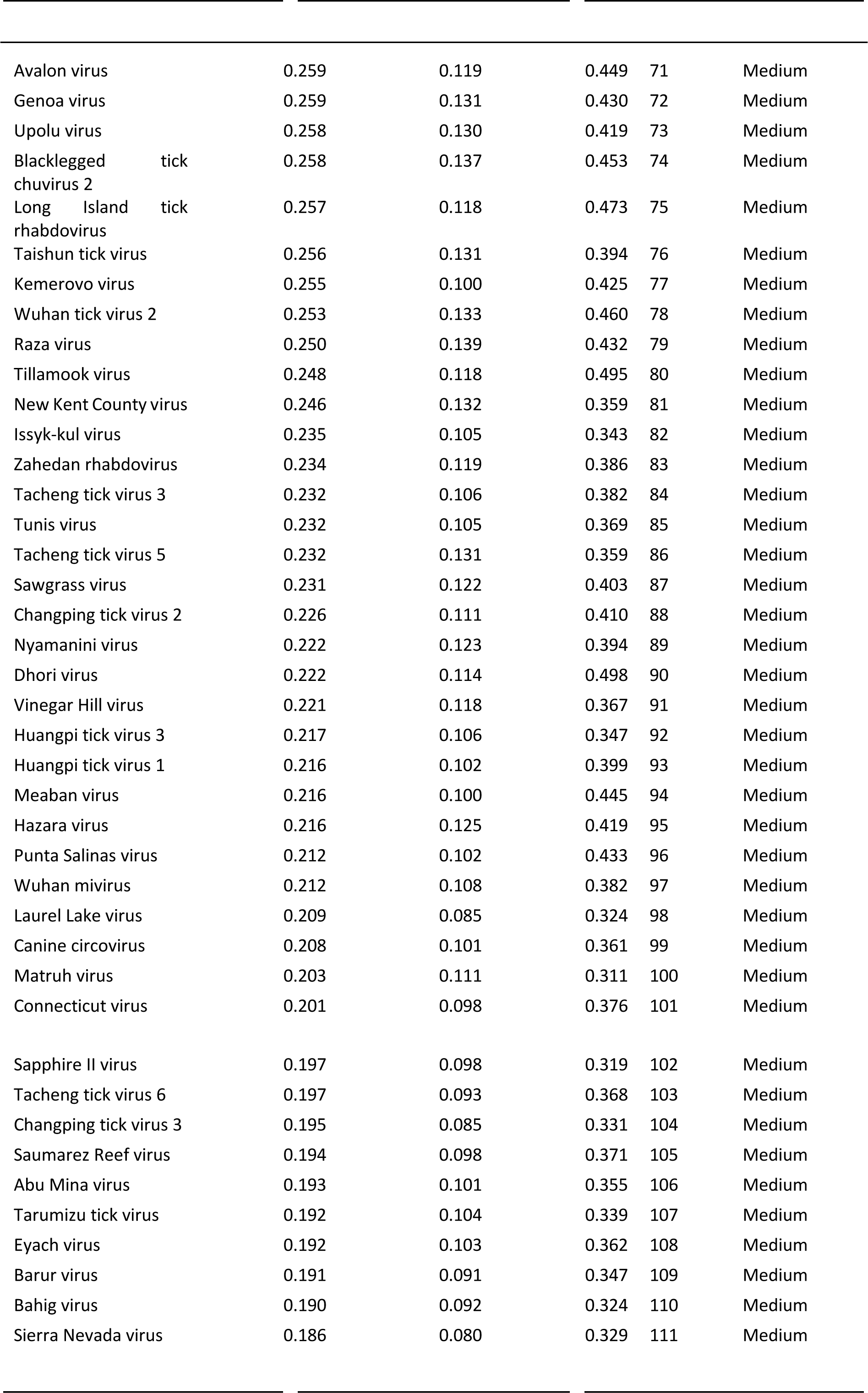
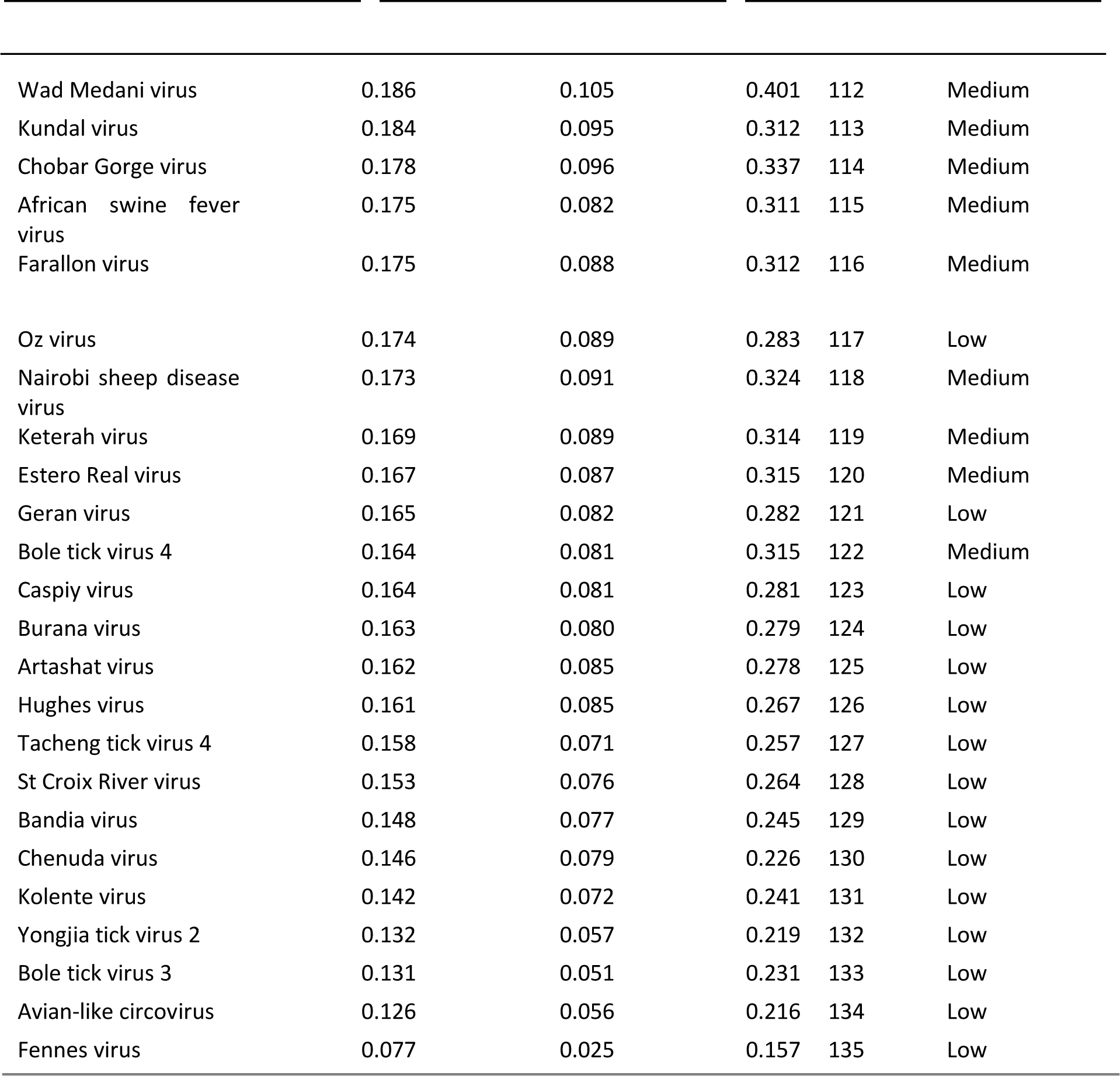
Zoonotic ranking of 136 known tick-borne viruses with complete genomes. Zoonotic risks of virus sequences (in GenBank format) were predicted using the GCB model.

