## Supplementary Table S1-6, Supplementary Figure S1-4 for "Characterisation and zoonotic risk of tick viruses in public datasets" (from another version of this preprint: version 1, 2022-12-05)

**SUPPLEMENTAL MATERIAL**

**Table S1.** Putative novel RNV viral contigs identified in this study.

**Table S2.** Putative novel RNA viral contigs that represent defective viruses.

**Table S3.** DNA viral contigs found after bioinformatic checking.

**Table S4.** GenBank ID of viruses phylogenetically related to the putative novel viral contigs. Zoonotic risks of virus sequences (in GenBank format) were predicted using the GCB model.

**Table S5.** Zoonotic ranking of 136 known tick-borne viruses with complete genomes. Zoonotic risks of virus sequences (in GenBank format) were predicted using the GCB model.

**Table S6.** Summary of currently known tick-borne viruses by 2021. Information from a wide variety of sources, references in the table.

**Figure S1.** Workflow of detecting viral RNA-dependent RNA polymerase sequences.

**Figure S2.** Phylogenetic relationship of the novel Orthomyxo-like viral contigs.

**Figure S3.** Phylogenetic relationship of the novel Alphatetra-like viral contigs.

**Figure S4.** Phylogenetic relationship of the novel Chu-like viral contigs.

**Table S1.** Putative novel RNV viral contigs identified in this study.

| **Putative novel viral contig** | **Genbank ID** | **Tick species** | **Provisional family** | **Genome structure** | **Sequence length (nt)** | **Closest BLASTn match (e-value 0.005)** | **Sequence identity (%)** | **Notes** | **Reference** |
| --- | --- | --- | --- | --- | --- | --- | --- | --- | --- |
| Alphatetra-like tick virus 1 | GIXL01013119.1 | *Ixodes ricinus* | Alphatetraviridae | ssRNA(+) | 5496 | Bulatov virus (MT025173.1) | 78 | One putative ORF; predicted protein contains homologs of RdRp 2 (PF00978) motif, nucleoside triphosphate hydrolase (SSF52540), viral (Superfamily 1) RNA helicase (PF01443) and viral methyltransferase 3 (PF01660) motifs. | Trentalman *et al*. 2020, 10.1038/s41598-020-76268-y |
| Alphatetra-like tick virus 2 | GBXQ01012426.1 | *Ixodes persulcatus* | Alphatetraviridae | ssRNA(+) | 1056 | Vovk virus (MN830235.1) | 81 | One putative ORF; predicted protein contains homologs of RdRp 2 (PF00978) motif. | Zhang 2014, Unpublished |
| Alphatetra-like tick virus 3 | GIXP02005464.1 | *Ornithodoros moubata* | Alphatetraviridae | ssRNA(+) | 5452 | Bulatov virus (MT025173.1) | 86 | One putative ORF; predicted protein contains homologs of RdRp 2 (PF00978) motif, nucleoside triphosphate hydrolase (SSF52540), viral (Superfamily 1) RNA helicase (PF01443) and viral methyltransferase 3 (PF01660) motifs. | Oleaga *et al*. 2021, 10.1371/journal.pntd.0009105 |
| Alphatetra-like tick virus 4 | GFJQ02007921.1 | *Ornithodoros moubata* | Alphatetraviridae | ssRNA(+) | 3873 | Bulatov virus (MT025173.1) | 67 | One putative ORF; predicted protein contains homologs of RdRp 2 (PF00978) motif, nucleoside triphosphate hydrolase (SSF52540) and viral (Superfamily 1) RNA helicase (PF01443) motifs. | Oleaga *et al*. 2017, 10.1016/j.ttbdis.2017.05.002 |
| Alphatetra-like tick virus 5 | GEMR01004364.1 | *Rhipicephalus microplus* | Alphatetraviridae | ssRNA(+) | 1488 | Hepelivirales sp. (MW722078.1) | 73 | One putative ORF; predicted protein contains homologs of RdRp 2 (PF00978) motif. | Munoz *et al*. 2017, 10.1371/journal.pone.0172326 |
| Orthomyxo-like tick virus 1 | HACP01027211.1 (HACW01024387.1) | *Rhipicephalus sanguineus* | Orthomyxoviridae | ssRNA(-) | 2136 | Zambezi tick virus 1 (MH267793.1) | 77 | One putative ORF; predicted protein contains homologs of Influenza RdRp subunit PB1 (PF00602) motif. | De Marco *et al*. 2015, Unpublished |
| Orthomyxo-like tick virus 2 | HACP01022575.1 | *Rhipicephalus sanguineus* | Orthomyxoviridae | ssRNA(-) | 1290 | Zambezi tick virus 1 (MH267793.1) | 77 | One putative ORF; predicted protein contains homologs of Influenza RdRp subunit PB1 (PF00602) motif. | De Marco *et al*. 2015, Unpublished |
| Orthomyxo-like tick virus 3 | HACW01018819.1 | *Rhipicephalus sanguineus* | Orthomyxoviridae | ssRNA(-) | 1290 | Zambezi tick virus 1 (MH267793.1) | 77 | One putative ORF; predicted protein contains homologs of Influenza RdRp subunit PB1 (PF00602) motif. | De Marco *et al*. 2015, Unpublished |
| Orthomyxo-like tick virus 4 | GIJA01018702.1 | *Rhipicephalus haemaphysaloides* | Orthomyxoviridae | ssRNA(-) | 2357 | Zambezi tick virus 1 (MH267793.1) | 77 | One putative ORF; predicted protein contains homologs of Influenza RdRp subunit PB1 (PF00602) motif. | Wang *et al*. 2020, 10.3389/fcimb.2020.00093 |
| Chu-like tick virus 1 | GGIX01201876.1 | *Ixodes scapularis* | Chuviridae | ssRNA(-) | 728 | Bole tick virus 3 (MH688550.1) | 87 | One putative ORF; predicted protein contains homologs of Mononegavirales RdRp (PF00946) motif. | Robertson 2016, Unpublished |

### **Table S2.** Putative novel RNA viral contigs that represent defective viruses.

| **SRA project ID** | **Tick species** | **Provisional family** | **Genome structure** | **Sequence length (nt)** | **Closest BLASTn match (e-value 0.005)** | **Sequence identity (%)** | **Reference** |
| --- | --- | --- | --- | --- | --- | --- | --- |
| SRR10549473.54432.1 | *Hy. asiaticum* | *Chuviridae* | ssRNA(-) | 4383 | Bole tick virus 3 (MH688550.1) | 84 | Yuan *et al*. 2020, 10.1016/j.jip.2020.107481 |
| SRR10549473.681872.1 | *Hy. asiaticum* | *Chuviridae* | ssRNA(-) | 4528 | Bole tick virus 3 (MH688550.1) | 82 | Yuan *et al*. 2020, 10.1016/j.jip.2020.107481 |
| SRR10549473.681877.1 | *Hy. asiaticum* | *Chuviridae* | ssRNA(-) | 4420 | Bole tick virus 3 (MH688550.1) | 84 | Yuan *et al*. 2020, 10.1016/j.jip.2020.107481 |
| SRR10549473.681878.1 | *Hy. asiaticum* | *Chuviridae* | ssRNA(-) | 4440 | Bole tick virus 3 (MH688550.1) | 86 | Yuan *et al*. 2020, 10.1016/j.jip.2020.107481 |
| SRR10549473.779074.1 | *Hy. asiaticum* | *Chuviridae* | ssRNA(-) | 5286 | Bole tick virus 3 (MH688550.1) | 84 | Yuan *et al*. 2020, 10.1016/j.jip.2020.107481 |
| SRR10549473.779079.1 | *Hy. asiaticum* | *Chuviridae* | ssRNA(-) | 5379 | Bole tick virus 3 (MH688550.1) | 85 | Yuan *et al*. 2020, 10.1016/j.jip.2020.107481 |
| SRR10549473.779081.1 | *Hy. asiaticum* | *Chuviridae* | ssRNA(-) | 5334 | Bole tick virus 3 (MH688550.1) | 83 | Yuan *et al*. 2020, 10.1016/j.jip.2020.107481 |
| SRR10549473.1447289.1 | *Hy. asiaticum* | *Chuviridae* | ssRNA(-) | 5272 | Bole tick virus 3 (MH688550.1) | 85 | Yuan *et al*. 2020, 10.1016/j.jip.2020.107481 |
| SRR10549473.1447293.1 | *Hy. asiaticum* | *Chuviridae* | ssRNA(-) | 5001 | Bole tick virus 3 (MH688550.1) | 87 | Yuan *et al*. 2020, 10.1016/j.jip.2020.107481 |
| SRR10549473.2865317.1 | *Hy. asiaticum* | *Chuviridae* | ssRNA(-) | 5556 | Bole tick virus 3 (MH688550.1) | 83 | Yuan *et al*. 2020, 10.1016/j.jip.2020.107481 |
| SRR10549473.2865320.1 | *Hy. asiaticum* | *Chuviridae* | ssRNA(-) | 5643 | Bole tick virus 3 (MH688550.1) | 87 | Yuan *et al*. 2020, 10.1016/j.jip.2020.107481 |
| SRR10549473.4598504.1 | *Hy. asiaticum* | *Chuviridae* | ssRNA(-) | 8170 | Bole tick virus 3 (MH688550.1) | 88 | Yuan *et al*. 2020, 10.1016/j.jip.2020.107481 |
| SRR10549473.4598510.1 | *Hy. asiaticum* | *Chuviridae* | ssRNA(-) | 8206 | Bole tick virus 3 (MH688550.1) | 87 | Yuan *et al*. 2020, 10.1016/j.jip.2020.107481 |
| SRR10549473.4598511.1 | *Hy. asiaticum* | *Chuviridae* | ssRNA(-) | 8182 | Bole tick virus 3 (MH688550.1) | 85 | Yuan *et al*. 2020, 10.1016/j.jip.2020.107481 |
| SRR10549473.7000060.1 | *Hy. asiaticum* | *Chuviridae* | ssRNA(-) | 4986 | Bole tick virus 3 (MH688550.1) | 84 | Yuan *et al*. 2020, 10.1016/j.jip.2020.107481 |
| SRR10549473.10341027.1 | *Hy. asiaticum* | *Chuviridae* | ssRNA(-) | 2448 | Bole tick virus 3 (MH688550.1) | 86 | Yuan *et al*. 2020, 10.1016/j.jip.2020.107481 |
| SRR10549473.10526706.1 | *Hy. asiaticum* | *Chuviridae* | ssRNA(-) | 6273 | Bole tick virus 3 (MH688550.1) | 87 | Yuan *et al*. 2020, 10.1016/j.jip.2020.107481 |
| SRR10549473.10526708.1 | *Hy. asiaticum* | *Chuviridae* | ssRNA(-) | 6276 | Nuomin virus (MW029964.1) | 78 | Yuan *et al*. 2020, 10.1016/j.jip.2020.107481 |
| SRR17281102.635386.1 | *I. scapualris* | *Chuviridae* | ssRNA(-) | 1390 | Suffolk virus (NC_028243.1) | 78 | University of Minnesota 2021, Unpublished |

### **Table S3.** DNA viral contigs found after bioinformatic checking.

| **WGS project ID** | **Tick species** | **Genome structure** | **Sequence length (bp)** | **Predicted protein** | **Reference** |
| --- | --- | --- | --- | --- | --- |
| GIXP02029293.1 | *Ornithodoros moubata* | dsDNA | 2232 | DdRp subunit 3 (encoded by *H359L*); ATP-dependent RNA helicase (encoded by *Q706L*); ATP-dependent RNA helicase (encoded by *QP509L*) | Oleaga *et al*. 2021, 10.1371/journal.pntd.0009105 |
| GIXP02006837.1 | *Ornithodoros moubata* | dsDNA | 1450 | DdRp subunit 3 (encoded by *H359L*); Uncharacterized protein (encoded by *H171R)* | Oleaga *et al*. 2021, 10.1371/journal.pntd.0009105 |
| CAJHNL010000002.1 | *Ornithodoros porcinus* | dsDNA | 7421 | DdRp subunit 2 (encoded by *EP1242L*); Helicase (encoded by *F1055L*) | Forth *et al*. 2020, 10.1186/s12915-020-00865-6 |
| CAJHNL010000003.1 | *Ornithodoros porcinus* | dsDNA | 6757 | Transcription factor SII homolog (encoded by *I243L*); pI226R; RPB7 homolog; Uncharacterized protein (encoded by *D339L)*; Uncharacterized protein (encoded by *D129L*) | Forth *et al*. 2020, 10.1186/s12915-020-00865-6 |
| CAJHNL010000004.1 | *Ornithodoros porcinus* | dsDNA | 6577 | Primase (encoded by *C962R*) | Forth *et al*. 2020, 10.1186/s12915-020-00865-6 |
| CAJHNL010000005.1 | *Ornithodoros porcinus* | dsDNA | 3455 | TATA-binding protein (encoded by *B263R*) | Forth *et al*. 2020, 10.1186/s12915-020-00865-6 |
| CAJHNL010000006.1 | *Ornithodoros porcinus* | dsDNA | 3089 | DdRp subunit 2 (encoded by *EP1242L*) | Forth *et al*. 2020, 10.1186/s12915-020-00865-6 |
| CAJHNL010000007.1 | *Ornithodoros porcinus* | dsDNA | 1557 | DdDp (encoded by *G1211R*) | Forth *et al*. 2020, 10.1186/s12915-020-00865-6 |
| CAJHNL010000008.1 | *Ornithodoros porcinus* | dsDNA | 629 | Ribonucleotide reductase large subunit (encoded by *F778R*); Uncharacterized protein p360 15R | Forth *et al*. 2020, 10.1186/s12915-020-00865-6 |
| CAJHNP010000002.1 | *Ornithodoros moubata* | dsDNA | 7383 | DdRp subunit 2 (encoded by *EP1242L*); Ribonucleotide reductase large subunit (encoded by *F778R*); Helicase (encoded by *F1055L*) | Forth et al. 2020, 10.1186/s12915-020-00865-6 |
| CAJHNP010000004.1 | *Ornithodoros moubata* | dsDNA | 670 | DdRp subunit 3 (encoded by *H359L*) | Forth et al. 2020, 10.1186/s12915-020-00865-6 |
| CAJHNP010000005.1 | *Ornithodoros moubata* | dsDNA | 5156 | Viral histone-like protein (encoded by *A104R*); Inhibitor of apoptosis protein (encoded by *A224L*); Thymidylate kinase (encoded by *A240L*); Uncharacterized protein (encoded by *A151R*) | Forth et al. 2020, 10.1186/s12915-020-00865-6 |
| CAJHNP010000006.1 | *Ornithodoros moubata* | dsDNA | 3634 | DdRp subunit 3 (encoded by *H359L*); Uncharacterized protein *CP123L*; ATP-dependent RNA helicase (encoded by *QP509L*); DNA topoisomerase 2 (encoded by *P1192R*) | Forth et al. 2020, 10.1186/s12915-020-00865-6 |
| CAJHNP010000007.1 | *Ornithodoros moubata* | dsDNA | 2713 | Early transcription factor large subunit (encoded by *D1133L*); Uncharacterized protein (encoded by *H124R*); Uncharacterized protein (encoded by *H171R*) | Forth et al. 2020, 10.1186/s12915-020-00865-6 |
| CAJHNP010000017.1 | *Ornithodoros moubata* | dsDNA | 1007 | Primase (encoded by *C962R*) | Forth et al. 2020, 10.1186/s12915-020-00865-6 |
| CAJHNQ010000010.1 | *Ornithodoros moubata* | dsDNA | 636 | DdRp subunit 2 (encoded by *EP1242L*) | Forth et al. 2020, 10.1186/s12915-020-00865-6 |
| CAJHNQ010000014.1 | *Ornithodoros moubata* | dsDNA | 679 | DdRp subunit 2 (encoded by *EP1242L*); Ribonucleotide reductase large subunit (encoded by *F778R*); Helicase (encoded by *F1055L*) | Forth et al. 2020, 10.1186/s12915-020-00865-6 |
| CAJHNQ010000015.1 | *Ornithodoros moubata* | dsDNA | 1193 | Helicase (encoded by *F1055L*) | Forth et al. 2020, 10.1186/s12915-020-00865-6 |
| CAJHNQ010000016.1 | *Ornithodoros moubata* | dsDNA | 1697 | DdRp subunit 2 (encoded by *EP1242L*); Helicase (encoded by *F1055L*) | Forth et al. 2020, 10.1186/s12915-020-00865-6 |
| CAJHNR010000004.1 | *Ornithodoros moubata* | dsDNA | 16841 | DdRp subunit 2 (encoded by *EP1242L*); Ribonucleotide reductase large subunit (encoded by *F778R*); Helicase (encoded by *F1055L*) | Forth et al. 2020, 10.1186/s12915-020-00865-6 |
| CAJHNR010000005.1 | *Ornithodoros moubata* | dsDNA | 15286 | TATA-binding protein (encoded by *B263R*) | Forth et al. 2020, 10.1186/s12915-020-00865-6 |
| CAJHNR010000006.1 | *Ornithodoros moubata* | dsDNA | 13138 | TATA-binding protein (encoded by *B263R*) | Forth et al. 2020, 10.1186/s12915-020-00865-6 |
| CAJHNR010000011.1 | *Ornithodoros moubata* | dsDNA | 8352 | Helicase (encoded by *A859L*); Uncharacterized protein p360 15R; Uncharacterized protein 5ELp28; IkB-like protein (encoded by *A238L*) | Forth et al. 2020, 10.1186/s12915-020-00865-6 |
| CAJHNR010000013.1 | *Ornithodoros moubata* | dsDNA | 7718 | Early transcription factor large subunit (encoded by *D1133L*); DdRp subunit 3 (encoded by *H359L*); Uncharacterized protein H171R | Forth et al. 2020, 10.1186/s12915-020-00865-6 |
| CAJHNR010000014.1 | *Ornithodoros moubata* | dsDNA | 7045 | TATA-binding protein (encoded by *B263R*) | Forth et al. 2020, 10.1186/s12915-020-00865-6 |
| CAJHNR010000015.1 | *Ornithodoros moubata* | dsDNA | 6370 | TATA-binding protein (encoded by *B263R*); Early transcription factor large subunit (encoded by *G1340L*); Primase (encoded by *C962R*) | Forth et al. 2020, 10.1186/s12915-020-00865-6 |
| CAJHNR010000016.1 | *Ornithodoros moubata* | dsDNA | 5808 | TATA-binding protein (encoded by *B263R*) | Forth et al. 2020, 10.1186/s12915-020-00865-6 |
| CAJHNR010000017.1 | *Ornithodoros moubata* | dsDNA | 5688 | TATA-binding protein (encoded by *B263R*) | Forth et al. 2020, 10.1186/s12915-020-00865-6 |
| CAJHNR010000018.1 | *Ornithodoros moubata* | dsDNA | 5489 | DdRp subunit 3 (encoded by *H359L*); Uncharacterized protein (encoded by *CP123L*); DNA topoisomerase II (encoded by *P1192R*) | Forth et al. 2020, 10.1186/s12915-020-00865-6 |
| CAJHNR010000019.1 | *Ornithodoros moubata* | dsDNA | 5396 | Uncharacterized protein p360 15R | Forth et al. 2020, 10.1186/s12915-020-00865-6 |
| CAJHNR010000020.1 | *Ornithodoros moubata* | dsDNA | 5152 | Histone-like protein (encoded by *A104R*); Inhibitor of apoptosis protein (encoded by *A224L*); Thymidylate kinase (encoded by *A240L*); Uncharacterized protein (encoded by *A151R*) | Forth et al. 2020, 10.1186/s12915-020-00865-6 |
| CAJHNR010000021.1 | *Ornithodoros moubata* | dsDNA | 4920 | DdRp subunit 10 (encoded by *CP80R*); Uncharacterized protein (encoded by *CP312R*) | Forth et al. 2020, 10.1186/s12915-020-00865-6 |
| CAJHNR010000023.1 | *Ornithodoros moubata* | dsDNA | 4152 | Primase (encoded by *C962R*) | Forth et al. 2020, 10.1186/s12915-020-00865-6 |
| CAJHNR010000024.1 | *Ornithodoros moubata* | dsDNA | 3824 | DdRp subunit 2 (encoded by *EP1242L*); Transmembrane protein (encoded by *EP84R*) | Forth et al. 2020, 10.1186/s12915-020-00865-6 |
| CAJHNR010000025.1 | *Ornithodoros moubata* | dsDNA | 3339 | DdRp subunit 2 (encoded by *EP1242L*) | Forth et al. 2020, 10.1186/s12915-020-00865-6 |
| CAJHNR010000026.1 | *Ornithodoros moubata* | dsDNA | 2712 | DdRp subunit 2 (encoded by *EP1242L*) | Forth et al. 2020, 10.1186/s12915-020-00865-6 |
| CAJHNR010000027.1 | *Ornithodoros moubata* | dsDNA | 2653 | Putative poly(A) polymerase catalytic subunit (encoded by *C475L*); Primase (encoded by *C962R*) | Forth et al. 2020, 10.1186/s12915-020-00865-6 |
| CAJHNR010000028.1 | *Ornithodoros moubata* | dsDNA | 2475 | DdRp subunit 3 (encoded by *H359L*) | Forth et al. 2020, 10.1186/s12915-020-00865-6 |
| CAJHNR010000029.1 | *Ornithodoros moubata* | dsDNA | 2196 | Ribonucleotide reductase large subunit (encoded by *F778R*) | Forth et al. 2020, 10.1186/s12915-020-00865-6 |
| CAJHNR010000030.1 | *Ornithodoros moubata* | dsDNA | 2179 | Uncharacterized protein (encoded by CP123L); mRNA-capping enzyme (encoded by *NP868R*) | Forth et al. 2020, 10.1186/s12915-020-00865-6 |
| CAJHNR010000032.1 | *Ornithodoros moubata* | dsDNA | 1763 | Ribonucleotide reductase large subunit (encoded by *F778R*) | Forth et al. 2020, 10.1186/s12915-020-00865-6 |
| CAJHNS010000026.1 | *Ornithodoros moubata* | dsDNA | 1078 | Helicase (encoded by *F1055L*) | Forth et al. 2020, 10.1186/s12915-020-00865-6 |
| CAJHNU010000002.1 | *Ornithodoros porcinus* | dsDNA | 3380 | DdRp subunit 2 (encoded by *EP1242L*); Helicase (encoded by *F1055L*) | Forth et al. 2020, 10.1186/s12915-020-00865-6 |
| CAJHNU010000003.1 | *Ornithodoros porcinus* | dsDNA | 641 | Ribonucleotide reductase large subunit (encoded by *F778R*) | Forth et al. 2020, 10.1186/s12915-020-00865-6 |
| CAJHNU010000004.1 | *Ornithodoros porcinus* | dsDNA | 1803 | TATA-binding protein (encoded by *B263R*) | Forth et al. 2020, 10.1186/s12915-020-00865-6 |
| CAJHNU010000009.1 | *Ornithodoros porcinus* | dsDNA | 776 | Transcription factor S-II-related protein (encoded by *I243L*); Late protein (encoded by *I226R*) | Forth et al. 2020, 10.1186/s12915-020-00865-6 |
| CAJHNU010000011.1 | *Ornithodoros porcinus* | dsDNA | 2551 | Primase (encoded by *C962R*) | Forth et al. 2020, 10.1186/s12915-020-00865-6 |
| CAJHNU010000014.1 | *Ornithodoros porcinus* | dsDNA | 3100 | DdRp subunit 2 (encoded by *EP1242L*) | Forth et al. 2020, 10.1186/s12915-020-00865-6 |
| CAJHNV010000002.1 | *Ornithodoros porcinus* | dsDNA | 4758 | DdRp subunit 2 (encoded by *EP1242L*) | Forth et al. 2020, 10.1186/s12915-020-00865-6 |
| CAJHNV010000004.1 | *Ornithodoros porcinus* | dsDNA | 3378 | Transcription factor SII homolog (encoded by *I243L*); Late protein (encoded by *I226R*); Uncharacterized protein (encoded by *D79L*) | Forth et al. 2020, 10.1186/s12915-020-00865-6 |
| CAJHNV010000005.1 | *Ornithodoros porcinus* | dsDNA | 2605 | DdRp subunit 2 (encoded by *EP1242L*) | Forth et al. 2020, 10.1186/s12915-020-00865-6 |
| CAJHNV010000007.1 | *Ornithodoros porcinus* | dsDNA | 777 | Transcription factor SII homolog (encoded by *I243L*) | Forth et al. 2020, 10.1186/s12915-020-00865-6 |
| CAJHNW010000016.1 | *Ornithodoros moubata* | dsDNA | 1078 | Helicase (encoded by *F1055L*) | Forth et al. 2020, 10.1186/s12915-020-00865-6 |

**Table S4.** GenBank ID of viruses phylogenetically related to the putative novel viral contigs. Zoonotic risks of virus sequences (in GenBank format) were predicted using the GCB model.

| **Putative novel viral contigs** | **Relative viral species** | **GenBank ID** |
| --- | --- | --- |
| Alphatetra-like viral contigs | *Nudaurelia capensis beta virus* | NC_001990.1 |
|  |  | NC_005898.1; NC_005899.1 |
|  | *Helicoverpa armigera stunt virus* | NC_001981.1; NC_001982.1 |
| Orthomyxo-like viral contigs | *Wellfleet Bay virus* | NC_025798.1; NC_025797.1; NC_025796.1; NC_025795.1; NC_025794.1; NC_025793.1; NC_025799.1 |
|  | *Araguari virus* | KX670389.1; KX670390.1; KX670391.1; KX670392.1; KX670393.1; KX670394.1; KX670395.1 |
|  | *Johnston Atoll quaranjavirus* | NC_052931.1; NC_052930.1; NC_052929.1; NC_052928.1; NC_052927.1; NC_052926.1; NC_052925.1 |
|  | *Lake Chad virus* | NC_052682.1; NC_052686.1; NC_052685.1; NC_052684.1; NC_052683.1; NC_052681.1; NC_052680.1 |
|  | *Tjuloc virus* | JQ928941.1; JQ928942.1; JQ928943.1; JQ928944.1; JQ928945.1; JQ928946.1; MT774252.1 |
|  | *Quaranfil quaranjavirus* | NC_038821.1; NC_038822.1; NC_038820.1; NC_038819.1; NC_038818.1; NC_038817.1 |
| Chu-like viral contigs | *Mivirus suffolkense* | NC_028243 |

**Table S5.** Zoonotic ranking of 136 known tick-borne viruses with complete genomes. Zoonotic risks of virus sequences (in GenBank format) were predicted using the GCB model.

| **Virus name** | **Predicted probability mean** | **Predicted probability lower** | **Predicted probability upper** | **Rank** | **Priority** |
| --- | --- | --- | --- | --- | --- |
| Tick-borne encephalitis virus | 0.777 | 0.446 | 0.946 | 1 | Very high |
| Omsk hemorrhagic fever virus | 0.739 | 0.443 | 0.951 | 2 | Very high |
| Powassan virus | 0.695 | 0.288 | 0.920 | 3 | High |
| Isfahan virus | 0.677 | 0.233 | 0.917 | 4 | High |
| Isfahan virus | 0.677 | 0.233 | 0.917 | 4 | High |
| Uukuniemi virus | 0.676 | 0.262 | 0.944 | 5 | High |
| Louping ill virus | 0.668 | 0.288 | 0.890 | 6 | High |
| Langat virus | 0.667 | 0.339 | 0.917 | 7 | Very high |
| Thogoto virus | 0.642 | 0.266 | 0.899 | 8 | High |
| Lonestar tick chuvirus 1 | 0.619 | 0.379 | 0.857 | 9 | Very high |
| Crimean-Congo hemorrhagic fever virus | 0.609 | 0.249 | 0.875 | 10 | High |
| Grotenhout virus | 0.601 | 0.389 | 0.808 | 11 | Very high |
| Severe fever with thrombocytopenia syndrome virus | 0.596 | 0.273 | 0.849 | 12 | High |
| Kasokero virus | 0.580 | 0.180 | 0.837 | 13 | High |
| Colorado tick fever virus | 0.562 | 0.121 | 0.854 | 14 | High |
| Quaranfil virus | 0.561 | 0.128 | 0.863 | 15 | High |
| Great Island virus | 0.550 | 0.115 | 0.832 | 16 | High |
| Dugbe virus | 0.540 | 0.157 | 0.825 | 17 | High |
| Johnston Atoll virus | 0.513 | 0.311 | 0.802 | 18 | Very high |
| Dabieshan tick virus | 0.505 | 0.289 | 0.747 | 19 | High |
| Taggert virus | 0.503 | 0.318 | 0.802 | 20 | Very high |
| Blacklegged tick phlebovirus 1 | 0.490 | 0.271 | 0.711 | 21 | High |
| Deer tick virus | 0.486 | 0.275 | 0.703 | 22 | High |
| Tacheng tick virus 2 | 0.482 | 0.241 | 0.774 | 23 | High |
| Lihan tick virus | 0.481 | 0.264 | 0.730 | 24 | High |
| Karshi virus | 0.481 | 0.266 | 0.954 | 25 | High |
| Alkhumra hemorrhagic fever virus | 0.469 | 0.234 | 0.736 | 26 | High |
| Yongjia tick virus 1 | 0.441 | 0.192 | 0.680 | 27 | High |
| American dog tick phlebovirus | 0.438 | 0.218 | 0.659 | 28 | High |
| Wenzhou tick virus | 0.425 | 0.234 | 0.710 | 29 | High |
| Tamdy virus | 0.413 | 0.227 | 0.642 | 30 | High |
| Blacklegged tick rhabdovirus 1 | 0.411 | 0.223 | 0.638 | 31 | High |
| Tacheng tick virus 8 | 0.404 | 0.210 | 0.692 | 32 | High |
| Jingmen tick virus | 0.401 | 0.219 | 0.663 | 33 | High |
| South Bay virus | 0.395 | 0.210 | 0.630 | 34 | High |
| Ngari virus | 0.393 | 0.214 | 0.624 | 35 | High |
| Kaisodi virus | 0.392 | 0.208 | 0.590 | 36 | High |
| Kupe virus | 0.391 | 0.160 | 0.642 | 37 | High |
| Lone star tick densovirus 1 | 0.391 | 0.221 | 0.654 | 38 | High |
| Blacklegged tick phlebovirus 3 | 0.371 | 0.195 | 0.640 | 39 | High |
| Pacific coast tick nairovirus | 0.370 | 0.197 | 0.656 | 40 | High |
| Clo Mor virus | 0.364 | 0.149 | 0.600 | 41 | High |
| Sakhalin virus | 0.345 | 0.177 | 0.657 | 42 | High |
| Tyuleniy virus | 0.340 | 0.160 | 0.739 | 43 | High |
| Meram virus | 0.335 | 0.196 | 0.574 | 44 | High |
| Jos virus | 0.333 | 0.159 | 0.579 | 45 | High |
| Midway virus | 0.327 | 0.163 | 0.609 | 46 | High |
| lone star virus | 0.326 | 0.166 | 0.567 | 47 | High |
| Norway phlebovirus 1 | 0.323 | 0.138 | 0.559 | 48 | High |
| Tofla virus | 0.323 | 0.198 | 0.601 | 49 | High |
| Tacheng tick virus 7 | 0.318 | 0.175 | 0.511 | 50 | High |
| New Minto virus | 0.318 | 0.168 | 0.526 | 51 | High |
| Bourbon virus | 0.311 | 0.164 | 0.497 | 52 | High |
| Suffolk virus | 0.309 | 0.151 | 0.562 | 53 | High |
| Silverwater virus | 0.307 | 0.156 | 0.537 | 54 | High |
| Heartland virus | 0.305 | 0.135 | 0.520 | 55 | High |
| Dera Ghazi Khan virus | 0.299 | 0.150 | 0.493 | 56 | High |
| Hunter Island virus | 0.299 | 0.153 | 0.528 | 57 | High |
| Great Saltee virus | 0.299 | 0.168 | 0.543 | 58 | High |
| Bhanja virus | 0.298 | 0.161 | 0.446 | 59 | High |
| Blanchseco virus | 0.295 | 0.153 | 0.500 | 60 | High |
| Thailand tick thogotovirus | 0.286 | 0.135 | 0.436 | 61 | Medium |
| Wuhan tick virus 1 | 0.280 | 0.145 | 0.461 | 62 | Medium |
| Gadgets Gully virus | 0.277 | 0.152 | 0.583 | 63 | Medium |
| Soldado virus | 0.277 | 0.164 | 0.450 | 64 | Medium |
| Abu Hammad virus | 0.275 | 0.138 | 0.471 | 65 | Medium |
| Kadam virus | 0.275 | 0.134 | 0.602 | 66 | Medium |
| Tacheng tick virus 1 | 0.272 | 0.134 | 0.514 | 67 | Medium |
| Zirqa virus | 0.269 | 0.146 | 0.439 | 68 | Medium |
| Bole tick virus 2 | 0.267 | 0.141 | 0.466 | 69 | Medium |
| Qalyub virus | 0.264 | 0.112 | 0.444 | 70 | Medium |
| Avalon virus | 0.259 | 0.119 | 0.449 | 71 | Medium |
| Genoa virus | 0.259 | 0.131 | 0.430 | 72 | Medium |
| Upolu virus | 0.258 | 0.130 | 0.419 | 73 | Medium |
| Blacklegged tick chuvirus 2 | 0.258 | 0.137 | 0.453 | 74 | Medium |
| Long Island tick rhabdovirus | 0.257 | 0.118 | 0.473 | 75 | Medium |
| Taishun tick virus | 0.256 | 0.131 | 0.394 | 76 | Medium |
| Kemerovo virus | 0.255 | 0.100 | 0.425 | 77 | Medium |
| Wuhan tick virus 2 | 0.253 | 0.133 | 0.460 | 78 | Medium |
| Raza virus | 0.250 | 0.139 | 0.432 | 79 | Medium |
| Tillamook virus | 0.248 | 0.118 | 0.495 | 80 | Medium |
| New Kent County virus | 0.246 | 0.132 | 0.359 | 81 | Medium |
| Issyk-kul virus | 0.235 | 0.105 | 0.343 | 82 | Medium |
| Zahedan rhabdovirus | 0.234 | 0.119 | 0.386 | 83 | Medium |
| Tacheng tick virus 3 | 0.232 | 0.106 | 0.382 | 84 | Medium |
| Tunis virus | 0.232 | 0.105 | 0.369 | 85 | Medium |
| Tacheng tick virus 5 | 0.232 | 0.131 | 0.359 | 86 | Medium |
| Sawgrass virus | 0.231 | 0.122 | 0.403 | 87 | Medium |
| Changping tick virus 2 | 0.226 | 0.111 | 0.410 | 88 | Medium |
| Nyamanini virus | 0.222 | 0.123 | 0.394 | 89 | Medium |
| Dhori virus | 0.222 | 0.114 | 0.498 | 90 | Medium |
| Vinegar Hill virus | 0.221 | 0.118 | 0.367 | 91 | Medium |
| Huangpi tick virus 3 | 0.217 | 0.106 | 0.347 | 92 | Medium |
| Huangpi tick virus 1 | 0.216 | 0.102 | 0.399 | 93 | Medium |
| Meaban virus | 0.216 | 0.100 | 0.445 | 94 | Medium |
| Hazara virus | 0.216 | 0.125 | 0.419 | 95 | Medium |
| Punta Salinas virus | 0.212 | 0.102 | 0.433 | 96 | Medium |
| Wuhan mivirus | 0.212 | 0.108 | 0.382 | 97 | Medium |
| Laurel Lake virus | 0.209 | 0.085 | 0.324 | 98 | Medium |
| Canine circovirus | 0.208 | 0.101 | 0.361 | 99 | Medium |
| Matruh virus | 0.203 | 0.111 | 0.311 | 100 | Medium |
| Connecticut virus | 0.201 | 0.098 | 0.376 | 101 | Medium |
| Sapphire II virus | 0.197 | 0.098 | 0.319 | 102 | Medium |
| Tacheng tick virus 6 | 0.197 | 0.093 | 0.368 | 103 | Medium |
| Changping tick virus 3 | 0.195 | 0.085 | 0.331 | 104 | Medium |
| Saumarez Reef virus | 0.194 | 0.098 | 0.371 | 105 | Medium |
| Abu Mina virus | 0.193 | 0.101 | 0.355 | 106 | Medium |
| Tarumizu tick virus | 0.192 | 0.104 | 0.339 | 107 | Medium |
| Eyach virus | 0.192 | 0.103 | 0.362 | 108 | Medium |
| Barur virus | 0.191 | 0.091 | 0.347 | 109 | Medium |
| Bahig virus | 0.190 | 0.092 | 0.324 | 110 | Medium |
| Sierra Nevada virus | 0.186 | 0.080 | 0.329 | 111 | Medium |
| Wad Medani virus | 0.186 | 0.105 | 0.401 | 112 | Medium |
| Kundal virus | 0.184 | 0.095 | 0.312 | 113 | Medium |
| Chobar Gorge virus | 0.178 | 0.096 | 0.337 | 114 | Medium |
| African swine fever virus | 0.175 | 0.082 | 0.311 | 115 | Medium |
| Farallon virus | 0.175 | 0.088 | 0.312 | 116 | Medium |
| Oz virus | 0.174 | 0.089 | 0.283 | 117 | Low |
| Nairobi sheep disease virus | 0.173 | 0.091 | 0.324 | 118 | Medium |
| Keterah virus | 0.169 | 0.089 | 0.314 | 119 | Medium |
| Estero Real virus | 0.167 | 0.087 | 0.315 | 120 | Medium |
| Geran virus | 0.165 | 0.082 | 0.282 | 121 | Low |
| Bole tick virus 4 | 0.164 | 0.081 | 0.315 | 122 | Medium |
| Caspiy virus | 0.164 | 0.081 | 0.281 | 123 | Low |
| Burana virus | 0.163 | 0.080 | 0.279 | 124 | Low |
| Artashat virus | 0.162 | 0.085 | 0.278 | 125 | Low |
| Hughes virus | 0.161 | 0.085 | 0.267 | 126 | Low |
| Tacheng tick virus 4 | 0.158 | 0.071 | 0.257 | 127 | Low |
| St Croix River virus | 0.153 | 0.076 | 0.264 | 128 | Low |
| Bandia virus | 0.148 | 0.077 | 0.245 | 129 | Low |
| Chenuda virus | 0.146 | 0.079 | 0.226 | 130 | Low |
| Kolente virus | 0.142 | 0.072 | 0.241 | 131 | Low |
| Yongjia tick virus 2 | 0.132 | 0.057 | 0.219 | 132 | Low |
| Bole tick virus 3 | 0.131 | 0.051 | 0.231 | 133 | Low |
| Avian-like circovirus | 0.126 | 0.056 | 0.216 | 134 | Low |
| Fennes virus | 0.077 | 0.025 | 0.157 | 135 | Low |

**Table S6.** Summary of currently known tick-borne viruses by 2021. Information from a wide variety of sources, references in the table.

| **Virus** | **Species** | **Genus** | **Family** | **Genome structure** | **Accession** | **Available sequence** | **Human infection** | **First description** | **Tick vectors** | **Distribution** |
| --- | --- | --- | --- | --- | --- | --- | --- | --- | --- | --- |
| African swine fever virus | *African swine fever virus* | *Asfivirus* | *Asfarviridae* | dsDNA | NC_044942 | Complete genome | ? | 1921 (Montgomery et al., 1921) | *O. erraticus, O. moubata* (Shi et al., 2018) | Sub-Saharan Africa, South America, Southern Europe (Shi et al., 2018) |
| Canine circovirus | *Canine circovirus* | *Circovirus* | *Circoviridae* | ssDNA (+/-) | NC_020904 | Complete genome | ? | 2012 (Kapoor et al., 2012) | *Haemaphysalis spp., Rhipicephalus spp.* (Kapoor et al., 2012) | Australia, Brazil, China, Columbia, Croatia, Germany, Iran, Italy, Norway, Thailand, Turkey, UK, USA, Viet Nam (Urbani et al., 2021) |
| Avian-like circovirus | *Tick associated circovirus 1* | *Circovirus* | *Circoviridae* | ssDNA (+/-) | NC_030199 | Complete genome | ? | 2018 (Tokarz et al., 2018) | *I. scapularis (Tokarz et al., 2018)* | USA (Tokarz et al., 2018) |
| Nayun tick torquevirus | Unclassified | *Thetatorquevirus* | *Anelloviridae* | ssDNA (+/-) | KP141758 | Partial genome | ? | 2015 (Xia et al., 2015) | *Rhipicephalus* spp. (Xia et al., 2015) | China (Xia et al., 2015) |
| Lone star tick densovirus 1 | Unclassified | Unclassified | *Parvoviridae* | ssDNA (+/-) | KX774632 | Complete genome | ? | 2018 (Tokarz et al., 2018) | *I. scapularis (Tokarz et al., 2018)* | USA (Tokarz et al., 2018) |
| Colorado tick fever virus | *Colorado tick fever coltivirus* | *Coltivirus* | *Reoviridae* | dsRNA | NC_004191, NC_004181, NC_004182, NC_004183, NC_004184, NC_004185, NC_004186, NC_004187, NC_004188, NC_004180, NC_004189, NC_004190 | Complete genome | Y (Shi et al., 2018) | 1946 (Florio et al., 1946) | *D. andersoni*, *D. albipictus*, *D. arumapertus*, D. occidentalis, *H. leporispalustris*, I. sculptus, I. spinipalpis, *Ot. lagophilus* (Shi et al., 2018) | USA (Shi et al., 2018) |
| Eyach virus | *Eyach coltivirus* | *Coltivirus* | *Reoviridae* | dsRNA | NC_003696, NC_003697, NC_003698, NC_003699, NC_003700, NC_003701, NC_003702, NC_003703, NC_003704, NC_003705, NC_003706, NC_003707 | Complete genome | ? | 1976 (Rehse-Kupper et al., 1976) | *I. ricinus*, *I. ventalloi* (Shi et al., 2018) | France, Germany (Shi et al., 2018) |
| Kundal virus | *Kundal coltivirus* | *Coltivirus* | *Reoviridae* | dsRNA | NC_055248, NC_055249, NC_055247, NC_055240, NC_055250, NC_055239, NC_055242, NC_055245, NC_055241, NC_055246, NC_055243, NC_055244 | Complete genome | ? | 2019 (Yadav et al., 2019) | *Hy. antolicum* (Yadav et al., 2019) | India (Yadav et al., 2019) |
| Tarumizu tick virus | *Tarumizu coltivirus* | *Coltivirus* | *Reoviridae* | dsRNA | NC_055255, NC_055254, NC_055261, NC_055259, NC_055262, NC_055251, NC_055260, NC_055258, NC_055252, NC_055257, NC_055253, NC_055256 | Complete genome | ? | 2017 (Fujita et al., 2017) | *H. flava* (Fujita et al., 2017) | Japan (Fujita et al., 2017) |
| Chenuda virus | *Chenuda virus* | *Orbivirus* | *Reoviridae* | dsRNA | NC_027534, NC_027535, NC_027536, NC_027537, NC_027538, NC_027548, NC_027549, NC_027550, NC_027551, NC_027552 | Complete genome | ? | 1978 (Karabatsos, 1978) | *A. hermanni* (Shi et al., 2018) | Egypt, Uzbekistan (Shi et al., 2018) |
| Chobar Gorge virus | *Chobar Gorge virus* | *Orbivirus* | *Reoviridae* | dsRNA | NC_027553, NC_027554, NC_027555, NC_027556, NC_027557, NC_027558, NC_027559, NC_027560, NC_027561, NC_027562 | Complete genome | ? | 1978 (Karabatsos, 1978) | *Ornithodoros* spp. (Shi et al., 2018) | Nepal (Shi et al., 2018) |
| Great Island virus | *Great Island virus* | *Orbivirus* | *Reoviridae* | dsRNA | NC_014522, NC_014523, NC_014524, NC_014525, NC_014526, NC_014527, NC_014528, NC_014529, NC_014530, NC_014531 | Complete genome | ? | 1981 (Nuttall et al., 1981) | *I. uriae* (Shi et al., 2018) | Canada (Shi et al., 2018) |
| Kemerovo virus | *Great Island virus* | *Orbivirus* | *Reoviridae* | dsRNA | MF939555, MF939556, MF939557, MF939558, MF939559, MF939560, MF939561, MF939562, MF939563, MF939564, KJ010787 | Complete genome | ? | 1970 (Libikova et al., 1970) | *I. persulcatus*, *I. ricinus* (Shi et al., 2018) | Russia, Slovakia (Shi et al., 2018) |
| Lipovnik virus | *Great Island virus* | *Orbivirus* | *Reoviridae* | dsRNA | HM543475, HM543476, HM543477 | Partial genome | ? | 1964 (Libikova et al., 1964) | *I. ricinus* (Shi et al., 2018) | Czech Republic, Slovakia (Shi et al., 2018) |
| Tribec virus | *Great Island virus* | *Orbivirus* | *Reoviridae* | dsRNA | HQ266581, HQ266582, HQ266583, HQ266584, HQ266585, HQ266586, HQ266587, HQ266588, HQ266589, HQ266590 | Partial genome | ? | 1964 (Libikova et al., 1964) | *H. punctata, I. ricinus* (Shi et al., 2018) | Belorussia, Italy, Slovakia (Shi et al., 2018) |
| St Croix River virus | *St Croix River virus* | *Orbivirus* | *Reoviridae* | dsRNA | NC_006006, NC_005998, NC_005999, NC_006000, NC_006001, NC_006002, NC_006003.1, NC_006004, NC_006005, NC_005997 | Complete genome | ? | 2001 (Attoui et al., 2001) | *I. scapularis*, *R. appendiculatus* (Shi et al., 2018) | Kenya, USA (Alberdi et al., 2012) |
| Baku virus | Unclassified | *Orbivirus* | *Reoviridae* | dsRNA | KY023349, KY023350, KY023351, KY023352, KY023353, KY023354, KY023355, KY023356, KY023357, KY023358 | Partial genome | ? | 1978 (Karabatsos, 1978) | *O. maritimus* (Shi et al., 2018) | Caspian Sea, Uzbekistan (Shi et al., 2018) |
| Matucare virus | Unclassified | *Orbivirus* | *Reoviridae* | dsRNA | HQ397640 | Partial genome | ? | 1970 (Justines et al., 1970) | *Ot. megnini (Palacios et al., 2011)* |  |
| Okhotskiy virus | Unclassified | *Orbivirus* | *Reoviridae* | dsRNA | KY023329, KY023330, KY023331, KY023332, KY023333, KY023334, KY023335, KY023336, KY023337, KY023338, KF981623, KF981624, KF981625, KF981626, KF981627, KF981628, KF981629, KF981630, KF981631, KF981632, MN830241, MN830242, MN830243, MN830244, MN830245 | Partial genome | ? | 1878 (L'vov et al., 1878) | *I. uriae* (Pettersson et al., 2020) | Northern Sweden, Okhotsk (Pettersson et al., 2020) |
| Wad Medani virus | *Wad Medani virus* | *Orbivirus* | *Reoviridae* | dsRNA | NC_027533, NC_027539, NC_027540, NC_027541, NC_027542, NC_027543, NC_027544, NC_027545, NC_027546, NC_027547 | Complete genome | ? | 1966 (Taylor et al., 1966) | *Hyalomma spp., R. sanguineus* (Shi et al., 2018) | Asia, East Africa, Jamaica (Shi et al., 2018) |
| Norway partiti-like virus 1 | Unclassified | Unclassified | *Partitiviridae* | dsRNA | MF141076 | Partial genome | ? | 2017 (Pettersson et al., 2017) | *I. ricinus* (Pettersson et al., 2017) | Norway (Pettersson et al., 2017) |
| Lonestar tick totivirus | Unclassified | Unclassified | *Totiviridae* | dsRNA | MG647775, MZ567079 | Partial genome | ? | 2018 (Tokarz et al., 2018) | *I. scapularis* (Tokarz et al., 2018) | USA (Tokarz et al., 2018) |
| Fennes virus | Unclassified | Unclassified | *Unclassified Riboviria* | dsRNA | MT025166, MT025167, MT025168, MT025169, MN830240 | Partial genome | ? | 2020 (Wille et al., 2020) | *I. uriae* (Wille et al., 2020) | Antarctica (Wille et al., 2020) |
| Shelly Beach virus | Unclassified | Unclassified | *Unclassified Riboviria* | dsRNA | MK026600, MK026601, MK026602, MK026603, MK026604, MK026605 | Partial genome | ? | 2018 (Harvey et al., 2018) | *I. holocyclus (Harvey et al., 2018)* | Australia (Harvey et al., 2018) |
| Shelly headland virus | Unclassified | Unclassified | *Unclassified Riboviria* | dsRNA | OL452144, OL452150, OL452158, OL452171, OL452185, OL452194, OL452203, OL452214, OL452222, OL452151, OL452172, OL452215, OL452223, OL452173, OL452216, OL452224, OL452174, OL452217, OL452225, OL452175, OL452226, OL452218, OL452227, OL452228, OL452176, OL452178, OL452229, OL452179, OL452219, OL452230, OL452180, OL452220, OL452231, OL452232, OL452181, OL452221, OL452233, OL452182, OL452234, OL452235, MK026577, MK026581, MK026582 | Partial genome | ? | 2018 (Harvey et al., 2018) | *I. holocyclus (Harvey et al., 2018)* | Australia (Harvey et al., 2018) |
| Lonestar tick chuvirus 1 | *Mivirus amblyommae* | *Mivirus* | *Chuviridae* | ssRNA(-) | NC_030204 | Complete genome | ? | 2018 (Tokarz et al., 2018) | *Am. americanum (Tokarz et al., 2018)* | USA (Tokarz et al., 2018) |
| Bole tick virus 3 | *Mivirus boleense* | *Mivirus* | *Chuviridae* | ssRNA(-) | NC_028259 | Complete genome | ? | 2015 (Li et al., 2015) | *Hy. Asiaticum, Hy. Aegyptium, R. turanicus, R. bursa (Li et al., 2015; Brinkmann et al., 2018; Ergunay et al., 2020)* | China, Turkey (Li et al., 2015; Brinkmann et al., 2018; Ergunay et al., 2020) |
| Changping tick virus 2 | *Mivirus changpingense* | *Mivirus* | *Chuviridae* | ssRNA(-) | NC_028260 | Complete genome | ? | 2015 (Li et al., 2015) | *Dermacentor spp., H. parva, R. sanguineus (Li et al., 2015; Temmam et al., 2019)* | China, Thailand, Turkey, Trinidad, Tobago (Li et al., 2015; Brinkmann et al., 2018; Temmam et al., 2019) |
| Changping tick virus 3 | *Mivirus dermacentoris* | *Mivirus* | *Chuviridae* | ssRNA(-) | NC_028261 | Complete genome | ? | 2015 (Li et al., 2015) | *Dermacentor spp. (Li et al., 2015)* | China (Li et al., 2015) |
| Genoa virus | *Mivirus genovaense* | *Mivirus* | *Chuviridae* | ssRNA(-) | OL452069 | Complete genome | ? | 2018 (Harvey et al., 2018) | *I. holocyclus (Harvey et al., 2018)* | Australia (Harvey et al., 2018) |
| Suffolk virus | *Mivirus suffolkense* | *Mivirus* | *Chuviridae* | ssRNA(-) | NC_028243 | Complete genome | ? | 2018 (Tokarz et al., 2018) | *I. scapularis (Tokarz et al., 2018)* | USA (Tokarz et al., 2018) |
| Wuhan tick virus 2 | *Mivirus wuhanense* | *Mivirus* | *Chuviridae* | ssRNA(-) | NC_028266 | Complete genome | ? | 2015 (Li et al., 2015) | *H. longicornis, R. microplus* (Gómez et al., 2020; Xu et al., 2021) | Brazil, China, Colombia, Guadeloupe, Thailand, Trinidad, Tobago (Gomez et al., 2020; Xu et al., 2021) |
| Wuhan mivirus | *Wuhan mivirus* | *Mivirus* | *Chuviridae* | ssRNA(-) | MZ244267 | Complete genome | ? | 2015 (Li et al., 2015) | *H. longicornis, R. microplus, R. sanguineus* (Sameroff et al., 2019) | China, Thailand, Trinidad and Tobago (Sameroff et al., 2019) |
| Tacheng tick virus 4 | *Morsusvirus argatis* | *Morsusvirus* | *Chuviridae* | ssRNA(-) | NC_028263 | Complete genome | ? | 2015 (Li et al., 2015) | *A. miniatus (Li et al., 2015)* | China (Li et al., 2015) |
| Blacklegged tick chuvirus 2 | *Nigecruvirus ixodes* | *Nigecruvirus* | *Chuviridae* | ssRNA(-) | MF360789 | Complete genome | ? | 2018 (Tokarz et al., 2018) | *I. scapularis* (Tokarz et al., 2018) | USA (Tokarz et al., 2018) |
| Amblyomma dissimile mivirus* | Unclassified | *Mivirus* | *Chuviridae* | ssRNA(-) | MZ502309.1 | Complete genome | ? | 2021 (Sameroff et al., 2021) | *Am. dissimile (Sameroff et al., 2021)* | Trinidad and Tobago (Sameroff et al., 2021) |
| Nuomin virus* | Unclassified | Unclassified | *Chuviridae* | ssRNA(-) | MW029986.1 | Partial genome | Y (Quan et al., 2020) | 2020 (Quan et al., 2020) | *I. persulcatus, I. crenulatus, H. conicinna, H. longicornis (Quan et al., 2020)* | China (Quan et al., 2020) |
| Tacheng tick virus 6 | *Tacheng arlivirus* | *Arlivirus* | *Lispiviridae* | ssRNA(-) | NC_031270 | Complete genome | ? | 2015 (Li et al., 2015) | *A. miniatus* (Li et al., 2015) | China (Li et al., 2015) |
| Grotenhout virus | *Grotenhout norwavirus* | *Norwavirus* | *Nairoviridae* | ssRNA(-) | KY700684, KY700683 | Complete genome | ? | 2017 (Vanmechelen et al., 2017) | *I. ricinus* (Vanmechelen et al., 2017) | Belgium (Vanmechelen et al., 2017) |
| Abu Hammad virus | *Abu Hammad orthonairovirus* | *Orthonairovirus* | *Nairoviridae* | ssRNA(-) | KU925434, KU925435, KU925436 | Complete genome | ? | 1976 (Darwish et al., 1976) | *A. hermanni* (Darwish et al., 1976) | Egypt (Darwish et al., 1976) |
| Abu Mina virus | *Abu Mina orthonairovirus* | *Orthonairovirus* | *Nairoviridae* | ssRNA(-) | KU925437, KU925438, KU925439 | Complete genome | ? | 1976 (Darwish et al., 1976) | *A. streptopelia* (Darwish et al., 1976) | Egypt (Darwish et al., 1976) |
| Artashat virus | *Artashat orthonairovirus* | *Orthonairovirus* | *Nairoviridae* | ssRNA(-) | NC_043440, NC_043442, NC_043441 | Complete genome | ? | 2014 (Al'khovskii et al., 2014) | *O. alactagalis* (Al'khovskii et al., 2014) | Armenia, Azerbaijan (Al'khovskii et al., 2014) |
| Avalon virus | *Avalon orthonairovirus* | *Orthonairovirus* | *Nairoviridae* | ssRNA(-) | NC_040460, NC_040459, NC_040458 | Complete genome | ? | 1976 (Main et al., 1976) | *I. uriae* (Pettersson et al., 2020) | Canada, France, Sweden (Pettersson et al., 2020) |
| Burana virus | *Burana orthonairovirus* | *Orthonairovirus* | *Nairoviridae* | ssRNA(-) | NC_043439, NC_043438, NC_043437 | Complete genome | ? | 2014 (L'vov et al., 2014) | *H. punctate* (L'vov et al., 2014) | Kyrgyzstan (L'vov et al., 2014) |
| Chim virus | *Chim orthonairovirus* | *Orthonairovirus* | *Nairoviridae* | ssRNA(-) | NC_043434, NC_043436, NC_043435 | Complete genome | ? | 2014 (L'vov et al., 2014) | *O. tartakovskyi* (L'vov et al., 2014) | Uzbekistan (L'vov et al., 2014) |
| Crimean-Congo hemorrhagic fever virus | *Crimean-Congo hemorrhagic fever orthonairovirus* | *Orthonairovirus* | *Nairoviridae* | ssRNA(-) | NC_005301, NC_005300, NC_005302 | Complete genome | Y (Garrison et al., 2020) | 1945 (Grashchenkov et al., 1945) | *Hy. asiaticum, Hy. marginatum*, *ixodid* spp., *R. rossicus* (Shi et al., 2018; Zhao et al., 2021) | Many countries in Asia and Africa; parts of Europe (e.g. Albania, Bulgaria) (Shi et al., 2018) |
| Dera Ghazi Khan virus | *Dera Ghazi Khan orthonairovirus* | *Orthonairovirus* | *Nairoviridae* | ssRNA(-) | NC_034520, NC_034510, NC_034521 | Complete genome | ? | 1970 (Begum et al., 1970) | *Hy. dromedarii* (Begum et al., 1970) | Pakistan (Begum et al., 1970) |
| Dugbe virus | *Dugbe orthonairovirus* | *Orthonairovirus* | *Nairoviridae* | ssRNA(-) | NC_004159, NC_004158, NC_004157 | Complete genome | Y (Garrison et al., 2020) | 1973 (David-West et al., 1973) | *Am. variegatum*, B. decoloratus, *Hy. truncatum*, and *R. appendiculatus* (Shi et al., 2018) | Sub-Saharan Africa (Shi et al., 2018) |
| Estero Real virus | *Estero Real orthonairovirus* | *Orthonairovirus* | *Nairoviridae* | ssRNA(-) | MH017280, MH017286, MH017274 | Complete genome | ? | 1985 (Malkova et al., 1985) | *O. tadaridae* (D Malkova et al., 1985*)* | Cuba (D Malkova et al., 1985) |
| Hazara virus | *Hazara orthonairovirus* | *Orthonairovirus* | *Nairoviridae* | ssRNA(-) | NC_038709, NC_038710, NC_038711 | Complete genome | ? | 1970 (Begum et al., 1970) | *I. redikorzevi* (Begum et al., 1970) | Japan, Pakistan (Begum et al., 1970; Matsumoto et al., 2019) |
| Huangpi tick virus 1 | *Huangpi orthonairovirus* | *Orthonairovirus* | *Nairoviridae* | ssRNA(-) | NC_031135, NC_031136, NC_031137 | Complete genome | ? | 2015 (Li et al., 2015) | *H. doenitzi, H. longicornis* (Shi et al., 2018) | China (Shi et al., 2018) |
| Caspiy virus | *Hughes orthonairovirus* | *Orthonairovirus* | *Nairoviridae* | ssRNA(-) | KP792708, KP792709, KP792710 | Complete genome | ? | 2014 (L'vov et al., 2014) | *Argasidae* (L'vov et al., 2014) | Azerbaijan (L'vov et al., 2014) |
| Farallon virus | *Hughes orthonairovirus* | *Orthonairovirus* | *Nairoviridae* | ssRNA(-) | NC_034502, NC_034494, NC_034503 | Complete genome | ? | 1967 (Radovsky et al., 1967) | *Ornithodorus* spp. (Shi et al., 2018) | USA (Shi et al., 2018) |
| Great Saltee virus | *Hughes orthonairovirus* | *Orthonairovirus* | *Nairoviridae* | ssRNA(-) | KU925467, KU925468, KU925469 | Complete genome | ? | 1976 (Keirans et al., 1976) | *O. maritimus* (Kuhn et al., 2016) | Ireland (Great Saltee Island) (Kuhn et al., 2016) |
| Hughes virus | *Hughes orthonairovirus* | *Orthonairovirus* | *Nairoviridae* | ssRNA(-) | KU925470, KU925471, KU925472 | Complete genome | ? | 1964 (Hughes et al., 1964) | *O. denmarki* (Shi et al., 2018) | Cuba, Trinidad, USA, Venezuela (Shi et al., 2018) |
| Raza virus | *Hughes orthonairovirus* | *Orthonairovirus* | *Nairoviridae* | ssRNA(-) | KU925479, KU925480, KU925481 | Complete genome | ? | 1968 (Clifford et al., 1968) | *O. denmarki* (Kuhn et al., 2016) | Mexico (Kuhn et al., 2016) |
| Issyk-kul virus | *Issyk-kul orthonairovirus* | *Orthonairovirus* | *Nairoviridae* | ssRNA(-) | KF892055, KF892056, KF892057 | Complete genome | ? | 1973 (L'vov et al, 1973) | *A. vespertlionis* (L'vov et al, 1973) | Kyrgyzstan (Atkinson et al., 2015) |
| Kasokero virus | *Kasokero orthonairovirus* | *Orthonairovirus* | *Nairoviridae* | ssRNA(-) | NC_036636, NC_029933, NC_029932 | Complete genome | ? | 1986 (Kalunda M et al., 1986) | *O. faini* (Amy et al., 2020) | Uganda (Walker et al., 2015) |
| Keterah virus | *Keterah orthonairovirus* | *Orthonairovirus* | *Nairoviridae* | ssRNA(-) | NC_034392, NC_034388, NC_034389 | Complete genome | ? | 1976 (Varma et al., 1976) | *Argasidae* (Varma et al., 1976) | Malaysia (Varma et al., 1976) |
| Kupe virus | *Kupe orthonairovirus* | *Orthonairovirus* | *Nairoviridae* | ssRNA(-) | EU257628, EU257627, EU257626 | Complete genome | ? | 2009 (Crabtree et al., 2009) | *Am. gemma, R. pulchellus* (Crabtree et al., 2009) | Kenya (Crabtree et al., 2009) |
| Meram virus | *Meram orthonairovirus* | *Orthonairovirus* | *Nairoviridae* | ssRNA(-) | MN972594, MN972595, MN972596 | Complete genome | ? | 2020 (Ergunay et al., 2020) | *Hy. aegyptium* (Ergunay et al., 2020) | Turkey (Ergunay et al., 2020) |
| Nairobi sheep disease virus | *Nairobi sheep disease orthonairovirus* | *Orthonairovirus* | *Nairoviridae* | ssRNA(-) | NC_034387, NC_034391, NC_034386 | Complete genome | Y (Yadav et al., 2011) | 1969 (Dandawate et al., 1969) | *Am. variegatum, H. longicornis, H. intermedia, R. appendiculatus, R. hemaphysaloides* (Krasteva et al., 2020) | China, Kenya, India (Krasteva et al., 2020) |
| Pacific coast tick nairovirus | *Pacific Coast orthonairovirus* | *Orthonairovirus* | *Nairoviridae* | ssRNA(-) | KU933933, KU933934, KU933935 | Complete genome | ? | 2017 (Bouquet et al., 2017) | *D. occidentalis* (Bouquet et al., 2017) | USA (Bouquet et al., 2017) |
| Punta Salinas virus | *Punta orthonairovirus* | *Orthonairovirus* | *Nairoviridae* | ssRNA(-) | KU925473, KU925474, KU925475 | Complete genome | ? | 1981 (Converse et al., 1981) | *O. amblus* (Kuhn et al., 2016) | Peru (Kuhn et al., 2016) |
| Bandia virus | *Qalyub orthonairovirus* | *Orthonairovirus* | *Nairoviridae* | ssRNA(-) | KU925446, KU925447, KU925448 | Complete genome | ? | 1967 (Bres et al., 1967) | *O. sonrai* (Kuhn et al., 2016) | Senegal (Kuhn et al., 2016) |
| Geran virus | *Qalyub orthonairovirus* | *Orthonairovirus* | *Nairoviridae* | ssRNA(-) | KP792714, KP792715, KP792716 | Complete genome | ? | 2014 (L'vov et al., 2014) | *O. asperus* (Kuhn et al., 2016) | Azerbaijan (Kuhn et al., 2016) |
| Qalyub virus | *Qalyub orthonairovirus* | *Orthonairovirus* | *Nairoviridae* | ssRNA(-) | NC_034511, NC_034522, NC_034512 | Complete genome | ? | 1952 (Kuhn et al., 2016) | *O. asperus, O. erraticus* (Walker et al., 2016; Kuhn et al., 2016) | Azerbaijan, Egypt, Senegal (Kuhn et al., 2016; Walker et al., 2016) |
| Clo Mor virus | *Sakhalin orthonairovirus* | *Orthonairovirus* | *Nairoviridae* | ssRNA(-) | NC_034561, NC_034554, NC_034562 | Complete genome | ? | 1976 (Main et al., 1976) | *A. cooleyi, I. uriae* (Walker et al., 2016) | UK, USA (Walker et al., 2016) |
| Sakhalin virus | *Sakhalin orthonairovirus* | *Orthonairovirus* | *Nairoviridae* | ssRNA(-) | KU925482, KU925483, KU925484 | Complete genome | ? | 1972 (L'vov et al., 1972) | *I. uriae* (Kuhn et al., 2016) | Antarctica, Australia, Russia, USA (L'vov et al., 2014; Kuhn et al., 2016; Pettersson et al., 2020) |
| Taggert virus | *Sakhalin orthonairovirus* | *Orthonairovirus* | *Nairoviridae* | ssRNA(-) | KU925491, KU925492, KU925493 | Complete genome | ? | 1985 (Doherty et al., 1985) | *I. uriae* (Pettersson et al., 2020; Wille et al., 2020) | Antarctica, Australia (Pettersson et al., 2020; Wille et al., 2020) |
| Tillamook virus | *Sakhalin orthonairovirus* | *Orthonairovirus* | *Nairoviridae* | ssRNA(-) | KU925494, KU925495, KU925496 | Complete genome | ? | 1973 (Thomas et al., 1973) | *I. uriae* (Kuhn et al., 2016) | USA (Kuhn et al., 2016) |
| Soldado virus | *Soldado orthonairovirus* | *Orthonairovirus* | *Nairoviridae* | ssRNA(-) | KU925488, KU925489, KU925490 | Complete genome | ? | 1973 (Jonkers et al., 1973) | *O. maritimus* (Shi et al., 2018) | France, Great Britain, Indian Ocean, North Wales, Seychelles (Shi et al., 2018) |
| Tacheng tick virus 1 | *Tacheng orthonairovirus* | *Orthonairovirus* | *Nairoviridae* | ssRNA(-) | NC_031284, NC_031285, NC_031286 | Complete genome | Y | 2015 (Li et al., 2015) | *D. marginatus, D. nuttalli, D. silvarum, Hy. asiaticum* (Shi et al., 2018) | China, Kazakhstan (Shi et al., 2018) |
| Tamdy virus | *Tamdy orthonairovirus* | *Orthonairovirus* | *Nairoviridae* | ssRNA(-) | MN792651, MN792652, MN792653 | Complete genome | Y (Moming et al., 2021) | 2014 (L'vov et al., 2014) | *Hy. asiaticum (L'vov et al., 1984; Moming et al., 2021)* | China, Uzbekistan (Moming et al., 2021) |
| Tofla virus | *Tofla orthonairovirus* | *Orthonairovirus* | *Nairoviridae* | ssRNA(-) | NC_029124, NC_029123, NC_029122 | Complete genome | ? | 2016 (Shimada et al., 2016) | *H. flava, H. formsensis* (Shimada et al., 2016) | Japan (Shimada et al., 2016) |
| Tunis virus | *Tunis orthonairovirus* | *Orthonairovirus* | *Nairoviridae* | ssRNA(-) | NC_040770, NC_040771, NC_040772 | Complete genome | ? | 1994 (Chastel et al., 1994) | *A. hermanni* (Chastel et al., 1994) | Tunisia (Chastel et al., 1994) |
| Vinegar Hill virus | *Vinegar Hill orthonairovirus* | *Orthonairovirus* | *Nairoviridae* | ssRNA(-) | MF176881, MF176882, MF176883 | Complete genome | ? | 2017 (Gauci et al., 2017) | *A. robertsi* (Gauci et al., 2017) | Australia (Gauci et al., 2017) |
| Wenzhou tick virus | *Wenzhou orthonairovirus* | *Orthonairovirus* | *Nairoviridae* | ssRNA(-) | NC_031291, NC_031288, NC_031289 | Complete genome | ? | 2015 (Li et al., 2015) | *H. hystricis* (Li et al., 2015) | China (Li et al., 2015) |
| South Bay virus | *South Bay sabavirus* | *Sabavirus* | *Nairoviridae* | ssRNA(-) | KJ746878, KJ746877 | Complete genome | ? | 2014 (Tokarz et al., 2014) | *D. nuttalli, H. concinna, H. longicornis, I. persulcatus, I. scapularis (Tokarz et al., 2014; Zhao et al., 2021)* | China, USA (Tokarz et al., 2014; Zhao et al., 2021) |
| Midway virus | *Midway nyavirus* | *Nyavirus* | *Nyamiviridae* | ssRNA(-) | NC_012702 | Complete genome | ? | 1982 (Takahashi et al., 1982) | *Ornithodoros* spp. (Shi et al., 2018) | Central Pacific, Japan (Shi et al., 2018) |
| Nyamanini virus | *Nyamanini nyavirus* | *Nyavirus* | *Nyamiviridae* | ssRNA(-) | NC_012703 | Complete genome | ? | 1966 (Taylor et al., 1966) | *A. arborerus, A. walkerae* (Shi et al., 2018) | Egypt, India, Nigeria, South Africa, Thailand (Shi et al., 2018) |
| Sierra Nevada virus | *Sierra Nevada nyavirus* | *Nyavirus* | *Nyamiviridae* | ssRNA(-) | NC_024376 | Complete genome | ? | 2014 (Rogers et al., 2014) | *O. coriaceus* (Rogers et al., 2014) | USA (Rogers et al., 2014) |
| Johnston Atoll virus | *Johnston Atoll quaranjavirus* | *Quaranjavirus* | *Orthomyxoviridae* | ssRNA(-) | NC_052925, NC_052926, NC_052927, NC_052928, NC_052929, NC_052930, NC_052931, FJ861696, FJ861697 | Complete genome | ? | 2009 (Presti et al., 2009) | *O. capensis* (Shi et al., 2018) | Australia, Central Pacific, New Zealand and Hawaii (Shi et al., 2018) |
| Quaranfil virus | *Quaranfil quaranjavirus* | *Quaranjavirus* | *Orthomyxoviridae* | ssRNA(-) | NC_038817, NC_038818, NC_038819, NC_038820, NC_038821, NC_038822 | Complete genome | Y (Mourya et al., 2019) | 1966 (Taylor et al., 1966) | *A. arboreus* (Shi et al., 2018) | Afghanistan, Egypt, Iraq, Kuwait, Nigeria, South Africa, Yemen and Iran (Shi et al., 2018) |
| Granville quaranjavirus* | *Granville quaranjavirus* | *Quaranjavirus* | *Orthomyxoviridae* | ssRNA(-) | MZ502307, MZ502306, MZ502305, MZ502304, MZ502308 | Partial genome | ? | 2021 (Sameroff et al., 2021) | *Am. dissimile (Sameroff et al., 2021)* | Trinidad and Tobago (Sameroff et al., 2021) |
| Dhori virus | *Dhori thogotovirus* | *Thogotovirus* | *Orthomyxoviridae* | ssRNA(-) | NC_034254, NC_034255, NC_034256, NC_034261, NC_034262, NC_034263 | Complete genome | Y (Kosoy et al., 2015) | 1973 (Anderson et al., 1973) | *Hyalomma* spp. (Shi et al., 2018) | China, Egypt, India, Kyrgyzstan, Russia (Shi et al., 2018) |
| Jos virus | *Thogoto thogotovirus* | *Thogotovirus* | *Orthomyxoviridae* | ssRNA(-) | HM627170, HM627171, HM627172, HM627173, HM627174, HM627175 | Complete genome | ? | 1974 (Lee et al., 1974) | *Amblyomma* spp., *Rhipicephalus* spp. (Shi et al., 2018) | Central Africa Republic, Guinea, Ivory Coast, Nigeria, Senegal (Shi et al., 2018) |
| Thogoto virus | *Thogoto thogotovirus* | *Thogotovirus* | *Orthomyxoviridae* | ssRNA(-) | NC_006495, NC_006496, NC_006504, NC_006506, NC_006507, NC_006508 | Complete genome | Y (Kosoy et al., 2015) | 1965 (Haig et al., 1965) | *Am. Variegatum, Boophilus spp., Hyalomma spp., Rhipicephalus* spp.(Shi et al., 2018) | Central and East Africa, Southern Europe, Southern Portugal (Shi et al., 2018) |
| Ngari virus | *Bunyamwera orthobunyavirus* | *Orthobunyavirus* | *Peribunyaviridae* | ssRNA(-) | KC608152, KC608153, KC608154 | Complete genome | Y (Groseth et al., 2012) | 1996 (Zeller et al., 1996) | *Am. variegatum, R. pulchellus* (Groseth et al., 2012) | Guinea, Kenya, Mauritania, Senegal, Somalia, Sudan (Groseth et al., 2012; Makenov et al., 2021) |
| Matruh virus | *Matruh orthobunyavirus* | *Orthobunyavirus* | *Peribunyaviridae* | ssRNA(-) | KP792693, KP792692, KP792691 | Complete genome | ? | 1973 (Balducci et al., 1973) | *Hy. Marginatum* (Balducci et al., 1973) | Italy (Balducci et al., 1973) |
| Bahig virus | *Tete orthobunyavirus* | *Orthobunyavirus* | *Peribunyaviridae* | ssRNA(-) | KP792654, KP792653, KP792652 | Complete genome | ? | 1973 (Balducci et al., 1973) | *Hy. Marginatum* (Balducci et al., 1973) | Italy (Balducci et al., 1973) |
| Bhanja virus | *Bhanja bandavirus* | *Bandavirus* | *Phenuiviridae* | ssRNA(-) | NC_027140, NC_027141, NC_027142 | Complete genome | Y (Matsuno et al., 2013) | 1977 (Vesenjak-Hirjan et al., 1977) | *H. intermedia, R. decoloratus* (Matsuno et al., 2013) | Africa, Asia, Southern Europe, USA (Shi et al., 2018) |
| Heartland virus | *Heartland bandavirus* | *Bandavirus* | *Phenuiviridae* | ssRNA(-) | NC_024494, NC_024495, NC_024496 | Complete genome | Y (McMullan et al., 2012) | 2012 (McMullan et al., 2012) | *Am. americanum (Mansfield et al., 2017)* | USA (Mansfield et al., 2017) |
| Hunter Island virus | *Hunter Island bandavirus* | *Bandavirus* | *Phenuiviridae* | ssRNA(-) | NC_027715, NC_027716, NC_027717 | Complete genome | ? | 2014 (Wang et al., 2014) | *I. eudyptidis* (Shi et al., 2018) | Australia (Shi et al., 2018) |
| lone star virus | *Lone star bandavirus* | *Bandavirus* | *Phenuiviridae* | ssRNA(-) | NC_021242, NC_021243, NC_021244 | Complete genome | ? | 1969 (Kokernot et al., 1969) | *Am. Americanum* (Shi et al., 2018) | USA (Shi et al., 2018) |
| Severe fever with thrombocytopenia syndrome virus | *Dabie bandavirus* | *Bandavirus* | *Phenuiviridae* | ssRNA(-) | NC_043450, NC_043451, NC_043452 | Complete genome | Y (Yu et al., 2011) | 2011 (Yu et al., 2011) | *Dermacentor spp., Haemaphysalis spp., I. persulcatus, Rhipicephalus spp. (Mansfield et al., 2017; Zhao et al., 2021)* | East Asia, North America (Mansfield et al., 2017; Zhao et al., 2021) |
| Norway phlebovirus 1 | *Norway ixovirus* | *Ixovirus* | *Phenuiviridae* | ssRNA(-) | NC_055433, NC_055434 | Complete genome | ? | 2017 (Pettersson et al., 2017) | *I. ricinus* (Pettersson et al., 2017) | Norway (Pettersson et al., 2017) |
| Blacklegged tick phlebovirus 3 | *Scapularis ixovirus* | *Ixovirus* | *Phenuiviridae* | ssRNA(-) | NC_055431, NC_055432 | Complete genome | ? | 2018 (Tokarz et al., 2018) | *I. scapularis (Tokarz et al., 2018)* | USA (Tokarz et al., 2018) |
| Laurel Lake virus | *Laurel Lake laulavirus* | *Laulavirus* | *Phenuiviridae* | ssRNA(-) | NC_043679, NC_043680, NC_043681 | Complete genome | ? | 2018 (Tokarz et al., 2018) | *I. scapularis (Tokarz et al., 2018)* | USA (Tokarz et al., 2018) |
| Dabieshan tick virus | *Dabieshan uukuvirus* | *Uukuvirus* | *Phenuiviridae* | ssRNA(-) | MW721887, MW721893 | Complete genome | ? | 2015 (Li et al., 2015) | *H. longicornis, I. persulcatus, R. microplus* (Wang et al., 2021; Zhao et al., 2021) | China (Wang et al., 2021; Zhao et al., 2021) |
| Kaisodi virus | *Kaisodi uukuvirus* | *Uukuvirus* | *Phenuiviridae* | ssRNA(-) | NC_040492, NC_040493, NC_040494 | Complete genome | ? | 1966 (Bhatt et al., 1966) | *H. spinigera* (Shi et al., 2018) | South India (Shi et al., 2018) |
| Lihan tick virus | *Lihan uukuvirus* | *Uukuvirus* | *Phenuiviridae* | ssRNA(-) | NC_055423, NC_055424 | Complete genome | ? | 2015 (Li et al., 2015) | *D. nitens, Hy. marginatum, R. microplus, R. sanguineus* (Temmam et al., 2019) | Brazil, China, USA, Guadeloupe, Thailand, Turkey, Trinidad. Tobago (Temmam et al., 2019; Gondard et al., 2020) |
| Silverwater virus | *Silverwater uukuvirus* | *Uukuvirus* | *Phenuiviridae* | ssRNA(-) | NC_055369, NC_055370, NC_055371 | Complete genome | ? | 1971 (Hoff et al., 1971) | *H. leporispalustris* (Shi et al., 2018) | Canada, USA (Shi et al., 2018) |
| Tacheng tick virus 2 | *Tacheng uukuvirus* | *Uukuvirus* | *Phenuiviridae* | ssRNA(-) | NC_055425, NC_055426 | Complete genome | Y (Dong et al., 2021) | 2015 (Li et al., 2015) | *D. marginatus, D. reticulatus, R. sanguineus, H. marginatum* (Li et al., 2015; Brinkmann et al., 2018; Bratuleanu et al., 2021) | China, Kazakhstan, Romania, Tailand, Turkey (Li et al., 2015; Brinkmann et al., 2018; Temmam et al., 2019; Bratuleanu et al., 2021) |
| Uukuniemi virus | *Uukuniemi uukuvirus* | *Uukuvirus* | *Phenuiviridae* | ssRNA(-) | NC_005214, NC_005220, NC_005221 | Complete genome | ? | 2013 (Palacios et al., 2013) | *I. ricinus, I. uriae* (Matsuno et al., 2015) | Czech Republic, Sweden, UK (Matsuno et al., 2015) |
| Yongjia tick virus 1 | *Yongjia uukuvirus* | *Uukuvirus* | *Phenuiviridae* | ssRNA(-) | NC_055427, NC_055428 | Complete genome | ? | 2015 (Li et al., 2015) | *H. hystricis* (Li et al., 2015) | China (Li et al., 2015) |
| Blanchseco virus | *Blanchseco alpharicinrhavirus* | *Alpharicinrhavirus* | *Rhabdoviridae* | ssRNA(-) | MN025503 | Complete genome | ? | 2019 (Sameroff et al., 2019) | *A. ovale* (Sameroff et al., 2019) | Trinidad and Tobago (Sameroff et al., 2019) |
| Bole tick virus 2 | *Bole alpharicinrhavirus* | *Alpharicinrhavirus* | *Rhabdoviridae* | ssRNA(-) | NC_031079 | Complete genome | ? | 2015 (Li et al., 2015) | *Hy. Asiaticum* (Li et al., 2015) | China (Li et al., 2015) |
| Wuhan tick virus 1 | *Wuhan alpharicinrhavirus* | *Alpharicinrhavirus* | *Rhabdoviridae* | ssRNA(-) | NC_031304 | Complete genome | ? | 2015 (Li et al., 2015) | *D. marginatus, R. microplus, R. annulatus, H. longicornis* (Temmam et al., 2019; Xu et al., 2021) | China, Turkey (Xu et al., 2021) |
| New Kent County virus | *Kent ephemerovirus* | *Ephemerovirus* | *Rhabdoviridae* | ssRNA(-) | MF615270 | Complete genome | ? | 2018 (Tokarz et al., 2018) | *I. scapularis (Tokarz et al., 2018)* | USA (Tokarz et al., 2018) |
| Barur virus | *Barur ledantevirus* | *Ledantevirus* | *Rhabdoviridae* | ssRNA(-) | NC_034535 | Complete genome | ? | 1981 (Butenko et al., 1981) | *Hy. Intermedia* (Shi et al., 2018) | India, Kenya, Somalia (Shi et al., 2018) |
| Kolente virus | *Kolente ledantevirus* | *Ledantevirus* | *Rhabdoviridae* | ssRNA(-) | NC_025342 | Complete genome | ? | 2013 (Ghedin et al., 2013) | *Am. Variegatum* (Shi et al., 2018) | Guinea (Shi et al., 2018) |
| Yongjia tick virus 2 | *Yongjia ledantevirus* | *Ledantevirus* | *Rhabdoviridae* | ssRNA(-) | NC_031305 | Complete genome | ? | 2015 (Li et al., 2015) | *H. hystricis* (Shi et al., 2018) | China (Shi et al., 2018) |
| Connecticut virus | *Connecticut sawgrhavirus* | *Sawgrhavirus* | *Rhabdoviridae* | ssRNA(-) | KM205020 | Complete genome | ? | 1978 (Ritter et al., 1978) | *I. dentatus* (Shi et al., 2018) | USA (Shi et al., 2018) |
| Long Island tick rhabdovirus | *Island sawgrhavirus* | *Sawgrhavirus* | *Rhabdoviridae* | ssRNA(-) | NC_025340 | Complete genome | ? | 2014 (Tokarz et al., 2014) | *Am. americanum (Tokarz et al., 2014)* | USA (Tokarz et al., 2014) |
| New Minto virus | *Minto sawgrhavirus* | *Sawgrhavirus* | *Rhabdoviridae* | ssRNA(-) | NC_055457 | Complete genome | ? | 1970 (Sather et al., 1970) | *H. leporispalustris* (Shi et al., 2018) | USA (Shi et al., 2018) |
| Sawgrass virus | *Sawgrass sawgrhavirus* | *Sawgrhavirus* | *Rhabdoviridae* | ssRNA(-) | NC_055461 | Complete genome | ? | 1970 (Sather et al., 1970) | *D. variabilis, H. leporispalustris* (Shi et al., 2018) | USA (Shi et al., 2018) |
| Isfahan virus | *Isfahan vesiculovirus* | *Vesiculovirus* | *Rhabdoviridae* | ssRNA(-) | NC_020806 | Complete genome | ? | 1977 (Tesh et al., 1977) | *Hy. Asiaticum* (Shi et al., 2018) | Parts of Asia, Turkmenistan (Shi et al., 2018) |
| Zahedan rhabdovirus | *Zahedan zarhavirus* | *Zarhavirus* | *Rhabdoviridae* | ssRNA(-) | NC_040664 | Complete genome | ? | 2015 (Dilcher et al., 2015) | *Hy. Anatolicum* (Shi et al., 2018) | Iran (Shi et al., 2018) |
| Beiji nairovirus | Unclassified | *Nairovirus* | *Nairoviridae* | ssRNA(-) | MW315106, MW315112 | Partial genome | Y (Wang et al., 2021) | 2019 (Meng et al., 2019) | *I. crenulatus, I. persulcatus* (Wang et al., 2021) | China (Wang et al., 2021) |
| Nayun tick nairovirus | Unclassified | *Nairovirus* | *Nairoviridae* | ssRNA(-) | MW561151, KP141756, KP141755 | Partial genome | ? | 2015 (Xia et al., 2015) | *R. sanguineus* (Xia et al., 2015; Bratuleanu et al., 2021) | China, Romania (Xia et al., 2015; Bratuleanu et al., 2021) |
| Norway nairovirus 1 | Unclassified | *Nairovirus* | *Nairoviridae* | ssRNA(-) | MF141045, MF141044 | Partial genome | ? | 2017 (Pettersson et al., 2017) | *I. ricinus* (Pettersson et al., 2017) | Northern Europe (Pettersson et al., 2017) |
| Paramushir virus | Unclassified | *Nairovirus* | *Nairoviridae* | ssRNA(-) | MH638289, MH124638, MH124637, MH124636, MH124635, MH124634, KF801657, KP792719, KP792718, KP792717 | Partial genome | ? | 1976 (L'vov et al., 1976) | *I. signatus, I. uriae* (L'vov et al., 1976) | Russia (Safonova et al., 2015) |
| Rondonia orthonairovirus | Unclassified | *Orthonairovirus* | *Nairoviridae* | ssRNA(-) | MN560621, MN560622, MN560623, MN560624, MN560625, MN560626, MN560627, MN560628, MN560629 | Partial genome | ? | 2019 (Blomstrom et al., 2019) | *Antricola* (Blomstrom et al., 2019) | Brazil (Blomstrom et al., 2019) |
| Songling virus | Unclassified | *Orthonairovirus* | *Nairoviridae* | ssRNA(-) | MT328780, MT328779, MT328778 | Partial genome | Y (Ma et al., 2021) | 2021 (Ma et al., 2021) | *H. longicornis* (Ma et al., 2021) | China (Ma et al., 2021) |
| Yezo virus | Unclassified | *Orthonairovirus* | *Nairoviridae* | ssRNA(-) | LC621360, LC621359, LC621358 | Partial genome | Y (Kodama et al., 2021) | 2021 (Kodama et al., 2021) | *H. megaspinosa* (Kodama et al., 2021) | Japan (Kodama et al., 2021) |
| Zambezi tick virus 1 | Unclassified | *Quarjavirus* | *Orthomyxoviridae* | ssRNA(-) | MH267793 | Partial genome | ? | 2018 (Cholleti et al., 2018) | *Rhipicephalus* (Cholleti et al., 2018) | Mozambique (Cholleti et al., 2018) |
| Aransas Bay virus | Unclassified | *Thogotovirus* | *Orthomyxoviridae* | ssRNA(-) | KC506162, KC506163, KC506164, KC506165, KC506166, KC506167 | Partial genome | ? | 1979 (Yunker et al., 1979) | *Ornithodoros* spp. (Briese et al., 2014) | North America (Briese et al., 2014) |
| Bourbon virus | Unclassified | *Thogotovirus* | *Orthomyxoviridae* | ssRNA(-) | KU708253, KU708254, KU708255, MH880287, MH880288, MH880289, MH880290, MH880291, MH880292 | Complete genome | Y (Kosoy et al., 2015) | 2015 (Kosoy et al., 2015) | *Am. Americanum* (Savage et al., 2017) | USA (Kosoy et al., 2015) |
| Oz virus | Unclassified | *Thogotovirus* | *Orthomyxoviridae* | ssRNA(-) | NC_040730, NC_040731, NC_040732, NC_040733, NC_040734, NC_040735 | Complete genome | ? | 2018 (Ejiri et al., 2018) | *Am. testudinarium* (Ejiri et al., 2018) | Japan (Ejiri et al., 2018) |
| Thailand tick thogotovirus | Unclassified | *Thogotovirus* | *Orthomyxoviridae* | ssRNA(-) | MN095539, MN095540, MN095541, MN095542, MN095543, MN095544 | Complete genome | ? | 2019 (Temmam et al., 2019) | *R. microplus* (Temmam et al., 2019) | Tailand (Temmam et al., 2019) |
| Upolu virus | Unclassified | *Thogotovirus* | *Orthomyxoviridae* | ssRNA(-) | KC506156, KC506157, KC506158, KC506159, KC506160, KC506161 | Complete genome | ? | 1968 (Doherty et al., 1968) | *O. capensis* (Briese et al., 2014) | Australia (Briese et al., 2014) |
| American dog tick phlebovirus | Unclassified | *Phlebovirus* | *Phenuiviridae* | ssRNA(-) | KM048312, KM048311 | Complete genome | ? | 2014 (Tokarz et al., 2014) | *D. variabilis (Tokarz et al., 2014)* | USA (Tokarz et al., 2014) |
| Blacklegged tick phlebovirus 1 | Unclassified | *Phlebovirus* | *Phenuiviridae* | ssRNA(-) | KX184200, KX184201 | Complete genome | ? | 2014 (Tokarz et al., 2014) | *I. scapularis (Tokarz et al., 2014)* | USA (Tokarz et al., 2014) |
| Blacklegged tick phlebovirus 2 | Unclassified | *Phlebovirus* | *Phenuiviridae* | ssRNA(-) | KM589357, KM589354 | Partial genome | ? | 2014 (Tokarz et al., 2014) | *I. scapularis (Tokarz et al., 2014)* | USA (Tokarz et al., 2014) |
| Bole tick virus 1 | Unclassified | *Phlebovirus* | *Phenuiviridae* | ssRNA(-) | MZ244253, MZ244252, KM817731, MH916622, MH916621, MH916620, MH916619, MH688504, MH836620, MH688505, MH688503, MH688501, MH688578, MH688577, MH688576, MH688575, MH688574, MH688573, MH688572, MH688571 | Partial genome | ? | 2015 (Li et al., 2015) | *Hy. Asiaticum* (Li et al., 2015) | China (Li et al., 2015) |
| Brown dog tick phlebovirus 1 | Unclassified | *Phlebovirus* | *Phenuiviridae* | ssRNA(-) | MZ244261, MZ244260, MZ244259, MZ244256, MN025507, MN025506 | Partial genome | ? | 2019 (Sameroff et al., 2019) | *R. sanguineus R. microplus* (Sameroff et al., 2019) | China, Trinidad and Tobago (Sameroff et al., 2019) |
| Brown dog tick phlebovirus 2 | Unclassified | *Phlebovirus* | *Phenuiviridae* | ssRNA(-) | MW561139, MW561138, MW561137, MW561136, MN025509, MN025508 | Partial genome | ? | 2019 (Sameroff et al., 2019) | *R. microplus, R. sanguineus* (Sameroff et al., 2019) | China, Trinidad and Tobago (Sameroff et al., 2019) |
| Changping tick virus 1 | Unclassified | *Phlebovirus* | *Phenuiviridae* | ssRNA(-) | MW561146, MW561145, MW561144, MW561143, MW561142, MW561141, MW561140, KM817732, KM817665 | Partial genome | ? | 2015 (Li et al., 2015) | *Dermacentor* spp. (Li et al., 2015) | China, Romania (Matsuno et al., 2013; Li et al., 2015) |
| Lanjan virus | Unclassified | *Phlebovirus* | *Phenuiviridae* | ssRNA(-) | KM370974 | Partial genome | ? | 1967 (Tan et al., 1967) | *D. auratus* (Shi et al., 2018) | Malaysia (Shi et al., 2018) |
| Palma virus | Unclassified | *Phlebovirus* | *Phenuiviridae* | ssRNA(-) | JX961630, JX961629, JX961628 | Partial genome | ? | 1994 (Filipe et al., 1994) | *H. punctate* (Shi et al., 2018) | Portugal (Shi et al., 2018) |
| Rhipicephalus associated phlebovirus 1 | Unclassified | *Phlebovirus* | *Phenuiviridae* | ssRNA(-) | MZ244255, MZ244254, MH814976, MH814975 | Partial genome | ? | 2021 (Shi et al., 2021) | *R. microplus* (Shi et al., 2021) | China (Shi et al., 2021) |
| Tick phlebovirus Anatolia 1 | Unclassified | *Phlebovirus* | *Phenuiviridae* | ssRNA(-) | MG764522, MG764521 | Partial genome | ? | 2018 (Brinkmann et al., 2018) | *R. sanguineus* (Brinkmann et al., 2018) | Turkey (Brinkmann et al., 2018) |
| Neke harbour virus | Unclassified | Unclassified | *Orthomyxoviridae* | ssRNA(-) | MN830238 | Partial genome | ? | 2020 (Pettersson et al., 2020) | *I. uriae* (Pettersson et al., 2020) | Antarctica (Pettersson et al., 2020) |
| Uumaja virus | Unclassified | Unclassified | *Orthomyxoviridae* | ssRNA(-) | MN830237 | Partial genome | ? | 2020 (Pettersson et al., 2020) | *I. uriae* (Pettersson et al., 2020) | Northern Sweden (Pettersson et al., 2020) |
| Khasan virus | Unclassified | Unclassified | *Peribunyaviridae* | ssRNA(-) | KF892046, KF892047, KF892048 | Partial genome | ? | 1978 (L'vov et al., 1978) | *H. longgicornis* (Shi et al., 2018) | Russia (Shi et al., 2018) |
| American dog tick rhabdovirus 2 | Unclassified | Unclassified | *Rhabdoviridae* | ssRNA(-) | MF962659, MF962661 | Partial genome | ? | 2018 (Tokarz et al., 2018) | *I. scapularis (Tokarz et al., 2018)* | USA (Tokarz et al., 2018) |
| Blacklegged tick rhabdovirus 1 | Unclassified | Unclassified | *Rhabdoviridae* | ssRNA(-) | MF360790 | Complete genome | ? | 2018 (Tokarz et al., 2018) | *I. scapularis (Tokarz et al., 2018)* | USA (Tokarz et al., 2018) |
| Dog tick rhabdovirus 1 | Unclassified | Unclassified | *Rhabdoviridae* | ssRNA(-) | MF360791 | Partial genome | ? | 2018 (Tokarz et al., 2018) | *I. scapularis (Tokarz et al., 2018)* | USA (Tokarz et al., 2018) |
| Huangpi tick virus 3 | Unclassified | Unclassified | *Rhabdoviridae* | ssRNA(-) | NC_031083 | Complete genome | ? | 2015 (Li et al., 2015) | *H. doenitzi* (Li et al., 2015) | China (Li et al., 2015) |
| Nayun tick rhabdovirus | Unclassified | Unclassified | *Rhabdoviridae* | ssRNA(-) | KP141757 | Partial genome | ? | 2015 (Xia et al., 2015) | *Rhipicephalus* spp. (Xia et al., 2015) | China (Xia et al., 2015) |
| Rhipicephalus associated rhabdo-like virus | Unclassified | Unclassified | *Rhabdoviridae* | ssRNA(-) | MH814974 | Partial genome | ? | 2021 (Shi et al., 2021) | *H. longicornis, R. microplus* (Shi et al., 2021) | China (Shi et al., 2021) |
| Tacheng tick virus 3 | Unclassified | Unclassified | *Rhabdoviridae* | ssRNA(-) | NC_031268 | Complete genome | ? | 2015 (Li et al., 2015) | *D. marginatus* (Li et al., 2015) | China (Li et al., 2015) |
| Tacheng tick virus 7 | Unclassified | Unclassified | *Rhabdoviridae* | ssRNA(-) | NC_031272 | Complete genome | ? | 2015 (Li et al., 2015) | *A. miniatus* (Li et al., 2015) | China (Li et al., 2015) |
| Taishun tick virus | Unclassified | Unclassified | *Rhabdoviridae* | ssRNA(-) | NC_031273 | Complete genome | ? | 2015 (Li et al., 2015) | *H. hystricis* (Li et al., 2015) | China, Russia (Li et al., 2015) |
| Deer tick mononegavirales-like virus | Unclassified | Unclassified | Unclassified | ssRNA(-) | MG880117, KM048317, KJ746903 | Partial genome | ? | 2014 (Tokarz et al., 2014) | *D. nuttalli, D. silvarum, H. concinna, I. persulcatus, I. scapularis (Tokarz et al., 2014; Zhao et al., 2021)* | China, USA (Tokarz et al., 2014; Zhao et al., 2021) |
| Bonden virus | Unclassified | Unclassified | *Unclassified Bunyavirales* | ssRNA(-) | MN830227 | Partial genome | ? | 2020 (Pettersson et al., 2020) | *I. uriae* (Pettersson et al., 2020) | Northern Sweden (Pettersson et al., 2020) |
| Bronnoya virus | Unclassified | Unclassified | *Unclassified Bunyavirales* | ssRNA(-) | MF141063, MF141062 | Partial genome | ? | 2017 (Pettersson et al., 2017) | *I. ricinus* (Pettersson et al., 2017) | Norway (Pettersson et al., 2017) |
| Ubmeje virus | Unclassified | Unclassified | *Unclassified Bunyavirales* | ssRNA(-) | MN830232 | Partial genome | ? | 2020 (Pettersson et al., 2020) | *I. uriae* (Pettersson et al., 2020) | Northern Sweden (Pettersson et al., 2020) |
| Norway mononegavirus 1 | Unclassified | Unclassified | *Unclassified Mononegavirales* | ssRNA(-) | MF141072 | Partial genome | ? | 2017 (Pettersson et al., 2017) | *I. ricinus* (Pettersson et al., 2017) | Norway (Pettersson et al., 2017) |
| Tick mononegavirus | Unclassified | Unclassified | *Unclassified Mononegavirales* | ssRNA(-) | MZ244310, MH835441, MH835440, MH835439, MH835438 | Partial genome | ? | 2018 (Yang et al., 2018) | *I. scapularis, H. longicornis* (Yang et al., 2018) | China (Yang et al., 2018) |
| Umea virus | Unclassified | Unclassified | *Unclassified Mononegavirales* | ssRNA(-) | MN830236 | Partial genome | ? | 2020 (Pettersson et al., 2020) | *I. uriae* (Pettersson et al., 2020) | Sweden (Pettersson et al., 2020) |
| Canne point virus | Unclassified | Unclassified | *Unclassified Riboviria* | ssRNA(-) | MK026566 | Partial genome | ? | 2018 (Harvey et al., 2018) | *Am. moreliae (Harvey et al., 2018)* | Australia (Harvey et al., 2018) |
| Fairlight virus | Unclassified | Unclassified | *Unclassified Riboviria* | ssRNA(-) | MK026565 | Partial genome | ? | 2018 (Harvey et al., 2018) | *Am. moreliae (Harvey et al., 2018)* | Australia (Harvey et al., 2018) |
| Manly virus | Unclassified | Unclassified | *Unclassified Riboviria* | ssRNA(-) | MK026564 | Partial genome | ? | 2018 (Harvey et al., 2018) | *Am. moreliae (Harvey et al., 2018)* | Australia (Harvey et al., 2018) |
| Messner virus | Unclassified | Unclassified | *Unclassified Riboviria* | ssRNA(-) | MT025174 | Partial genome | ? | 2020 (Wille et al., 2020) | *I. uriae* (Wille et al., 2020) | Antarctica (Wille et al., 2020) |
| North Shore virus | Unclassified | Unclassified | *Unclassified Riboviria* | ssRNA(-) | OL452126 | Partial genome | ? | 2018 (Harvey et al., 2018) | *I. holocyclus (Harvey et al., 2018)* | Australia (Harvey et al., 2018) |
| Old quarry swamp virus | Unclassified | Unclassified | *Unclassified Riboviria* | ssRNA(-) | MK026595, MK026596, MK026597, MK026598, MK026599 | Partial genome | ? | 2018 (Harvey et al., 2018) | *I. holocyclus (Harvey et al., 2018)* | Australia (Harvey et al., 2018) |
| Piguzov virus | Unclassified | Unclassified | *Unclassified Riboviria* | ssRNA(-) | MT025176, MT025177 | Partial genome | ? | 2020 (Pettersson et al., 2020) | *I. uriae* (Pettersson et al., 2020) | Antarctica (Pettersson et al., 2020) |
| Quarantine Head virus | Unclassified | Unclassified | *Unclassified Riboviria* | ssRNA(-) | MK026568 | Partial genome | ? | 2018 (Harvey et al., 2018) | *Am. moreliae (Harvey et al., 2018)* | Australia (Harvey et al., 2018) |
| Ronne virus | Unclassified | Unclassified | *Unclassified Riboviria* | ssRNA(-) | MT025165, MT025164 | Partial genome | ? | 2020 (Pettersson et al., 2020; Wille et al., 2020) | *I. uriae* (Pettersson et al., 2020; Wille et al., 2020) | Antarctica, Northern Sweden (Pettersson et al., 2020; Wille et al., 2020) |
| Tacheng tick virus 5 | Unclassified | Unclassified | *Unclassified Riboviria* | ssRNA(-) | NC_028264 | Complete genome | ? | 2015 (Li et al., 2015) | *D. marginatus (Li et al., 2015)* | China, Kazakhstan (Li et al., 2015) |
| Timbillica virus | Unclassified | Unclassified | *Unclassified Riboviria* | ssRNA(-) | MK026589, MK026590 | Partial genome | ? | 2018 (Harvey et al., 2018) | *I. holocyclus (Harvey et al., 2018)* | Australia (Harvey et al., 2018) |
| Gadgets Gully virus | *Gadgets Gully virus* | *Flavivirus* | *Flaviviridae* | ssRNA(+) | NC_033723 | Complete genome | ? | 1985 (St St George et al., 1985) | *I. uriae* (Pettersson et al., 2020) | Antarctica, Australia (Pettersson et al., 2020) |
| Kadam virus | *Kadam virus* | *Flavivirus* | *Flaviviridae* | ssRNA(+) | NC_033724 | Complete genome | ? | 1970 (Henderson et al., 1970) | *R. pravus* (Shi et al., 2018) | Saudi Arabia, Uganda (Shi et al., 2018) |
| Alkhumra hemorrhagic fever virus | *Kyasanur Forest disease virus* | *Flavivirus* | *Flaviviridae* | ssRNA(+) | AF331718 | Complete genome | Y (Grard et al., 2007) | 1957 (Work et al., 1957) | *H. spinigera, O. savignyi* (Shi et al., 2018) | India, Saudi Arabia (Shi et al., 2018) |
| Langat virus | *Langat virus* | *Flavivirus* | *Flaviviridae* | ssRNA(+) | AF253419 | Complete genome | ? | 1956 (Smith et al., 1956) | *I. granulatus* (Shi et al., 2018) | Malaysia (Shi et al., 2018) |
| Louping ill virus | *Louping ill virus* | *Flavivirus* | *Flaviviridae* | ssRNA(+) | Y07863 | Complete genome | Y (Davidson et al., 1991) | 1931 (Greig et al., 1931) | *I. ricinus* (Shi et al., 2018) | England, Ireland, Scotland, Wales (Shi et al., 2018) |
| Meaban virus | *Meaban virus* | *Flavivirus* | *Flaviviridae* | ssRNA(+) | NC_033721 | Complete genome | ? | 1985 (Chastel et al., 1985) | *O. maritimus* (Shi et al., 2018) | France (Shi et al., 2018) |
| Omsk hemorrhagic fever virus | *Omsk hemorrhagic fever virus* | *Flavivirus* | *Flaviviridae* | ssRNA(+) | AY193805 | Complete genome | Y (Grard et al., 2007) | 1989 (Gaidamovich et al., 1989) | *Dermacentor spp., I. persculatus* (Ruzek et al., 2010) | Russia (Grard et al., 2007) |
| Deer tick virus | *Powassan virus* | *Flavivirus* | *Flaviviridae* | ssRNA(+) | AF311056 | Complete genome | ? | 1997 (Telford et al., 1997) | *I. scapularis (Mansfield et al., 2017; Shi et al., 2018)* | New England, North America (Mansfield et al., 2017; Shi et al., 2018) |
| Powassan virus | *Powassan virus* | *Flavivirus* | *Flaviviridae* | ssRNA(+) | L06436 | Complete genome | Y (Tokarz et al., 2014) | 1959 (McLean and Donohue 1959) | *I. cookei, Ixodes spp. (Tokarz et al., 2014)* | North America, Russia (Mansfield et al., 2017) |
| Karshi virus | *Royal Farm virus* | *Flavivirus* | *Flaviviridae* | ssRNA(+) | AY863002 | Complete genome | ? | 1976 (L'vov et al., 1976) | *A. hermanni, O. papillipes* (Shi et al., 2018) | North of Central Asia, Uzbek (Shi et al., 2018) |
| Saumarez Reef virus | *Saumarez Reef virus* | *Flavivirus* | *Flaviviridae* | ssRNA(+) | DQ235150 | Complete genome | ? | 1977 (St George et al., 1977) | *I. eudyptidis, O. capensis* (Shi et al., 2018) | Australia (Shi et al., 2018) |
| Tick-borne encephalitis virus | *Tick-borne encephalitis virus* | *Flavivirus* | *Flaviviridae* | ssRNA(+) | NC_001672 | Complete genome | Y (Shi et al., 2018) | 1931 (Schneider et al., 1931) | *I. ovatus, I. persulcatus, I. ricinus* (Shi et al., 2018) | Northern Asia, Northern Europe, Siberia (Shi et al., 2018) |
| Tyuleniy virus | *Tyuleniy virus* | *Flavivirus* | *Flaviviridae* | ssRNA(+) | NC_023424 | Complete genome | ? | 1971 (L'vov et al., 1971) | *I. putus* (Shi et al., 2018) | Tuleniy Island (Shi et al., 2018) |
| Ixodes holocyclus iflavirus | *Ixodes holocyclus iflavirus* | *Iflavirus* | *Iflaviridae* | ssRNA(+) | OL452115 | Partial genome | ? | 2018 (O'Brien et al., 2018) | *I. holocyclus* (O'Brien et al., 2018) | Australia (O'Brien et al., 2018) |
| Blacklegged tick associated ilarvirus | Unclassified | *Ilarvirus* | *Bromoviridae* | ssRNA(+) | MG647776, MG647777 | Partial genome | ? | 2018 (Tokarz et al., 2018) | *I. scapularis (Tokarz et al., 2018)* | USA (Tokarz et al., 2018) |
| Trinbago virus | Unclassified | *Pestivirus* | *Flaviviridae* | ssRNA(+) | MN025505 | Partial genome | ? | 2019 (Sameroff et al., 2019) | *A. ovale, R. microplus, R. sanguineus* (Sameroff et al., 2019) | Trinidad, Tobago (Sameroff et al., 2019) |
| Cattle tick tymovirus-like virus 1 | Unclassified | *Tymovirus* | *Tymoviridae* | ssRNA(+) | MN025504 | Partial genome | ? | 2019 (Sameroff et al., 2019) | *R. microplus* (Sameroff et al., 2019) | Trinidad and Tobago (Sameroff et al., 2019) |
| Lone star tick dicistrovirus | Unclassified | Unclassified | *Dicistroviridae* | ssRNA(+) | KX774633 | Partial genome | ? | 2018 (Tokarz et al., 2018) | *I. scapularis (Tokarz et al., 2018)* | USA (Tokarz et al., 2018) |
| Alongshan virus | Unclassified | Unclassified | *Flaviviridae* | ssRNA(+) | MN648770, MN648771, MN648772, MN648773 | Partial genome | Y (Wang et al., 2019) | 2019 (Kuivanen et al., 2019) | *I. persulcatus, I. ricinus* (Kholodilov et al., 2020) | China, France, Finland (Kholodilov et al., 2020) |
| Jingmen tick virus | Unclassified | Unclassified | *Flaviviridae* | ssRNA(+) | MK721629, MK721630, MK721631, MK721632 | Complete genome | Y (Jia et al., 2019) | 2014 (Qin et al., 2014) | *Am. javanense, D. nuttalli, Haemaphysalis spp., Ixodes spp.,* *Rhipicephalus* spp. (Zhao et al., 2021) | Many countries in Asia, America, and Africa (Vandegrift et al., 2020; Zhao et al., 2021) |
| Norway luteo-like virus 1 | Unclassified | Unclassified | *Luteoviridae* | ssRNA(+) | MF141065 | Partial genome | ? | 2017 (Pettersson et al., 2017) | *I. ricinus* (Pettersson et al., 2017) | Norway (Pettersson et al., 2017) |
| Norway luteo-like virus 2 | Unclassified | Unclassified | *Luteoviridae* | ssRNA(+) | MF141068 | Partial genome | ? | 2017 (Pettersson et al., 2017) | *I. ricinus* (Pettersson et al., 2017) | Norway (Pettersson et al., 2017) |
| Norway luteo-like virus 3 | Unclassified | Unclassified | *Luteoviridae* | ssRNA(+) | MF141069 | Partial genome | ? | 2017 (Pettersson et al., 2017) | *I. ricinus* (Pettersson et al., 2017) | Norway (Pettersson et al., 2017) |
| Norway luteo-like virus 4 | Unclassified | Unclassified | *Luteoviridae* | ssRNA(+) | MF141070 | Partial genome | ? | 2017 (Pettersson et al., 2017) | *I. ricinus* (Pettersson et al., 2017) | Norway (Pettersson et al., 2017) |
| Lone star tick nodavirus | Unclassified | Unclassified | *Nodaviridae* | ssRNA(+) | KX774635, KX774634 | Partial genome | ? | 2018 (Tokarz et al., 2018) | *I. scapularis (Tokarz et al., 2018)* | USA (Tokarz et al., 2018) |
| Paradise bay virus | Unclassified | Unclassified | *Tombusviridae* | ssRNA(+) | MN830247 | Partial genome | ? | 2020 (Pettersson et al., 2020) | *I. uriae* (Pettersson et al., 2020) | Antarctica (Pettersson et al., 2020) |
| Upmeje virus | Unclassified | Unclassified | *Tombusviridae* | ssRNA(+) | MN830246 | Partial genome | ? | 2020 (Pettersson et al., 2020) | *I. uriae* (Pettersson et al., 2020) | Northern Sweden (Pettersson et al., 2020) |
| Bagotville virus | Unclassified | Unclassified | *Unclassified Picornavirales* | ssRNA(+) | MW741892 | Partial genome | ? | 2021 (Chandra et al., 2021) | *I. holocyclus* (Chandra et al., 2021) | Australia (Chandra et al., 2021) |
| Blacklegged tick picorna-like virus 1 | Unclassified | Unclassified | *Unclassified Picornavirales* | ssRNA(+) | MG647769, MG647774 | Partial genome | ? | 2018 (Tokarz et al., 2018) | *I. scapularis (Tokarz et al., 2018)* | USA (Tokarz et al., 2018) |
| Blacklegged tick picorna-like virus 2 | Unclassified | Unclassified | *Unclassified Picornavirales* | ssRNA(+) | MG647773, MG647772, MG647771, MG647770 | Partial genome | ? | 2018 (Tokarz et al., 2018) | *I. scapularis (Tokarz et al., 2018)* | USA (Tokarz et al., 2018) |
| Wardell virus | Unclassified | Unclassified | *Unclassified Picornavirales* | ssRNA(+) | MW741896, MW741895, MW741894 | Partial genome | ? | 2021 (Chandra et al., 2021) | *I. holocyclus* (Chandra et al., 2021) | Australia (Chandra et al., 2021) |
| Woodburn virus | Unclassified | Unclassified | *Unclassified Picornavirales* | ssRNA(+) | MW741893 | Partial genome | ? | 2021 (Chandra et al., 2021) | *I. holocyclus* (Chandra et al., 2021) | Australia (Chandra et al., 2021) |
| Blue Fish Point virus | Unclassified | Unclassified | *Unclassified Riboviria* | ssRNA(+) | MK026570, MK026571 | Partial genome | ? | 2018 (Harvey et al., 2018) | *I. holocyclus (Harvey et al., 2018)* | Australia (Harvey et al., 2018) |
| Bole tick virus 4 | Unclassified | Unclassified | *Unclassified Riboviria* | ssRNA(+) | MZ244286 | Complete genome | ? | 2015 (Shi et al., 2015) | *D. marginatus, D. reticulatus, H. asiaticum, Hy. asiaticum, Hy. detritum, R. microplus, R. sanguineus (Temmam et al., 2019; Bratuleanu et al., 2021; Zakham et al., 2021)* | China, Romania, Saudi Arabia, Thailand, Trinidad, Tobago (Temmam et al., 2019; Bratuleanu et al., 2021; Zakham et al., 2021) |
| Bulatov virus | Unclassified | Unclassified | *Unclassified Riboviria* | ssRNA(+) | MN830234, MT025173 | Partial genome | ? | 2020 (Pettersson et al., 2020; Wille et al., 2020) | *I. uriae* (Pettersson et al., 2020) | Antarctica (Pettersson et al., 2020) |
| Collins beach virus | Unclassified | Unclassified | *Unclassified Riboviria* | ssRNA(+) | MK026585 | Partial genome | ? | 2018 (Harvey et al., 2018) | *I. holocyclus (Harvey et al., 2018)* | Australia (Harvey et al., 2018) |
| Fairfax Lookout virus | Unclassified | Unclassified | *Unclassified Riboviria* | ssRNA(+) | MK026588, MK026587, MK026586 | Partial genome | ? | 2018 (Harvey et al., 2018) | *I. holocyclus (Harvey et al., 2018)* | Australia (Harvey et al., 2018) |
| Gerbovich virus | Unclassified | Unclassified | *Unclassified Riboviria* | ssRNA(+) | MN830239, MT025175 | Partial genome | ? | 2020 (Pettersson et al., 2020; Wille et al., 2020) | *I. uriae* (Pettersson et al., 2020; Wille et al., 2020) | Antarctica (Pettersson et al., 2020; Wille et al., 2020) |
| Ingleside virus | Unclassified | Unclassified | *Unclassified Riboviria* | ssRNA(+) | OL452105 | Partial genome | ? | 2018 (Harvey et al., 2018) | *I. holocyclus (Harvey et al., 2018)* | Australia (Harvey et al., 2018) |
| Ixodes scapularis associated virus 3 | Unclassified | Unclassified | *Unclassified Riboviria* | ssRNA(+) | LC094965, MG677814 | Partial genome | ? | 2017 (Nakao et al., 2017) | *I. scapularis (Tokarz et al., 2018; Nakao et al., 2017)* | USA, Japan (Tokarz et al., 2018; Nakao et al., 2017) |
| Jump Rock virus | Unclassified | Unclassified | *Unclassified Riboviria* | ssRNA(+) | MK026583 | Partial genome | ? | 2018 (Harvey et al., 2018) | *I. holocyclus (Harvey et al., 2018)* | Australia (Harvey et al., 2018) |
| Nadgee virus | Unclassified | Unclassified | *Unclassified Riboviria* | ssRNA(+) | MK026592 | Partial genome | ? | 2018 (Harvey et al., 2018) | *I. holocyclus (Harvey et al., 2018)* | Australia (Harvey et al., 2018) |
| Store beach virus | Unclassified | Unclassified | *Unclassified Riboviria* | ssRNA(+) | MK026567 | Partial genome | ? | 2018 (Harvey et al., 2018) | *Am. moreliae (Harvey et al., 2018)* | Australia (Harvey et al., 2018) |
| Tacheng tick virus 8 | Unclassified | Unclassified | *Unclassified Riboviria* | ssRNA(+) | NC_028367 | Complete genome | ? | 2015 (Shi et al., 2015) | *D. marginatus (Shi et al., 2015)* | China (Shi et al., 2015) |
| Vovk virus | Unclassified | Unclassified | *Unclassified Riboviria* | ssRNA(+) | MN830235, MT025162 | Partial genome | ? | 2020 (Pettersson et al., 2020; Wille et al., 2020) | *I. uriae* (Pettersson et al., 2020; Wille et al., 2020) | Antarctica (Pettersson et al., 2020; Wille et al., 2020) |
| Wangarabell virus | Unclassified | Unclassified | *Unclassified Riboviria* | ssRNA(+) | MK026593 | Partial genome | ? | 2018 (Harvey et al., 2018) | *I. holocyclus (Harvey et al., 2018)* | Australia (Harvey et al., 2018) |
| Yambulla virus | Unclassified | Unclassified | *Unclassified Riboviria* | ssRNA(+) | MK026594 | Partial genome | ? | 2018 (Harvey et al., 2018) | *I. holocyclus (Harvey et al., 2018)* | Australia (Harvey et al., 2018) |
| Antioquia tymovirus-like 1 | Unclassified | Unclassified | *Unclassified Tymovirales* | ssRNA(+) | MK683451 | Partial genome | ? | 2020 (Gomez et al., 2020) | *R. microplus* (Gomez et al., 2020) | Colombia (Gomez et al., 2020) |
| Guarapuava tymovirus-like 1 | Unclassified | Unclassified | *Unclassified Tymovirales* | ssRNA(+) | MH155882 | Partial genome | ? | 2018 (de Souza et al., 2018) | *R. microplus* (de Souza et al., 2018) | Brazil (de Souza et al., 2018) |
| American dog tick associated virus 1 | Unclassified | Unclassified | Unclassified | Unknown | MF962660 | Partial genome | ? | 2018 (Tokarz et al., 2018) | *D. variabilis (Tokarz et al., 2018)* | USA (Tokarz et al., 2018) |
| American dog tick associated virus 2 | Unclassified | Unclassified | Unclassified | Unknown | MF962662, MF962657 | Partial genome | ? | 2018 (Tokarz et al., 2018) | *D. variabilis (Tokarz et al., 2018)* | USA (Tokarz et al., 2018) |
| Finch Creek virus | Unclassified | Unclassified | Unclassified | Unknown | EU267169 | Partial genome | ? | 2009 (Major et al., 2009) | *I. uriae* (Major et al., 2009) | Australia (Major et al., 2009) |
| Ganjam virus | Unclassified | Unclassified | Unclassified | Unknown | KU925466, KU925465, KU925464 | Partial genome | ? | 1969 (Dandawate et al., 1969) | *H. intermedia* (Kuhn et al., 2016) | India (Kuhn et al., 2016) |
| Ixodes scapularis associated virus 1 | Unclassified | Unclassified | Unclassified | Unknown | LC094964, KM048318 | Partial genome | ? | 2014 (Tokarz et al., 2014) | *I. scapularis (Tokarz et al., 2014; Nakao et al., 2017)* | Japan, USA (Tokarz et al., 2014; Nakao et al., 2017) |
| Ixodes scapularis associated virus 2 | Unclassified | Unclassified | Unclassified | Unknown | KM048319 | Partial genome | ? | 2014 (Tokarz et al., 2014) | *I. scapularis (Tokarz et al., 2014)* | USA (Tokarz et al., 2014) |
| Ixodes scapularis associated virus 4 | Unclassified | Unclassified | Unclassified | Unknown | MF962656 | Partial genome | ? | 2018 (Tokarz et al., 2018) | *I. scapularis (Tokarz et al., 2018)* | USA (Tokarz et al., 2018) |
| Ixodes scapularis associated virus 5 | Unclassified | Unclassified | Unclassified | Unknown | MG256513 | Partial genome | ? | 2018 (Tokarz et al., 2018) | *I. scapularis (Tokarz et al., 2018)* | USA (Tokarz et al., 2018) |
| Ixodes scapularis associated virus 6 | Unclassified | Unclassified | Unclassified | Unknown | MG256514 | Partial genome | ? | 2018 (Tokarz et al., 2018) | *I. scapularis (Tokarz et al., 2018)* | USA (Tokarz et al., 2018) |
| Lone star tick associated virus 1 | Unclassified | Unclassified | Unclassified | Unknown | MF962658 | Partial genome | ? | 2018 (Tokarz et al., 2018) | *I. scapularis (Tokarz et al., 2018)* | USA (Tokarz et al., 2018) |
| Sapphire II virus | Unclassified | Unclassified | Unclassified | Unknown | MK896468, MK896467, MK896466 | Complete genome | ? | 1972 (Yunker et al., 1972) | *A. cooleyi* (Yunker et al., 1972) | USA (Yunker et al., 1972) |
| Tick borne tetravirus-like virus | Unclassified | Unclassified | Unclassified | Unknown | MW334982, KM048322 | Partial genome | ? | 2014 (Tokarz et al., 2014) | *D. variabilis (Tokarz et al., 2014)* | USA (Tokarz et al., 2014) |
| Zirqa virus | Unclassified | Unclassified | Unclassified | Unknown | KU925500, KU925501, KU925502 | Complete genome | ? | 1973 (Varma et al., 1973) | *A. cooleyi, Ornithodorus* spp. (Shi et al., 2018) | Abu Dhabi (Shi et al., 2018) |
| Bovine hepacivirus** | *Hepacivirus N* | *Hepacivirus* | *Flaviviridae* | ssRNA(+) | NC_026797 | Complete genome | ? | 2015 (Baechlein et al., 2015) | Unknown (Temmam et al., 201) | Brazil, China, Germany, Ghana, US (Baechlein et al., 2019) |
| HoBi-like pestivirus** | *Pestivirus H* | *Pestivirus* | *Flaviviridae* | ssRNA(+) | MK689376 | Partial genome | ? | 2004 (Schirrmeier et al., 2004) | Unknown (Temmam et al., 201) | Argentina, Bangladesh, Brazil, China, Germany, India, Italy, Mexico, Thailand, Turkey, USA (Bauermann et al., 2021) |
| Erve virus** | *Erve orthonairovirus* | *Orthonairovirus* | *Nairoviridae* | ssRNA(-) | JF911697, JF911698, JF911699 | Complete genome | ? | 1989 (Chastel et al., 1989) | Unknown (Dilcher et al., 2012) | Czech Republic, France, Germany, Netherland (Dilcher et al., 2012) |
| Bovine parvovirus-2** | *Ungulate copiparvovirus 1* | *Copiparvovirus* | *Parvoviridae* | ssDNA(+/-) | KT148961 | Complete genome | ? | 2001 (Allander et al., 2001) | Unknown (Temmam et al., 201) | Brazil, China, USA (Ng et al., 2015; Cibulski et al., 2016; Wang et al., 2018) |
| Nkolbisson virus** | *Nkolbisson ledantevirus* | *Ledantevirus* | *Rhabdoviridae* | ssRNA(-) | NC_034539 | Complete genome | ? | 2010 (Dacheux et al., 2010) | Unknown (Temmam et al., 2019) | Cameroon (Dacheux et al., 2010) |
| Spring viremia of carp virus** | *Carp sprivivirus* | *Sprivivirus* | *Rhabdoviridae* | ssRNA(-) | NC_002803 | Complete genome | ? | 1996 (Bjorklund et al., 1996) | Unknown (Temmam et al., 201) | Austria, China, Czech Republic, Denmark, Hungary, Iran, Italy, Mexico, Moldova, Poland, Serbia, South Korea, Ukraine, UK, USA (Marsella et al., 2021) |
| Maraba virus** | *Maraba virus* | *Vesiculovirus* | *Rhabdoviridae* | ssRNA(-) | NC_025255 | Complete genome | ? | 2010 (Brun et al., 2010) | Unknown (Temmam et al., 201) | Brazil (Brun et al., 2010) |
| Wellfleet Bay virus** | Unclassified | *Quaranjavirus* | *Orthomyxoviridae* | ssRNA(-) | NC_025793, NC_025794, NC_025795, NC_025796, NC_025797, NC_025798, NC_025799 | Complete genome | ? | 2015 (Allison et al., 2015) | Unknown (Temmam et al., 201) | USA (Allison et al., 2015) |
| Araguari virus** | Unclassified | *Thogotovirus* | *Orthomyxoviridae* | ssRNA(-) | KX670389, KX670390, KX670391, KX670392, KX670393, KX670394, KX670395 | Complete genome | ? | 1969 (Silva et al., 2005) | Unknown tick vectors (Silva et al., 2005) | Brazil (Silva et al., 2005) |
| Eelpout rhabdovirus** | Unclassified | Unclassified | *Rhabdoviridae* | ssRNA(-) | KR612230 | Partial genome | ? | 2017 (Axen et al., 2017) | Unknown (Temmam et al., 201) | Sweden (Axen et al., 2017) |
| Wuhan redfin culter dimarhabodovirus** | Unclassified | Unclassified | *Rhabdoviridae* | ssRNA(-) | MG600013 | Complete genome | ? | 2018 (Shi et al., 2018) | Unknown (Temmam et al., 201) | China (Shi et al., 2018) |
| Ahun nairovirus** | Unclassified | Unclassified | Unclassified | Unknown | Not available | Not available | ? | 2014 (Dacheux et al., 2014) | Unknown (Dacheux et al., 2014) | France (Dacheux et al., 2014) |
| Essaouira virus** | Unclassified | Unclassified | Unclassified | Unknown | Not available | Not available | ? | 1993 (Chastel et al., 1993) | *O. maritimus* (Chastel et al., 1993) | Morocco (Chastel et al., 1993) |
| Huacho virus** | Unclassified | Unclassified | Unclassified | Unknown | Not available | Not available | ? | 1967 (Belaganahalli et al., 2015) | *C. amblus* (Belaganahalli et al., 2015) | Peru (Belaganahalli et al., 2015) |
| Kala Iris virus** | Unclassified | Unclassified | Unclassified | Unknown | Not available | Not available | ? | 1993 (Chastel et al., 1993) | *O. maritimus* (Chastel et al., 1993) | Morocco (Chastel et al., 1993) |
| Mono Lake virus** | Unclassified | Unclassified | Unclassified | Unknown | Not available | Not available | ? | 1988 (Calisher et al., 1988) | *A. cooleyi* (Calisher et al., 1988) | USA (Calisher et al., 1988) |
| Sixgun city virus** | Unclassified | Unclassified | Unclassified | Unknown | Not available | Not available | ? | 1988 (Calisher et al., 1988) | *A. cooleyi* (Calisher et al., 1988) | USA (Calisher et al., 1988) |
| Ellidaey virus** | Unclassified | Unclassified | Unclassified | Unknown | Not available | Not available | ? | 1986 (Moss et al., 1986) | *I. uriae* (Moss et al., 1986) | Iceland (Elliđaey island) (Moss et al., 1986) |
| Foula virus** | Unclassified | Unclassified | Unclassified | Unknown | Not available | Not available | ? | 1986 (Nuttall et al., 1986) | *I. uriae* (Nuttall et al., 1986) | Scotland (Shetland Islands) (Nuttall et al., 1986) |
| Grimsey virus** | Unclassified | Unclassified | Unclassified | Unknown | Not available | Not available | ? | 1986 (Moss et al., 1986) | *I. uriae* (Moss et al., 1986) | Iceland (Grimsey island) (Moss et al., 1986) |
| Inner Farne Island virus** | Unclassified | Unclassified | Unclassified | Unknown | Not available | Not available | ? | 1986 (Nuttall et al., 1986) | *I. uriae* (Nuttall et al., 1986) | England (Nuttall et al., 1986) |
| Mykines virus** | Unclassified | Unclassified | Unclassified | Unknown | Not available | Not available | ? | 1978 (Main et al., 1978) | *I. uriae* (Main et al., 1978) | Denmark (Faroe Islands) (Main et al., 1978) |
| Kao Shuan virus** | Unclassified | Unclassified | Unclassified | Unknown | Unavailable | Not available | ? | 1989 (Moussa et al., 1989) | *A. robertsi* (Moussa et al., 1989) | Indonesia, Taiwan (Moussa et al., 1989) |
| Pretoria virus** | Unclassified | Unclassified | Unclassified | Unknown | Not available | Not available | ? | 1975 (Converse et al., 1975) | *A. africolumbae* (Converse et al., 1975) | South Africa (Converse et al., 1975) |
| Puffin Island virus** | Unclassified | Unclassified | Unclassified | Unknown | Not available | Not available | ? | 1983 (Gould et al., 1983) | *O. maritimus* (Gould et al., 1983) | Puffin Island (Gould et al., 1983) |
| Seletar virus** | Unclassified | Unclassified | Unclassified | Unknown | Not available | Not available | ? | 1961 (Shi et al., 2018) | *B. microplus* (Shi et al., 2018) | Malaysia, Singapore (Shi et al., 2018) |

* Amblyomma dissimile mivirus, Nuomin virus, and Granville quaranjavirus were later added tick-borne viruses which were not included in the original tick virus database.

** Viruses with unknown tick vectors or that sequences unavailable.

###### Acronyms: *Argas* (*A.*); *Amblyomma* (*Am.*); *Boophilus* (*B.*); *Dermacentor* (*D.*); *Haemaphysalis* (*H.*); *Hyalomma* (*Hy.*); *Ixodes* (*I.*); *Ornithodoros* (*O.*); *Otobius* (*Ot.*); *Rhipicephalus* (*R.*).

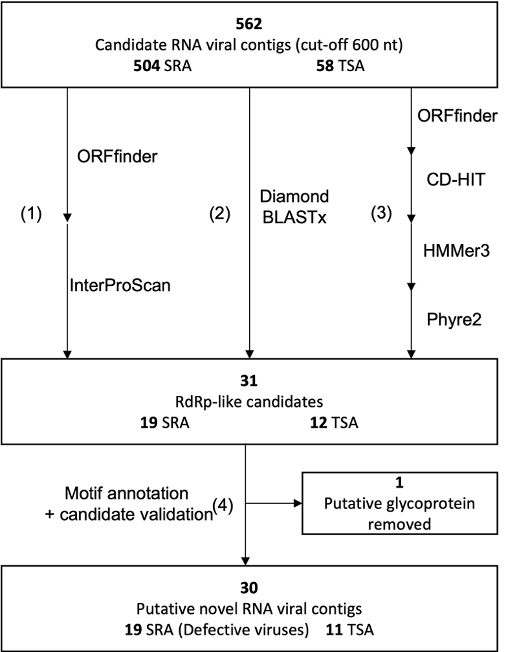

### **Figure S1.** Workflow of detecting viral RNA-dependent RNA polymerase sequences. (1) ORFs were translated using ORFfinder (cut-off 200 aa). The translated sequences were predicted using the InterProScan-5.55-88.0 (--minsize 600; -goterms). (2) BLASTx using Diamond v2.0.14.152 against both non-redundant protein database and RdRp-scan database (-e 1e-5 --min-orf 600 --very-sensitive). (3) ORFs were translated using ORFfinder (cut-off 200 aa). Redundant sequences were then removed using CD-HIT v4.6.8 at 90% level of identity (-c 0.98 sequence identity threshold). Remaining sequences were scanned using HMMer3 v3.2.1 against RdRp HMM profile database (full sequence e-value 1e-06). HMM-RdRp hits were compared against PDB using Phyre2 server. In total, 31 RdRp-like candidates from SRA and TSA databases were found using RdRp-scan and InterProScan after removing duplicates. (4) RdRp motifs were mapped to candidate sequences using Geneious v2022.1. RdRp-scan database, RdRp HMM profile database, RdRp motifs database and RdRp key word lists were obtained from Charon et al. (77) (https://github.com/JustineCharon/RdRp-scan).

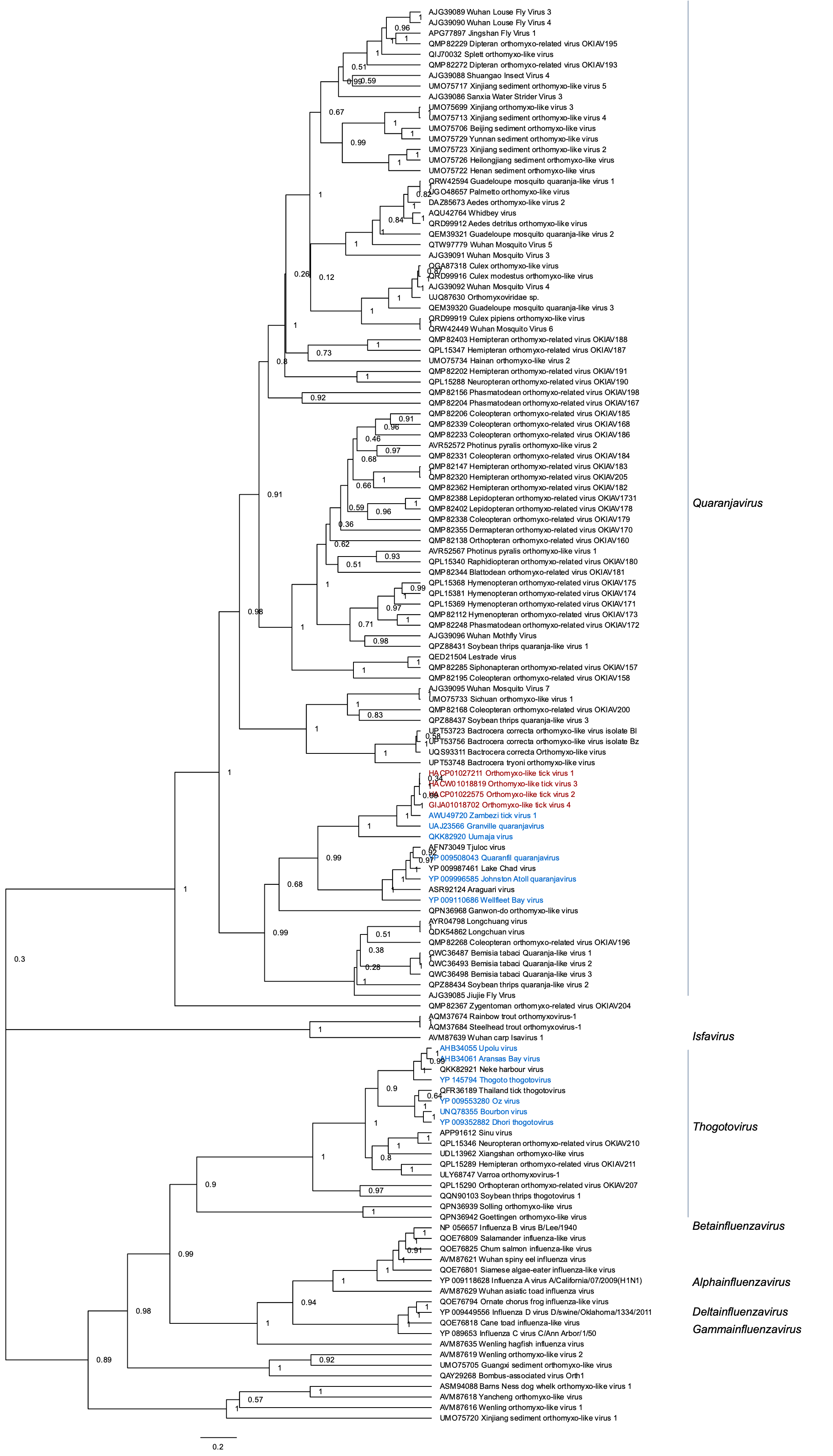

**Figure S2.** Phylogenetic relationship of the novel Orthomyxo-like viral contigs. Maximum clade credibility tree was inferred from the Bayesian posterior sample. Putative novel viral contigs found in the present study are shown in red, and previously known tick-borne viruses are shown in blue.

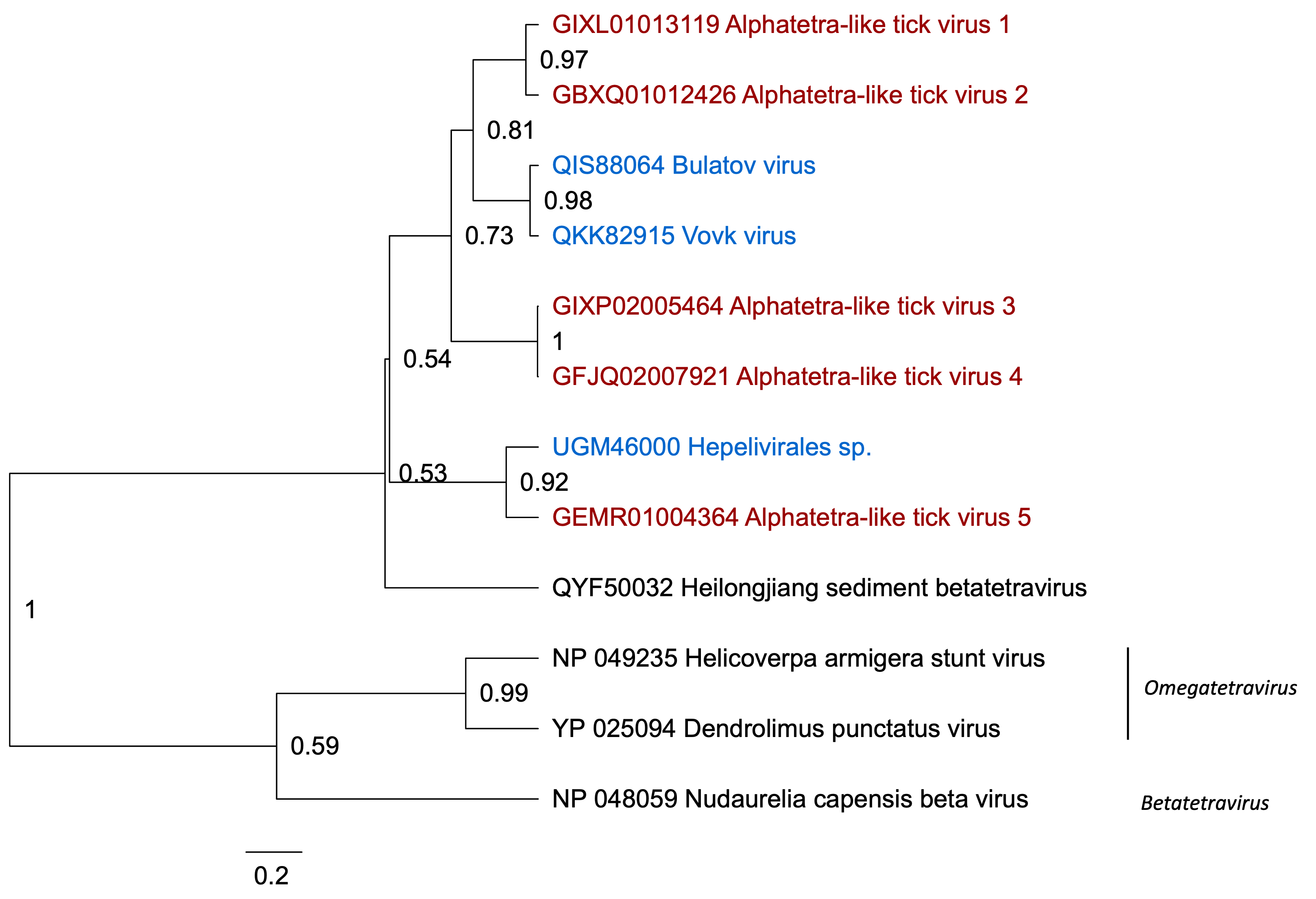

**Figure S3.** Phylogenetic relationship of the novel Alphatetra-like viral contigs. Maximum clade credibility tree was inferred from the Bayesian posterior sample. Putative novel viral contigs found in the present study are shown in red, and previously known tick-borne viruses are shown in blue.

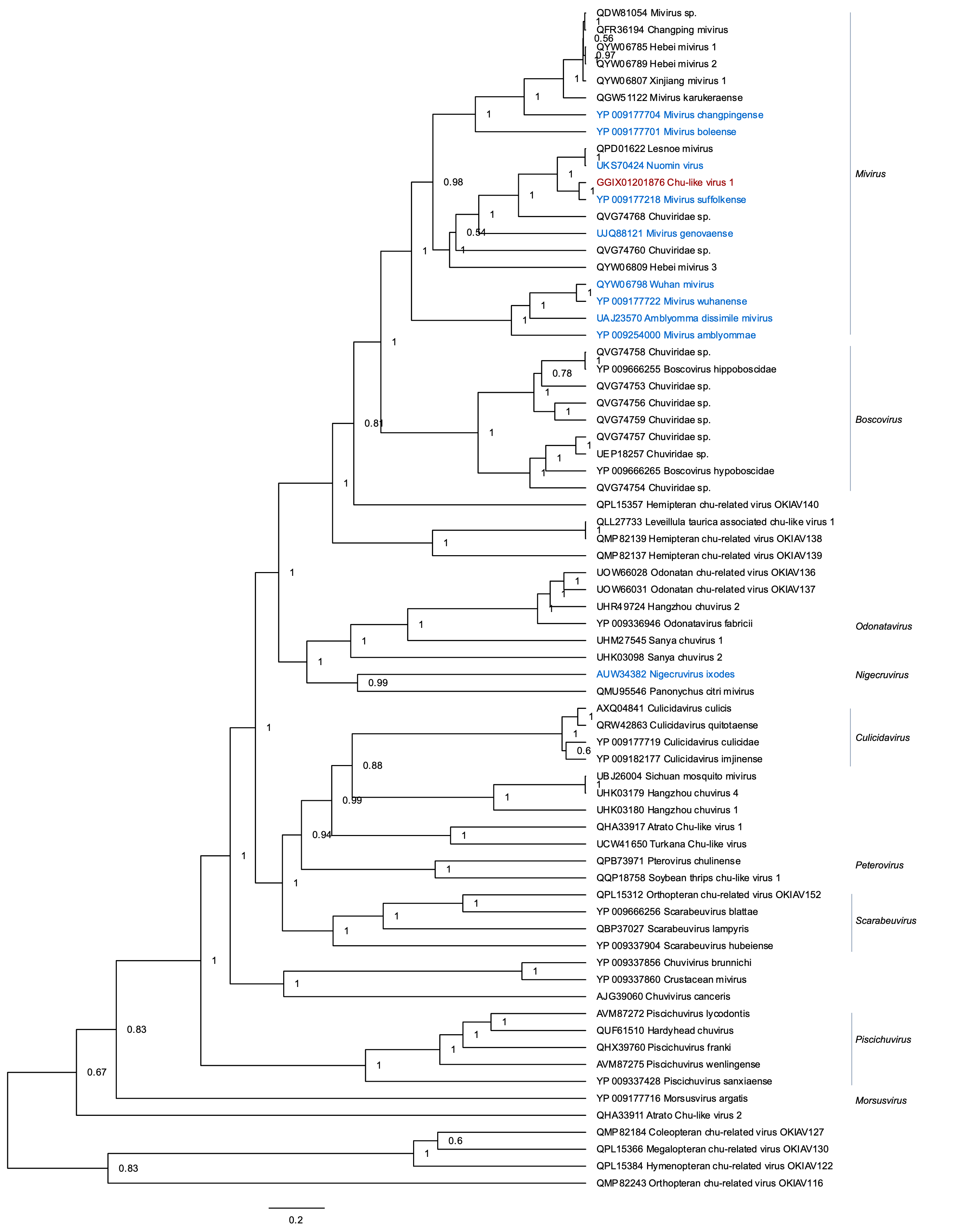

**Figure S4.** Phylogenetic relationship of the novel Chu-like viral contigs. Maximum clade credibility tree was inferred from the Bayesian posterior sample. Putative novel viral contigs found in the present study are shown in red, and previously known tick-borne viruses are shown in blue.
